# GeneMates: an R package for Detecting Horizontal Gene Co-transfer between Bacteria Using Gene-gene Associations Controlled for Population Structure

**DOI:** 10.1101/2020.02.29.970970

**Authors:** Yu Wan, Ryan R. Wick, Justin Zobel, Danielle J. Ingle, Michael Inouye, Kathryn E. Holt

**Affiliations:** Department of Biochemistry and Molecular Biology, Bio21 Molecular Science and Biotechnology Institute, University of Melbourne, Parkville, Victoria 3010, Australia; Department of Infectious Diseases, Central Clinical School, Monash University, Melbourne, Victoria 3004, Australia; School of Computing and Information Systems, University of Melbourne, Parkville, Victoria 3010, Australia; Microbiological Diagnostic Unit Public Health Laboratory, Department of Microbiology and Immunology, University of Melbourne at The Peter Doherty Institute for Infection and Immunity, Parkville, Victoria 3010, Australia; National Centre for Epidemiology and Population Health, Australian National University, Canberra, Australian Capital Territory 2601, Australia; Cambridge Baker Systems Genomics Initiative, Baker Heart and Diabetes Institute, Melbourne, Victoria 3004, Australia; Cambridge Baker Systems Genomics Initiative, Department of Public Health and Primary Care, University of Cambridge, Cambridge, England CB1 8RN, UK; Department of Infection Biology, London School of Hygiene & Tropical Medicine, London, England WC1E 7HT, UK

## Abstract

Antimicrobial resistance (AMR) in bacteria has been a global threat to public health for decades. A well-known driving force for the emergence, evolution and dissemination of genetic AMR determinants in bacterial populations is horizontal gene transfer, which is frequently mediated by mobile genetic elements (MGEs). Some MGEs can capture, maintain, and rearrange multiple AMR genes in a donor bacterium before moving them into recipients, giving rise to a phenomenon called horizontal gene co-transfer (HGcoT). This physical linkage or co-localisation between mobile AMR genes is of particular concern because it facilitates rapid dissemination of multidrug resistance within and across bacterial populations, providing opportunities for co-selection of AMR genes and limiting our therapeutic options. The study of HGcoT can be benefited from large-scale whole-genome sequencing (WGS) data, however, by far most published studies of HGcoT only consider simple co-occurrence measures, which can be confounded by strong bacterial population structure due to clonal reproduction, leading to spurious associations. To address this issue, we present GeneMates, an R package implementing a network approach to identification of HGcoT using WGS data. The package enables users to test for associations between presence-absence of bacterial genes using univariate linear mixed models controlling for population structure based on core-genome variation. Furthermore, when physical distances between genes of interest are measurable in bacterial genomes, users can evaluate distance consistency to further support their inference of putative horizontally co-transferred genes, whose co-occurrence cannot be completely explained by the population structure. We demonstrate how this package can be used to identify co-transferred AMR genes and recover known MGEs from *Escherichia coli* and *Salmonella* Typhimurium WGS data. GeneMates is accessible at github.com/wanyuac/GeneMates.

## 1 Background

Horizontal gene transfer (HGT) accelerates bacterial genome innovation and evolution [1]. It facilitates intra-and interspecies gene flow, dividing each species gene pool (“pan-genome”) into core and accessory genes [2]. Mobile genetic elements (MGEs), such as plasmids, bacteriophages and transposons, are common vectors for HGT [3]. It is well-known that when genes are physically linked (namely, co-localised in an MGE or otherwise physically close in DNA molecules), they can be horizontally co-transferred between bacteria, leading to positive gene-gene associations known as genetic linkage [4, 5].

The rapid accumulation of bacterial whole-genome sequencing (WGS) data in the most recent two decades [6] enables us to study horizontal gene co-transfer (HGcoT) at the population level. Since bacteria reproduce asexually and HGT can occur across different levels of taxonomic boundaries [7], gene-gene associations that cannot be completely explained by bacterial population structure (which determines the distribution of co-inherited genes) suggests HGcoT [8]. Consequently, it is usually trivial to identify candidates of interspecies HGcoT using simple association tests (such as chi-squared tests and simple logistic regression), whereas for detecting intraspecies HGcoT, we must overcome two related challenges arising from the presence of population structure within a species: (1) how to control for population structure in association tests; and (2) how to accurately estimate or represent the population structure to be controlled for.

Univariate linear mixed models (LMMs), which have been widely used in human genome-wide association studies (GWAS) [9] and recently applied to bacterial GWAS [10], provide a solution to address both challenges. Each model explains a response variable using a fixed effect of an independent variable and mixed random effects of population structure and environmental factors. For each LMM, the population structure is represented by a relatedness matrix, whose principal components (PCs) can be used for an orthonormal transformation of genetic variation underlying the population structure [11]. McVean demonstrates that not only do these PCs simplify computations, but also correlate with bacterial genealogies [12]. Compared to phylogeny-based association tests, such as phylogenetically independent contrasts and phylogenetic generalised least squares [13], LMMs do not rely on specific phylogenetic models nor assume that mutation rather than recombination dominates genetic variation.

Here, we introduce an R package GeneMates, which implements a novel network approach to the identification of HGcoT within a bacterial species. This approach takes as input specific information extracted from bacterial WGS data and produces a network showing allele-level evidence of HGcoT. We validated GeneMates using published WGS data from two bacterial species, *Escherichia coli* and *Salmonella enterica*, with which we identify horizontally co-transferred AMR genes. We also provide helper scripts to assist users in preparing necessary input files from standard formats. Since GeneMates is theoretically applicable to any kind of acquired genes in bacteria, it has the potential to also be used for investigating structures and dissemination of other mobile gene clusters of interest, such as virulence genes.

## 2 Implementation

GeneMates consists of R functions performing network construction, topological analysis and data visualisation. It works at the allele level to track recent HGcoT. Particularly, we assume that bacterial isolates under study will typically be collected in a period that is too short to accumulate any mutation in recently acquired genes of interest, such as AMR genes. In addition, for convenience we assume that every gene has zero or one allele in each genome as it is not possible to reliably resolve exact alleles of the same gene using short reads, which by far account for the majority of WGS data.

We developed a network approach to integrate and visualise evidence of physical linkage between alleles of mobile genes in bacteria. In the network, nodes represent alleles and weighted directed edges reflect the strength of evidence. GeneMates produces such a network in each run. Let vector *e* denote an edge, then it represents a linear model *Y* ~ *X*, where scalar variables *Y* and *X* denote presence-absence of alleles *Y* and *X* in an isolate, respectively. The model explains the distribution of allele *Y* (response allele) with that of allele *X* (explanatory allele) and covariates. For GeneMates, the covariates are isolate projections in an orthonormal transformation of the population structure based on a core-genome relatedness matrix. Correspondingly, the edge is directed from node *X* to node *Y*. A user may filter edges of a resulting network based on the association score *s_a_* (*e*) and distance score *s_d_*(*e*) in order to identify edges showing strong evidence of physical linkage, and carry out topological analysis for the filtered network afterwards. In following subsections, we describe key elements of our approach. Additional details of implementation are provided in Section 3 of Additional File 1.

### 2.1 Network construction

Figure 1 illustrates our work flow for network construction – the core functionality of GeneMates. In order to integrate functions implementing this work flow, we created a wrapper function *findPhysLink* (find physical linkage), which takes as input a binary allelic presence-absence matrix (PAM) of target genes across genomes, a matrix of biallelic core-genome SNPs (cgSNPs), and optionally, a table of allelic physical distances (APDs) for target genes. These inputs can be extracted from read alignments and genome assemblies using our helper scripts (released with GeneMates). Function *findPhysLink* determines nodes and edges of a resulting association network and produces tables that can be exported to Cytoscape [14] as node and edge attributes for network visualisation.

**Figure 1:**
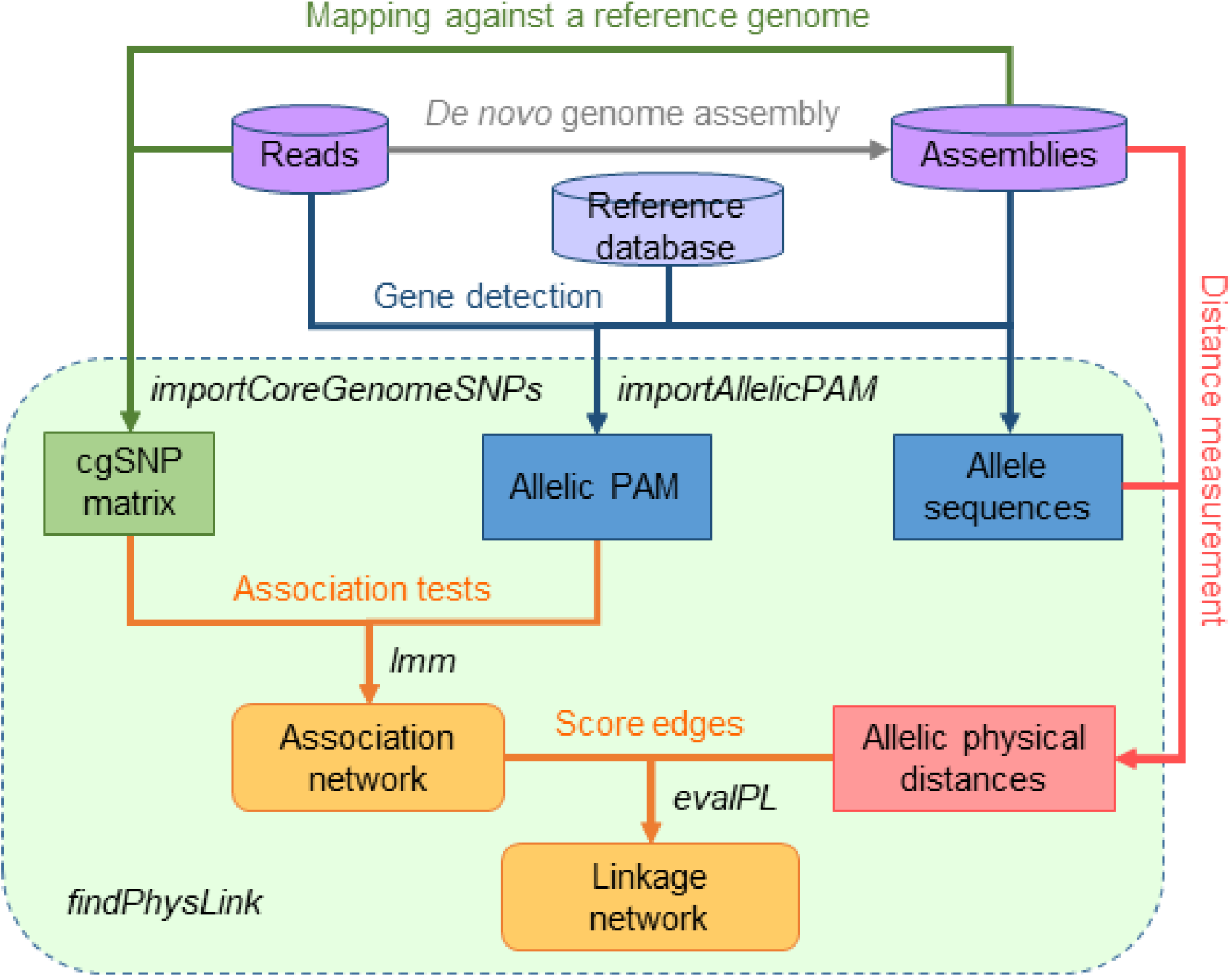
An overall work flow of function *findPhysLink*. In this flowchart, cylinders denote both the input WGS data and a non-redundant (namely, with no sequence duplication) reference database of mobile AMR genes; rounded rectangles denote the two key outputs - an allele network, and a linkage network when reliable APDs are available; ordinary rectangles denote intermediate results, which are matrices or data frames; each arrow represents a process of specific data analysis, which starts from the input and ends at its outcome, with the process name labelled besides the line. Steps integrated with *findPhysLink* are shaded within the dashed rectangular border. Abbreviations: cgSNPs, core-genome SNPs, which are particularly restricted to biallelic SNPs conserved in all isolates for this package; PAM: a binary presence-absence matrix.

#### 2.1.1 Node generation

Assuming *m* alleles of target genes are detected in n bacterial isolates, GeneMates function *importAllelicPAM* imports an *n* × *m* binary PAM ***A*** = (*a_ij_*), where an entry *a_i,j_* = 1 if the *j*-th allele is found in the *i*-th isolate, and *a_i,j_* = 0 otherwise. This function can discard alleles of insufficient frequencies and/or co-occurrence frequencies with two optional filters. In order to reduce the number of tests for allele-allele associations, we followed the implementation of R package BugWAS [10] and coded *importAllelicPAM* to de-duplicate each group of identical columns of PAM into a binary vector called an allelic distribution pattern (Section 3.1.2 of Additional File 1). The function uses column means to apply a column-wise zero-centring to the pattern matrix, which is a common technique used for simplifying matrix algebra without affecting the distribution of data points [15].

#### 2.1.2 Edge weights

GeneMates evaluates two kinds of evidence to assess HGcoT between alleles *X* and *Y*. The first evidence is a significant positive fixed effect of *X* on presence-absence of *Y* when controlling for bacterial population structure. Testing for this effect is an analogue of GWAS, which test for genotype-phenotype associations. Specifically, for edge e in an association network, GeneMates function *lmm* estimates parameters of a univariate LMM using a residual maximum likelihood (REML) approach to test for the fixed effect of *X*, and another function *evalPL* transforms the effect size *β* and its Bonferroni-corrected p-value into an association score *s_a_*(*e*) with possible values 1 (significant positive association), −1 (significant negative association), and 0 (insignificant association). See Section 3.1 in Additional File 1 for details.

The second evidence comes from consistent physical distances between alleles *X* and *Y* in different bacterial genomes (namely, APDs) as structural variation is likely to only occur at a limited level within a mobile gene cluster in a short period. For instance, the same ARG cluster *sul2-strA-strB* has been circulating among Gram-negative bacteria for decades due to its association with plasmids and transposons [16]. Since APDs are measured in genome assemblies, whose completeness determines the amount of measurable APDs when *X* and *Y* are co-localised in the same genomic region, for edge e, we also consider its distance measurability *m_in_*(*e*) - the percentage of genomes in which the APDs between *X* and *Y* are actually measurable. This measurability value is calculated by GeneMates function *summariseDist*, which also evaluates the consistency of APDs included for the distance assessment and assigns a consistency score *c*(*e*) with values −1 (evidence against physical linkage), 0 (insufficient evidence) and 1 (evidence supporting physical linkage). Notably, this function estimates the probability of distance identity-by-descent (IBD) and compares it to a user pre-defined threshold (default: 0.9) for the assignment of each consistency score. See Section 3.3.2 and Figure s18 in Additional File 1 for details. A summary distance score is defined for edge *e* as the consistency score weighted by measurability: *s_d_*(*e*) = *m_in_*(*e*)*c*(*e*). Finally, the association and distance scores are summed to get a linkage score *s*(*e*), reflecting the evidence of physical linkage, where –2 ≤ *s*(*e*) ≤ 2 because 0 ≤ *m_in_*(*e*) ≤ 1. Particularly, we define a linkage network as an association network in which weights of each edge are comprised of a fixed-effect size and a linkage score (Section 3.3 in Additional File 1).

### 2.2 Network visualisation and topological analysis

GeneMates comprises several functions used for displaying resulting networks and exploring network topology for evidence of HGcoT under given conditions. For instance, function *mkNetwork* (make network) extracts user-specified node attributes (such as the frequency and associated AMR phenotype of each allele) and edge attributes (such as the estimated fixed effect size 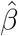 and linkage score *s* per edge) from result tables of *findPhysLink* and print them into plain-text files that can be imported by Cytoscape for network visualisation. Function *extractSubgraphs* follows a predefined node list to pull out subnetworks from a parental network produced by *findPhysLink*. The subnetworks can be maximal cliques identified using the function *max_diques* of an R package igraph [17]. In addition, function *countNeighbours* lists the number of neighbours per node in a network and *getClusterMemberCooccurrence* finds out isolates in which member alleles of each subnetwork are co-occurring.

## 3 Results

We assessed the performance of GeneMates and validated our methodology using published WGS data sets of two bacterial pathogens of great clinical concern – multidrug-resistant *Escherichia coli* and *Salmonella enterica* serovar Typhimurium. Genomes in these well studied data sets have distinct population structures, contain diverse AMR genes and MGEs, and show known gene-gene associations that we expected GeneMates to identify. See Section 4 of Additional File 1 for details of materials and methods.

### 3.1 Characteristics of example data sets

Both the *E. coli* and *Salmonella* data sets consisted of paired-end Illumina short reads, generated from 169 genomes of *E. coli* collected during the Global Enteric Multicentre Study (GEMS) [18, 19] and 359 genomes of typical *S*. Typhimurium Definitive Type 104 (DT104) [20], respectively. Additional File 2 provides detailed genome information.

#### AMR gene content

In the 169 *E. coli* genomes, we identified 178 alleles of 33 AMR genes conferring resistance to eight antimicrobial classes (Figure s1). The four known intrinsic AMR genes of *E. coli* (*ampH, ampC*1, *ampC*2 and *mrdA*) displayed higher frequencies than the 29 acquired AMR genes (> 87% versus < 63%). Altogether, we detected 67 alleles of acquired AMR genes, including 45 alleles showing frequencies less than 3%. In the 359 *Salmonella* genomes, we identified 57 alleles of 24 AMR genes (conferring resistance to six antimicrobial classes), including a single allele of the known intrinsic AMR gene (*aac6-Iaa*) of *S. enterica* and 56 alleles of 23 acquired AMR genes (Figure s3). Notably, five acquired AMR genes (*sul1, aadA*, bla_cARB_, *tet*(G) and *floR*) that are known to be frequently present in the *Salmonella* genomic island 1 (SGI1) [21] were only detected in the dominant lineage of the *Salmonella* genomes (Figure s4).

#### Core-genome SNP sites

We performed SNP analysis with the method described in Section 4.2 of Additional File 1. Particularly, we define cgSNP sites as SNP sites detected in all genomes and outside of repetitive or prophage regions. Altogether, the numbers of cgSNP sites used for correcting for population structure (namely, biallelic cgSNP sites) of the 169 *E. coli* genomes and 359 *Salmonella* genomes were 209,097 and 2,316, respectively. The percentage of total genetic variation captured by PCs of each cgSNP matrix is illustrated in Figure s8a. Furthermore, the REML estimate of parameter λ (which reflects the effect of population structure on the distribution of the response allele) in the null LMM for presence-absence of each allele of AMR genes perfectly predicted whether ≥ 5 PCs had significant effects (Bonferroni-corrected p-values ≤ 0.05) on the distribution of this allele (Figure s8b).

### 3.2 Effects of controlling for population structure

In order to evaluate the effect of controlling for population structure using LMMs when measuring associations between alleles of acquired AMR genes, we compared unadjusted p-values of fixed effects estimated using the LMMs to those estimated using simple penalised logistic models (PLMs) [22] (Figure 2). Particularly, we considered a fixed effect estimated with either kind of models significant if its Bonferroni-corrected p-value was ≤ 0.05. Figure 3 illustrates a directed comparative network for detected alleles of AMR genes in each example data set. In this network, each edge starts from an explanatory allele and terminates at a response allele, representing a significant fixed effect of the explanatory allele in an LMM or PLM or both. For the 67 alleles in *E. coli* genomes, 3,364 LMMs were applied to 60 allele distribution patterns, and 118 significant pairwise associations were detected. Simple PLMs for the same allele patterns of AMR genes in *E. coli* genomes identified 70 significant associations, 50 of which overlapped with those from LMMs. The resulting comparative network consisted of 45 nodes, 138 edges and two connected graph components (Figure 3a). Regarding the 56 alleles in *Salmonella* genomes, 2,040 LMMs were applied to 48 allele distribution patterns and 112 significant pairwise associations were detected. Simple PLMs for the same allele patterns of AMR genes in *Salmonella* genomes identified 48 associations, 36 of which overlapped with those from LMMs. The resulting comparative network consisted of 32 nodes, 124 edges and a single connected graph component (Figure 3b). In addition, for both the *E. coli* and *Salmonella* data sets, estimates of fixed effects in LMMs displayed a complete sign identity to those in PLMs given a maximum type-I error rate 0.05.

**Figure 2:**
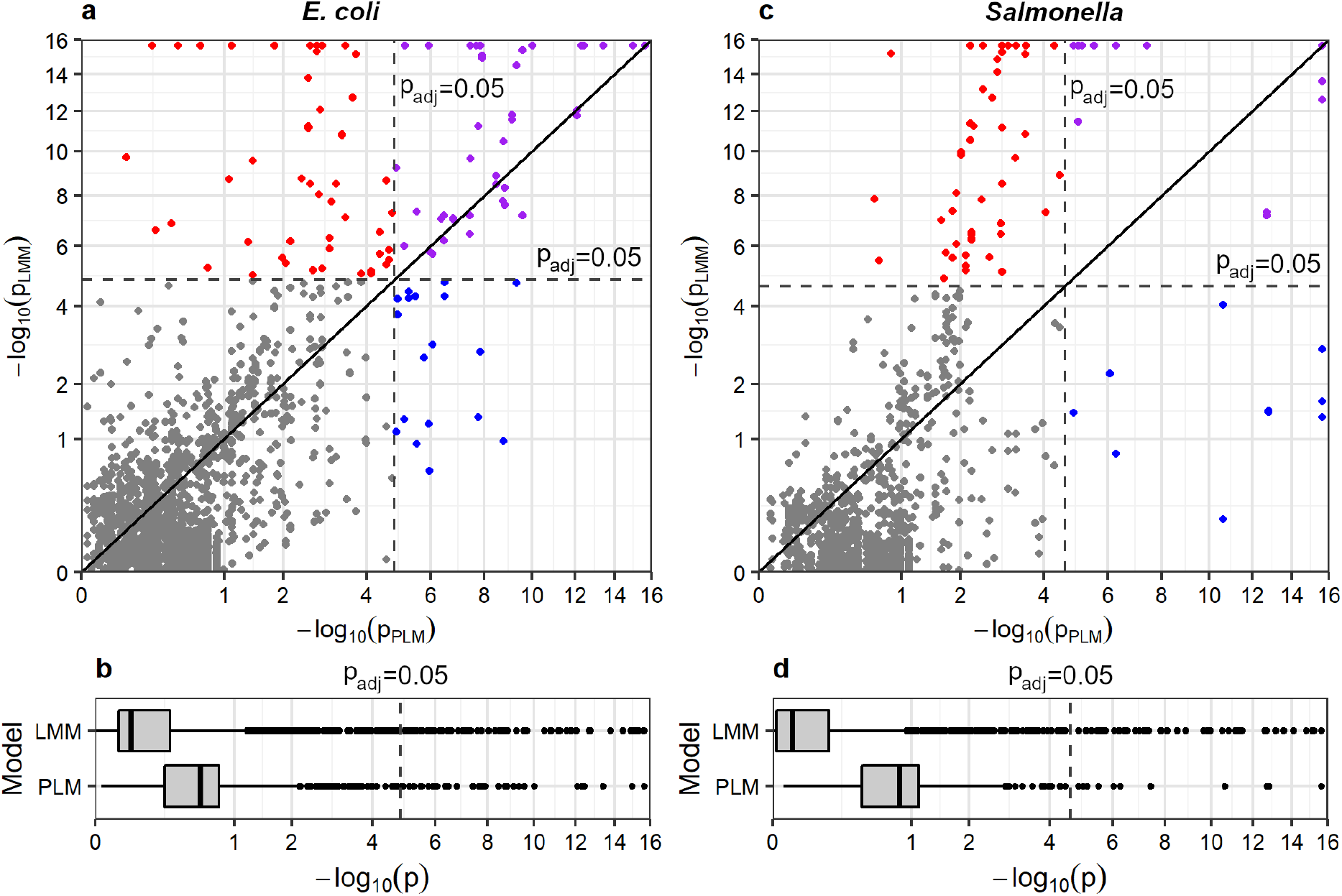
A comparison of unadjusted p-values from PLMs and LMMs for the same pairs of allelic distribution patterns of AMR genes. For the *E. coli* data set (panels **a** and **b**) and *Salmonella* data set (panels **c** and **d**), respectively, a scatter plot and box plot of paired p-values are drawn on square-root transformed axes to compare the p-values. Any p-value less than 2.2 × 10^-16^ is rounded to 2.2 × 10^-16^ owing to the smallest precise floating number in our computer. In panels **a** and **c**, black diagonal lines indicate equality between p-values from these two kinds of models, and grey dashed lines indicate the p-value corresponding to the Bonferroni corrected p-value 0.05, which is used in this study as a cut-off for significant associations. Associations that were only significant in PLMs, only significant in LMMs, significant in both PLMs and LMMs, and significant in neither kind of models, are represented by blue, red, purple, and grey circles, respectively.

**Figure 3:**
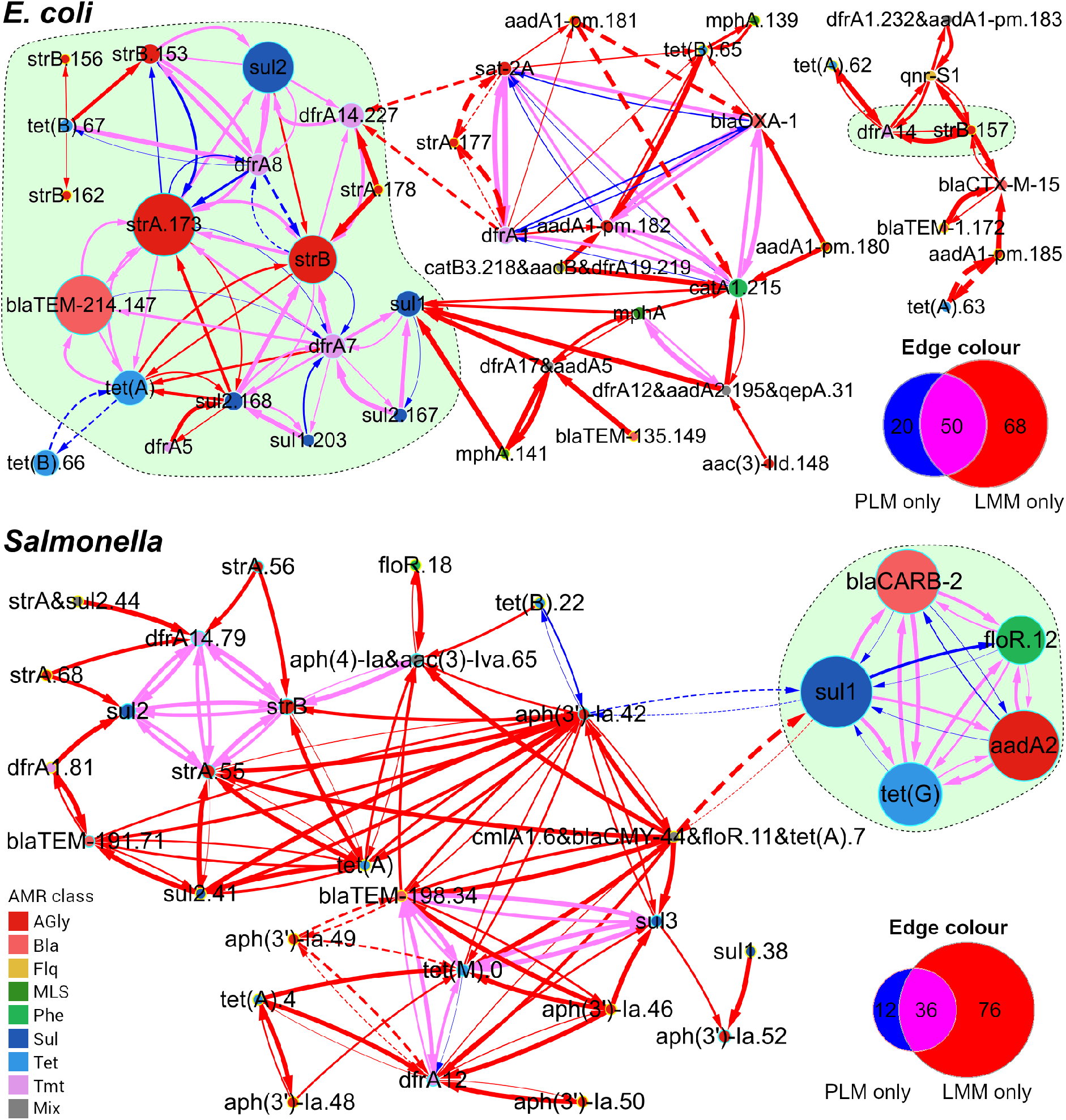
Comparative networks for detected alleles of acquired AMR genes in 169 *E. coli* and 359 *Salmonella* genomes. Each node represents an allele or a cluster of identically distributed alleles, with a diameter proportional to the allele frequency and a fill colour indicating the AMR phenotype encoded. The yellow node border denotes an allele or allele cluster detected in one genome. The edge width is proportional to the strength of a significant association 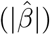 determined using an LMM. Edge colours indicate significant associations from LMMs or PLMs or both. Solid and dashed edges represent significant positive 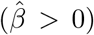 and significant negative associations 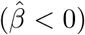, respectively, identified using models indicated by edge colours. The Venn diagram of each panel counts edges under each colour. The shaded area (light green) encircles alleles of known co-mobilised AMR genes. AMR classes defined by antimicrobials that were resistant to: AGly (aminoglycosides), Bla (beta-lactams), Flq (fluoroquinolones), MLS (macrolides, lincosamides, and streptogramins), Phe (phenicols), Sul (sulfonamides), Tet (tetracyclines), Tmt (trimethoprim), and Mix (multiple classes of antimicrobials).

For most allele pairs, correcting for population structure using LMMs yielded higher p-values: the majority of fixed effects (74% for *E. coli* and 85% for *Salmonella*) in LMMs became less significant than those in PLMs (Figure 2b, d). Nonetheless, associations for some allele pairs became more significant after adjusting for population structure, and in most of such cases (93% and 99% for *E. coli* and *Salmonella*, respectively), the distribution of at least one of the alleles showed moderate to strong structural random effects 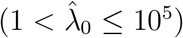. Notably, for both data sets the number of significant associations became greater when adjusting for population structure (Figures 2 and 3), although the increase was not significant (two-sided p-value = 0.58 using a two-proportion Z-test for the null hypothesis of identical proportions).

### 3.3 Effects of adding APDs to association networks

APDs were measured by the shortest-path distance (SPD), defined as the smallest distance between two given loci in an assembly graph (constructed using Unicycler [23]). This approach allows us to recover query sequences that are split into adjacent contigs and to measure APDs between loci that are located in different contigs, potentially increasing distance measurability of the graph, defined as the percentage of measured APDs in all possible APDs of a complete circular genome. Nonetheless, since SPDs are affected by the topology of each assembly graph, which is usually tangled and partially resolved when only short reads are used for genome assembly, it is necessary to determine appropriate criteria for filtering out inaccurate SPDs. Therefore, we downloaded from GenBank [24] 10 complete reference genomes of multidrug resistant *E. coli* and *S*. Typhimurium, respectively (Additional File 2), and compared SPDs measured in *de novo* assembly graphs (constructed from simulated Illumina reads) to true physical distances extracted from the original circularised complete genomes (Supplementary Section 4.6). Specifically, we used Bandage [25] to identify BLAST hits to random coding sequences (CDSs) in each assembly graph and to extract the SPDs for each pair of hits.

#### Filters determined for removing inaccurate SPDs

We considered an SPD accurate if its error fell within a given tolerance range (for instance, ±2 kbp), and hence defined the accuracy rate as the percentage of accurate SPDs in all SPDs, either filtered or not. In practice, any parameter for BLAST hits and SPD measurement can be taken into account for excluding SPDs. In this study, we assessed the accuracy rate of SPDs measured within various maximum distances and node numbers for each assembly graph when confining BLAST hits to a minimum query coverage and nucleotide identity 95%. Across *E. coli* and *S*. Typhimurium genomes, we constantly saw that the accuracy rate reached ≥ 90% under an error tolerance ±1 kbp when SPDs were measured within 250 kbp and no more than two nodes (Figures s9 and s11).

Moreover, since the accuracy rate of SPDs measured within contigs stayed above 90% when tolerating errors within ±1 kbp (Figures s10 and s12), we implemented prioritisation of SPDs based on their sources (namely, contigs or assembly graphs) in order to exclude inaccurate SPDs from assembly graphs where repeats have not been resolved by the assembler. Specifically, when the physical distance between two CDSs is measurable in both a contig and an assembly graph of the same genome, this method overrides the graph-based SPD with its corresponding contig-based SPD, thereby taking advantage of both the high accuracy rate of contig-based SPDs and high measurability of graph-based SPDs. As shown in Tables s5 and s6, the prioritisation method led to an accuracy rate of 100% for the majority of SPDs measured between acquired AMR genes, which are often embedded within tangled sub-graphs owning to surrounding repeats.

We found that inaccurate SPDs measured in two nodes of *Salmonella* assembly graphs (Table s6) were caused by chimeric alleles (that is, highly similar alleles of the same gene were mistakenly assembled into one allele) in the assembly, owing to the limited capacity of short reads in resolving repeats. We addressed this issue through increasing the threshold for both the nucleotide identity and query coverage of BLAST hits from 95% to 99% (BLAST hits to chimeric alleles were hence discarded), and improved the accuracy rate for two genomes at the cost of reducing distance measurability (Table s7). Accordingly, we applied this adjusted threshold to the measurement of SPDs between acquired AMR genes in the 358 *Salmonella* draft genomes. Besides, we directly calculated SPDs for strain DT104, whose complete genome is publicly accessible on GenBank.

#### SPDs between alleles of acquired AMR genes

From *de novo* assemblies of the 169 *E. coli* genomes, we obtained 1,550 SPDs for 301 allele pairs that were tested for associations (Hence alleles of each pair did not have identical presence-absence status across genomes) and 20 SPDs for nine pairs of identically distributed alleles. The largest SPD was 59,433 bp (measured across 17 nodes) and the greatest node traversal to measure an SPD was across 39 nodes (yielding an SPD of 4,628 bp). The exclusion of SPDs measured in more than two nodes resulted in 673 (43%) SPDs reliably measured for 163 tested allele pairs and 18 (90%) SPDs for eight pairs of identically distributed alleles. From *de novo* assemblies of the 359 *Salmonella* genomes, we obtained 2,880 SPDs for 224 allele pairs tested for associations, including 2,322 SPDs between alleles of the five SGI1-borne AMR genes (Table s8), and another 10 SPDs from five pairs of identically distributed alleles. The largest SPD was 710,475 bp (measured across 44 nodes) and the greatest node number was 57 (yielding an SPD of 687,051 bp). We saw large SPDs (> 56 kbp, which were extraordinarily larger than the others) when the node number exceeded 14. The exclusion of SPDs that were greater than 250 kbp and measured in more than two nodes resulted in 994 SPDs from 59 tested allele pairs and eight SPDs from three pairs of identically distributed alleles.

Overall, positively associated alleles of acquired AMR genes in *E. coli* and *Salmonella* genomes showed higher measurability of SPDs than those measured between negatively associated alleles. Specifically, for pairs of positively associated alleles, 64 (72%) out of 89 pairs in *E. coli* and 25 (26%) out of 97 pairs in *Salmonella* had at least two SPDs measured, respectively (Tables s9 and s10). Moreover, after the removal of SPDs greater than 250 kbp and measured in more than two nodes, 31 (35%) out of the 89 allele pairs in *E. coli* and 10 (10%) out of the 97 allele pairs in *Salmonella* had distance measurability above 75%. By contrast, no SPD was measurable between negatively associated alleles as these alleles did not co-occur in any genome.

#### Linkage networks showing support of SPDs to HGcoT

For each pathogen, we created a linkage network, in which nodes represent alleles or clusters of identically distributed alleles of acquired AMR genes and directed edges indicate significant associations obtained from LMMs (Figure 4). The edge width is proportional to an estimated effect size 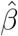, solid lines and dashed lines indicate positive associations and negative ones, respectively, and the edge colour indicates the distance score *s_d_*. No filter was applied to distance scores. The linkage network for *E. coli* consisted of 122 edges linking 46 nodes corresponding to 52 alleles of 26 acquired AMR genes. The distance score followed a bimodal distribution, with 57 out of 83 allele pairs (69%) having *s_d_* = 0 (84 edges) and 23 (28%) allele pairs having *s_d_* > 0.5 (38 edges). Considering only the edges that yielded *s_d_* > 0.5 and connected alleles encoding distinct kinds of resistance phenotypes, aminoglycoside resistance alleles linked to 14 alleles – the greatest number of connections, followed by sulphonamide resistance alleles (linked to 11 alleles). By contrast, *bla*_OXA-1_, the only beta-lactam resistance allele having edges with distance scores above 0.5, was linked to two alleles (*aadA1-pm*.182 and *catA1.215*). Notably, as shown in Figure 4, five alleles (*dfrA14.227, strB, sul2.168, strA.173* and *sul2*) formed a cluster that was interconnected by bidirectional edges with high distance scores (*s_d_* > 0.6).

The linkage network for *Salmonella* consisted of 37 alleles (21 AMR genes) connected by 162 edges (Figure 4). No identically distributed alleles could be collectively represented by a single node in this network due to absence of perfect measurability or consistency of SPDs. The distance score again followed a bimodal distribution, with 95 out of 104 (91%) pairs having *s_d_* < 0.05, and seven (7%) pairs (nine alleles in total) having *s_d_* > 0.5 (13 edges), all of which corresponded to significant positive associations. Considering only the 13 edges having distance scores above 0.5, *strB* (aminoglycoside resistance) linked to the greatest number of alleles (four, altogether), followed by *sul2* and *dfrA14.79*, each connected to two alleles.

**Figure 4:**
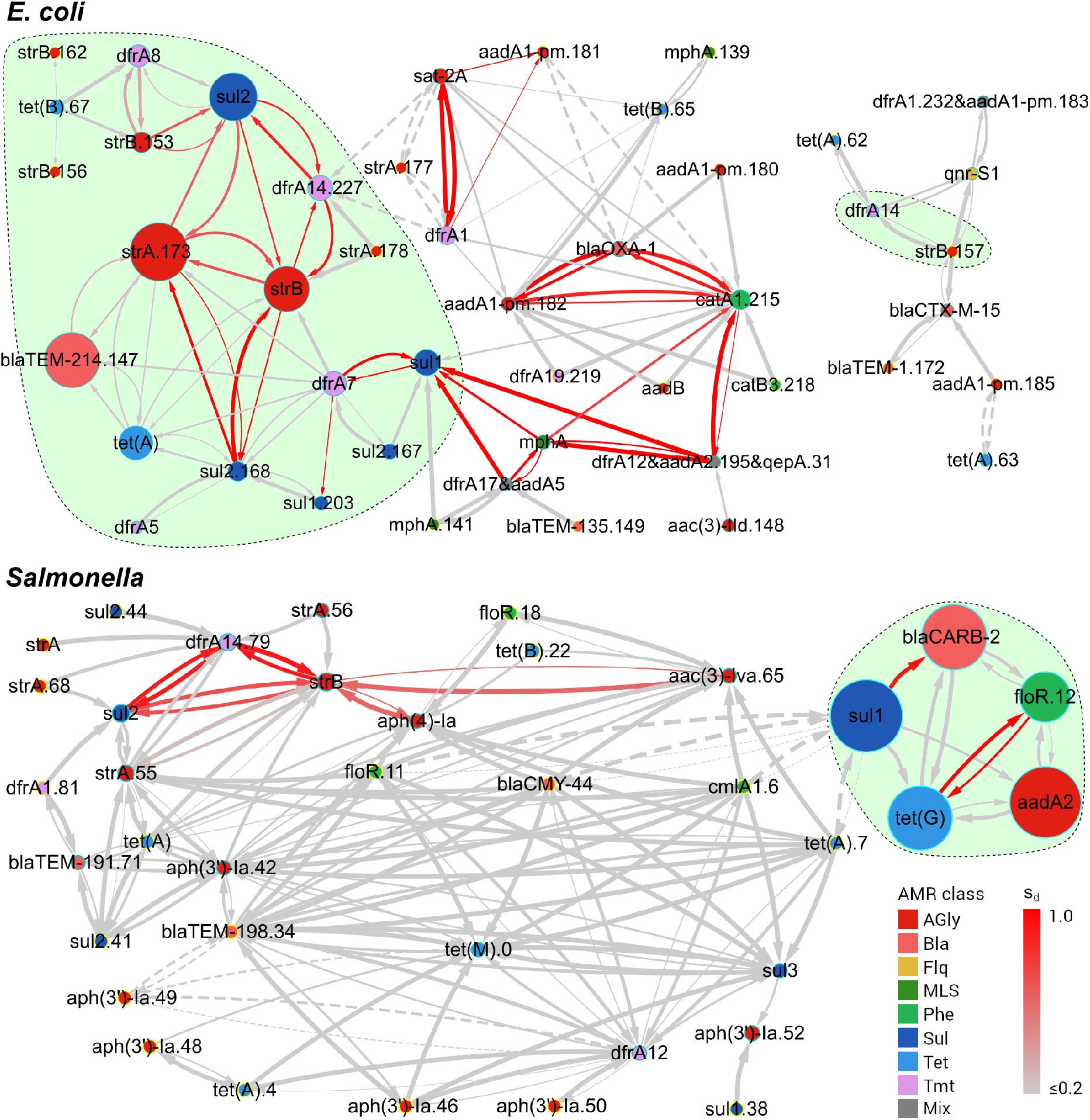
Linkage networks for detected alleles of acquired AMR genes in 169 E. *coli* and 359 *Salmonella* genomes. Each node represents an allele or a cluster of identically distributed alleles, with a diameter proportional to the allele frequency and a colour indicating the AMR phenotype encoded. Every edge is directed, starting from an explanatory allele and terminating at a response allele, representing a significant association determined using an LMM. The edge width is proportional to 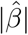 of the explanatory allele in an LMM. Solid edges represent significant positive associations 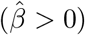 while dashed edges represent significant negative associations 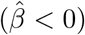. The edge colour follows a gradient of the distance score *s_d_* to indicate the strength of evidence for physical linkage. The shaded area (light green) encircles alleles of known co-transferred AMR genes. AMR classes defined by antimicrobials that were resistant to: AGly (aminoglycosides), Bla (beta-lactams), Flq (fluoroquinolones), MLS (macrolides, lin-cosamides, and streptogramins), Phe (phenicols), Sul (sulfonamides), Tet (tetracyclines), Tmt (trimethoprim), and Mix (multiple classes of antimicrobials).Reasons for inconsistency in measured physical distances In both the *E. coli* and *Salmonella* data sets, a lack of consistency in SPDs (namely, c = 0) measured between several positively associated alleles of acquired AMR genes was observed (Tables s9 and s10). We investigated this issue based on GeneMates outputs and identified two common explanations.

#### Reasons for inconsistency in measured physical distances

In both the *E. coli* and Salmonella data sets, a lack of consistency in SPDs (namely, *c* = 0) measured between several positively associated alleles of acquired AMR genes was observed (Tables s9 and s10). We investigated this issue based on GeneMates outputs and identified two common explanations.

First, in many cases diverse genetic structures were found carrying the same combination of alleles of AMR genes. For example, using *de novo* assembly graphs of *E. coli* genomes, we recovered six distinct genetic structures linking alleles *bla*_TeM-214_.147 and *tet*(A), which showed significant positive associations in both LMMs and PLMs but had six distinct SPDs (Figure s13). We found that these SPDs followed a lineage-specific distribution. As illustrated in Figure s14, the allele *bla*_TeM-214_.147 was carried by transposon Tn2, which was common to all the six structures, either in its complete or truncated form. The variety of insertion sites and orientations of this transposon relative to the allele *tet*(A) in *E. coli* genomes, as well as plausible gene gain/loss events inferred from structural comparisons, resulted in differences in SPDs measured between the two alleles.

Second, the GeneMates algorithm depreciates physical distances showing IBD for scoring the distance consistency (Section 3.3.2 of Additional File 1). For instance, alleles of SGI1-borne AMR genes *sul1* and *aadA2* in *Salmonella* were frequent amongst the 359 *Salmonella* isolates, with an occurrence count 328 (91%) and 323 (90%), respectively (Table s8). Co-occurrence of these two alleles were also frequent, with a count 318 (89%) in total. SPDs between *sul1* and *aadA2* were obtained from genome assemblies of 295 (93%) out of the 318 isolates where the alleles were co-occurring. After removing the only SPD measured across more than two nodes (504 bp, across three nodes), we obtained 294 SPDs, consisting of 293 SPDs measured in either contigs or assembly graphs and a single SPD measured in the complete chromosome genome of the reference strain DT104. All the filtered SPDs were 504 bp, except the one from the complete genome (9,964 bp). Despite this consistency in SPDs, the consistency score c = 0 as all the 294 SPDs were obtained from the same lineage highlighted in Figure s15a (IBD probability of reliable SPDs: 95%).

### 3.4 Validation of GeneMates

We validated our approach by examining known and novel physical clusters of mobile AMR genes in the example data sets. Networks were constructed at the allele level for these genes using the GeneMates function *findPhysLink*.

#### Identifying known clusters of mobile AMR genes

The first set of positive controls in our validation study consisted of 28 pairs of AMR genes that are known to be co-mobilised by MGEs between *E. coli* genomes [19]. As shown in Table 1, LMMs and PLMs identified significant positive associations at the allele level in 19 (68%) and 16 (57%) pairs, respectively. The second set of positive controls for validation consisted of five AMR genes (*aadA2, floR, tet*(G), *bla*_CARB-2_, and *sul1*) that are co-localised in the acquired multidrug-resistant element SGI1 in *Salmonella* genomes [20]. For these genes, LMMs and PLMs identified significant positive associations between eight and ten allele pairs, respectively (Table 2). The exclusion of allele pairs having *s_d_* < 0.6 led to a substantial reduction of co-mobilisation candidates, with 12 out of 33 (36%) allele pairs in *E. coli* genomes and two out of 20 (10%) allele pairs in *Salmonella* genomes passed this filter.

**Table 1:**
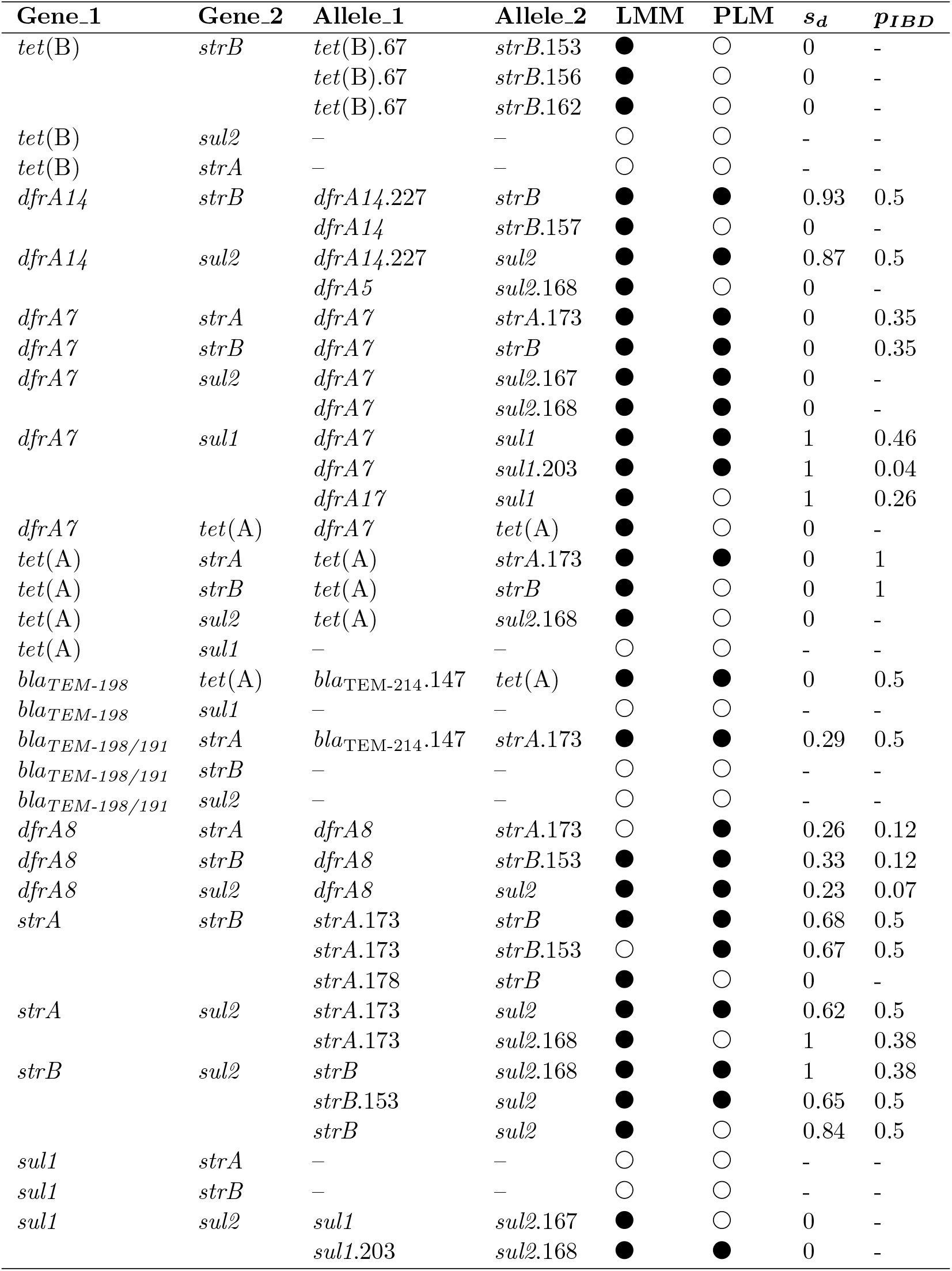
A comparison of significant positive associations in the network for *E. coli* genomes to known co-localisation of mobile AMR genes. Co-localisation of AMR genes in MGEs were previously determined by Ingle, et al. [19]. Each pair of significantly associated alleles (denoted by alleles 1 and 2, regardless of their roles in a linear model) is identified in the network shown in Figure 3. Directionality of associations is omitted in this table, hence each pair of alleles only appears once in the table, although the alleles may mutually associated in linear models. Notice an AMR gene may have multiple alleles (whose names are listed in the column Allele_1 or Allele_2) or no allele (denoted by a dash sign in the table) present in the association network. Abbreviations: LMM, linear mixed model; PLM, penalised logistic model. Symbols indicating whether a significant association is determined using either an LMM or a PLM: (●), yes; (○), no. *s_d_*: the distance score, which takes into account the distance consistency, measurability, and the probability of IBD.*_pIBD_*: an estimate of the probability that APDs used for calculating the *s_d_* are in IBD. This probability does not exist when no APD is available.

**Table 2:** LMM-based significant associations between five alleles of AMR genes in SGI1. Allele_Y and Allele_X denote the response allele and explanatory allele in an LMM *Y* ~ *X*, respectively. An association is denoted by a filled circle in the column LMM when it is significant, otherwise, an unfilled circle is drawn. Directionality is shown in this table in order to compare the value of 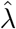, which denotes an REML estimate of the parameter λ for random structural effects in an LMM. *s_d_*: score of APDs. *_pIBD_*: an estimate of the probability that APDs used for calculating the *s_d_* are in IBD.

| Gene_Y | Gene_X | Allele_Y | Allele_X | $\hat{\lambda}$ | LMM | PLM | $s_d$ | $p_{IBD}$ |
| --- | --- | --- | --- | --- | --- | --- | --- | --- |
| *floR* | *sul1* | *floR*.12 | *sul1* | 64.98 | ○ | ● | 0.03 | 0 |
| *sul1* | *floR* | *sul1* | *floR*.12 | $\geq 10^5$ | ○ | ● | 0.03 | 0 |
| *aadA2* | *floR* | *aadA2* | *floR*.12 | 137.49 | ● | ● | 0.02 | 0 |
| *floR* | *aadA2* | *floR*.12 | *aadA2* | 82.73 | ● | ● | 0.02 | 0 |
| *bla*<sub>CARB</sub> | *floR* | *bla*<sub>CARB-2</sub> | *floR*.12 | 110.28 | ● | ● | 0.04 | 0 |
| *floR* | *bla*<sub>CARB</sub> | *floR*.12 | *bla*<sub>CARB-2</sub> | 57.27 | ● | ● | 0.04 | 0 |
| *floR* | *tet*(G) | *floR*.12 | *tet*(G) | 64.45 | ● | ● | 0.99 | 0.5 |
| *tet*(G) | *floR* | *tet*(G) | *floR*.12 | 91.3 | ● | ● | 0.99 | 0.5 |
| *bla*<sub>CARB</sub> | *sul1* | *bla*<sub>CARB-2</sub> | *sul1* | 70.79 | ● | ● | 0.92 | 0.85 |
| *sul1* | *bla*<sub>CARB</sub> | *sul1* | *bla*<sub>CARB-2</sub> | $\geq 10^5$ | ○ | ● | 0.92 | 0.85 |
| *aadA2* | *bla*<sub>CARB</sub> | *aadA2* | *bla*<sub>CARB-2</sub> | 130.1 | ○ | ● | 0 | 0 |
| *bla*<sub>CARB</sub> | *aadA2* | *bla*<sub>CARB-2</sub> | *aadA2* | 123.6 | ○ | ● | 0 | 0 |
| *bla*<sub>CARB</sub> | *tet*(G) | *bla*<sub>CARB-2</sub> | *tet*(G) | 244.33 | ● | ● | 0.04 | 0 |
| *tet*(G) | *bla*<sub>CARB</sub> | *tet*(G) | *bla*<sub>CARB-2</sub> | 160.14 | ● | ● | 0.04 | 0 |
| *sul1* | *tet*(G) | *sul1* | *tet*(G) | $\geq 10^5$ | ○ | ● | 0.03 | 0 |
| *tet*(G) | *sul1* | *tet*(G) | *sul1* | 55.9 | ● | ● | 0.03 | 0 |
| *aadA2* | *tet*(G) | *aadA2* | *tet*(G) | 118.93 | ● | ● | 0.04 | 0 |
| *tet*(G) | *aadA2* | *tet*(G) | *aadA2* | 77.76 | ● | ● | 0.04 | 0 |
| *aadA2* | *sul1* | *aadA2* | *sul1* | 73.42 | ● | ● | 0 | 0.95 |
| *sul1* | *aadA2* | *sul1* | *aadA2* | $\geq 10^5$ | ○ | ● | 0 | 0.95 |

#### Identifying novel clusters of AMR genes

We noticed that some edges in linkage networks possibly indicated novel physical linkage between several alleles of AMR genes in *E. coli* and *Salmonella* genomes. For instance, 20 novel edges in the linkage network for *E. coli* (Figure 4) had the maximum association score 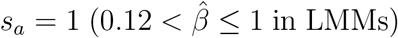 and high distance scores (0.75 < *s_d_* < 1), and these edges formed five maximal cliques each consisting of three alleles. Similarly, a three-allele clique showing high distance scores (0.7 < *s_d_* < 0.9) was identified in the linkage network for *Salmonella* (Figure 4). As a further validation of GeneMates, we investigated two maximal cliques through resolving their genetic structures and vectors in genome assemblies.

The first clique consisted of alleles *dfrA1, aadA1-pm.181*, and *sat-2A* detected in *E. coli* genomes. As illustrated in Figure s16a, LMMs identified significant positive associations between alleles this clique. Moreover, we saw perfect identity and measurability in APDs between these alleles (that is, *s_d_* =1 for every edge of this clique, see Table s11 for more details). The combination of these three alleles occurred in two genomes belonging to two distantly related lineages (Figure s16b), with the alleles present in the same gene-cassette array of a class-2 integron (Figure 5a, b). Further, we confirmed that this integron was carried by variants of a Tn*7* transposon (100% coverage under a nucleotide identity of 99%), each interrupted by an insertion of a distinct IS element (Figure 5c). Therefore, there is strong evidence that these alleles were co-transferred between *E. coli* lineages by the MGE Tn*7*.

**Figure 5:**
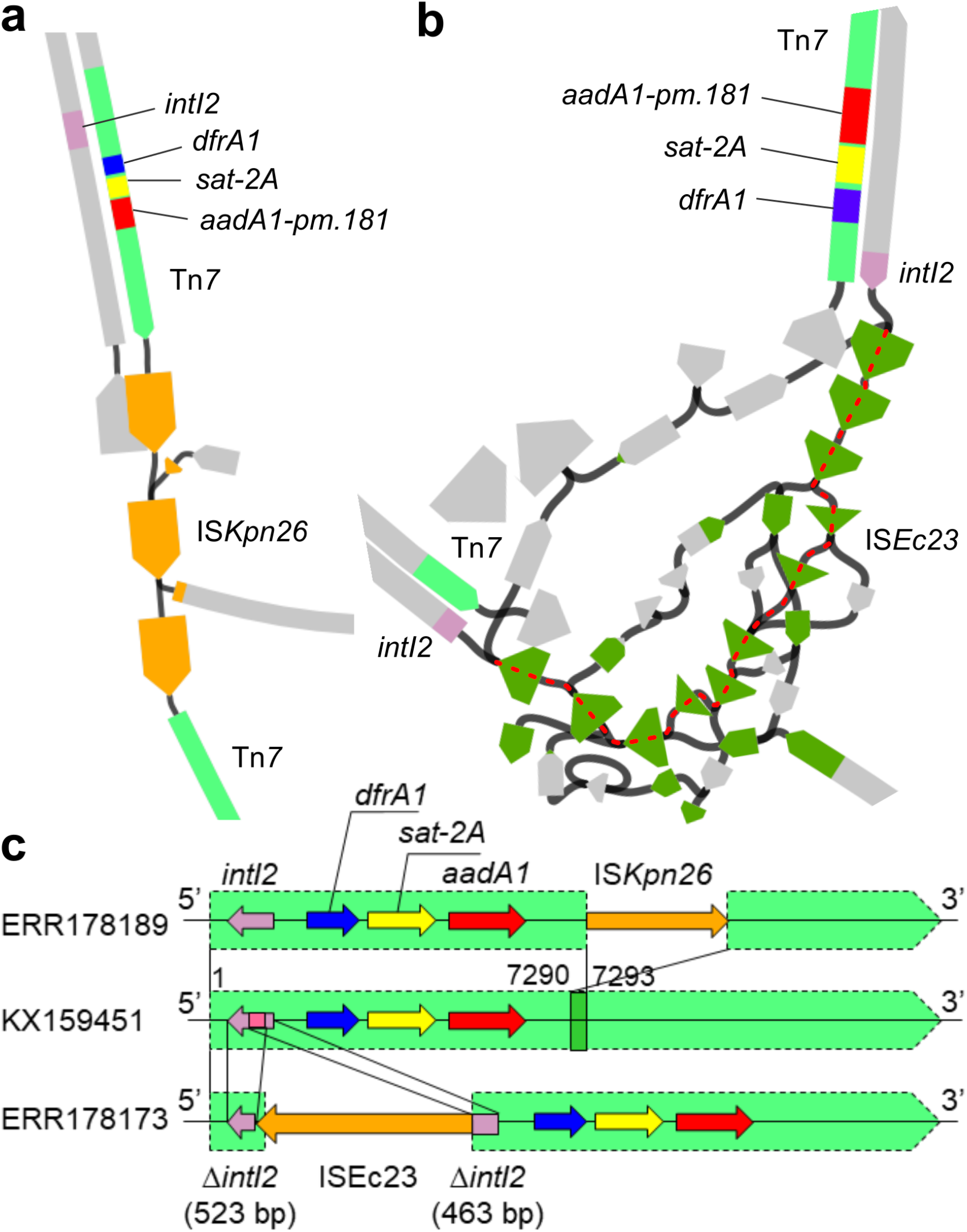
Putative physical linkage between three resistance alleles in two *E. coli* genomes. (**a**) A path constituting an inferred MGE in the assembly graph of genome ERR178189. This plot is drawn with Bandage in a double-strand style, where the orientation of each DNA strand is indicated by an arrow-like node end. The width of each node is proportional to its read depth. Some nodes not contributing to any MGE-related paths were deleted from this graph for visual conciseness. (**b**) A path constituting the other inferred MGE (following the red dashed line) in the assembly graph of genome ERR178173. This plot was drawn in the same way as panel a. (**c**) Alignment of the two Tn*7* variants in *E. coli* genomes (ERR178189 and ERR178173) to a reference Tn*7* sequence (GenBank accession: KX159451, denoted by the green shaded areas). Two direct repeats flanking the ISs, including inverted repeats, are denoted by green and pink boxes, respectively. Reference DNA sequences of IS*Kpn26* (1,196 bp) and IS*Ec23* (2,532 bp) were retrieved from the ISFinder database [26] in January 2018. Each IS in the resolved region showed a 100% coverage and a nucleotide identity of 99% to its reference.

The second clique consisted of alleles *str*A.55, *strB, dfrA14.79*, and *sul2* detected in *Salmonella* genomes (Figure 4). Both LMMs and PLMs identified that associations between all of these alleles were significantly positive 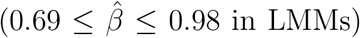. Edges between these alleles had distance scores between 0.7 and 0.9 except edges linking strA.55 (0 < *s_d_* < 0.25). As shown in Figure 6a, these four alleles co-occurred in 13 *Salmonella* genomes from distantly related clades. Using Bandage and nucleotide BLAST, we confirmed co-localisation of these alleles in a 3,084 bp region that was present in 10 out of the 13 genomes under a 100% nucleotide identity and coverage, and these 10 genomes were sparsely distributed across the phylogenetic tree (Figure 6a). Furthermore, we saw an insertion of the allele *dfrA14.79* into strA.55, splitting the latter allele into two segments that covered 65.8% and 34.5% of its length, respectively. In the assembly graph of one of the 10 genomes, ERR026101, we found the 3,084 bp multidrug-resistance (MDR) region in a 6,790 bp node, which appeared as a self-circularised sequence independent to other graph components (Figure 6b). Using megaBLAST under its default parameters, a sequence search of this MDR region against the NCBI nucleotide database of the *Enterobacteriaceae* group (taxid: 91347, accessed in April, 2018) showed exact matches (100% nucleotide identity and coverage) to a known and widely distributed MDR plasmid pCERC1 (GenBank accession: JN012467) as well as a number of plasmids widely distributed in bacteria of *Enterobacterales* (Table s12). Hence this MDR region is shared amongst a great variety of plasmids.

**Figure 6:**
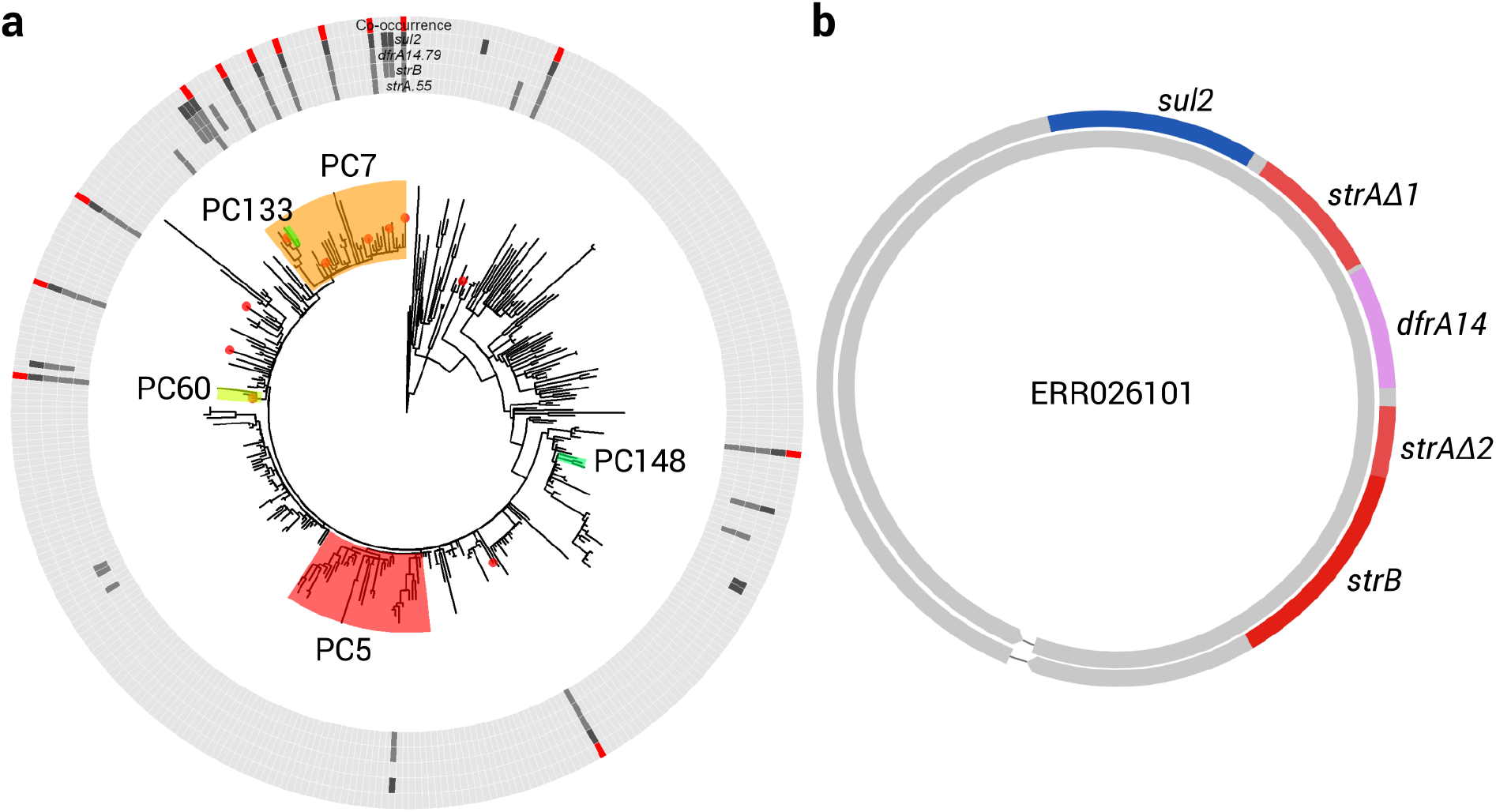
Distribution and genetic structure of the four-allele clique in *Salmonella* genomes. (**a**) A ring plot displaying co-occurrence events (red ring) of the four alleles (darker boxes in grey rings) against the midpoint rooted phylogeny of the *Salmonella* genomes. Clades are coloured by their top-10 most correlated PCs that are significantly explaining the presence-absence status of an arbitrarily designated response allele *sul2*. Tips highlighted with red circles denote genomes where the exact 3,084 bp MDR region harboured all the four alleles were found using nucleotide BLAST. (**b**) A putative 6,790 bp plasmid sequence restored from the assembly graph of genome ERR026101. The resistance alleles were present in this sequence at a nucleotide identity and coverage of 100%.

## 4 Discussion

GeneMates implements a novel network approach to detection of HGcoT between bacteria. Compared to existing methods that rely on co-occurrence counts or naive association coefficients of AMR genes, GeneMates enables us to test for gene-gene associations when controlling for population structure, the main confounding factor in bacterial GWAS [27], through incorporating PCs into LMMs. It has been shown that this procedure can retain statistical power of association tests [10, 28]. In our examples, association networks constructed using GeneMates reveal extensive associations between alleles of horizontally acquired AMR genes (Figure 3). Moreover, as expected, the majority of p-values from association tests became greater after correcting for population structure using LMMs, while the other p-values saw a reduction, indicating increased significance of associated alleles (Figure 2). LMMs provide us with another advantage: we only need to estimate three parameters (*β, γ* and *τ*) besides the intercept term *α* to fit a model, hereby circumventing the problem of over-fitting as well as relaxing the requirement for sample size. Notably, long-tailed curves of cumulative percentages of total genetic variations captured by PCs (Figure s8) indicates the necessity of including all PCs obtained from the cgSNP matrix for accurate modelling in association analysis.

On the grounds of examples where stable structures of acquired AMR genes are shared between bacteria via MGEs in a short period of time [16, 19, 20, 29], we have implemented another innovation in GeneMates - the evaluation of APD consistency for evidence of HGcoT. Since the physical distances between two loci in bacterial genomes are correlated with both locus co-occurrence and PCs representing population structure, APDs cannot be directly incorporated into LMMs or PLMs as additional covariates. Instead, GeneMates scores the variation of APDs while taking the population structure into account. As for APDs, it is self-evident that they can be precisely calculated from genetic coordinates in finished-grade genome assemblies, which remain the minority of available bacterial genome sequences. In order to overcome some of the limitations of measuring APDs in draft genomes, we measured APDs in the form of SPDs in assembly graphs, and developed a simulation-based approach to determining reliability filters of SPDs. Using this approach, the accuracy of SPDs from *de novo* assembly graphs of reference multidrug-resistant *E. coli* or *S*. Typhimurium genomes consistently exceeded 90% when the distances were only measured in one or two nodes (Figures s9 and s11), implying a universal filter for other bacterial genomes. Nevertheless, as summarised in Section 3.3 and displayed in Figure 4, the short-read genome assemblies intrinsically confine both the measurability and accuracy of SPDs, losing evidence for real physical linkage in HGT.

In Section 3.4, the identification of known and novel physical clusters of mobile AMR genes suggests that maximal cliques in association networks are useful starting points for recovering structures of horizontally co-transferred genes. A plausible reason is that loci in strong physical linkage tend to predict the presence of each other (namely, mutual positive associations) in HGT, and strong physical linkage is often related to close physical proximity, which in turn increases the measurability of their physical distances, leading to a greater distance score given the same consistency score. In practice, users could apply other filters to the network to identify edges addressing specific questions.

GeneMates offers a framework (Figure 1) for further development, such as adding new modules and introducing other statistical models. Our methodology and the analysis demonstrated in example studies are applicable to other kinds of acquired genes in haploid genomes, as long as we can accurately determine the genotype of each genome. Nevertheless, the inability of short reads to resolve repeats, either through read mapping or *de novo* assembly, may cause false negatives and errors in allele calls when homologues of a target gene coexist in a genome. Therefore, we expect a better performance of GeneMates in analysing high-quality complete genomes or long-read sequencing data that are able to resolve at least most of the repeats. Further, biological experiments are necessary as a gold standard for validating candidates of HGcoT.

## 5 Conclusions

We have developed an R package called GeneMates and supplementary tools for analysing associations between allelic presence-absence status of acquired AMR genes and for inferring physical linkage between these alleles. We have also demonstrated utilities of this package using publicly available WGS data of 169 *E. coli* isolates and 359 *Salmonella* isolates. Functions of the package reported known co-localised resistance alleles and discovered their distributions in certain bacterial species. GeneMates differs from contemporary bacterial GWAS tools in three aspects. First, it focuses on gene-to-gene associations rather than genotype-to-phenotype associations. Second, it only performs association tests for acquired genes rather than genome-wide single-nucleotide polymorphisms (SNPs). Third, it evaluates evidence of physical distances between associated loci to infer physical lin-kage, although a user may opt to turn this utility off for pure gene-to-gene associations. This is a scalable and versatile approach, readily applicable to other kinds of horizontally acquired genotypes. It is, however, confounded by limitations of short-read assembly, and its power will increase in the future as we are accumulating complete genomes and enhancing our ability in resolving repeats using sequencing data.

## Supporting information

Details of bacterial genomes used in this study

Detailed results of prophage detection, read mapping, and de novo genome assembly

## 6 Availability and requirements

- **Project name:** GeneMates
- **Project home page:** github.com/wanyuac/GeneMates
- **Programming language:** R
- **Operating system(s):** platform independent
- **Other requirements:** GEMMA v0.96
- **Licence:** Apache License, Version 2.0

## 7 Declarations

### Ethics approval and consent to participate

Not applicable.

### Consent for publication

Not applicable.

### Availability of data and materials

All sequence data analysed during this study are publicly available in NCBI databases. See Additional File 2 for accession numbers.

### Authors’ contributions

YW derived algebra of our methodology, developed and validated GeneMates and its helper scripts, curated and analysed the test data, interpreted results, and prepared the manuscript for publication. RRW developed and implemented the method for measuring APDs in assembly graphs, and provided innovative suggestions on short-read simulation and sequence analysis. KEH and MI conceived the methodology and this project. KEH, MI and JZ supervised this project and substantially contributed to research strategy and result interpretation. DJI participated in study design and result interpretation. All authors read and approved the final manuscript as well as contributing to revisions.

## Acknowledgements

We thank Kelly Wyres and David Edwards (Monash University), Claire Gorrie (University of Melbourne), and Gittan Blezer (University of Melbourne and Utrecht University) for their suggestions on data collection, quality control and read mapping. We appreciate Guoqi Qian (University of Melbourne) for his comments on our statistical methods. We also express our gratitude to the maintenance team of GEMMA (github.com/genetics-statistics/GEMMA) and the GitHub community for constructive discussions about software usage and statistics.

## 8 Additional Files

**Additional file 1 – Supplementary tables and figures, implementation details of GeneMates:**

A PDF document consisting of supplementary tables and figures, and a rigorous mathematical justification of statistical methods implemented in the package GeneMates.

**Additional file 2 – Sample information:** A spreadsheet of sample information, including database accessions, sampling location and year, sequencing layout, and so forth. It consists of four tables with the names Ecoli_reads (Illumina read sets of *E. coli* isolates), STyphimurium_reads (Illumina read sets of *Salmonella* isolates), MDR_Ecoli_ref (GenBank records for complete genomes of 10 multidrug-resistant *E. coli* isolates) and MDR_STyphimurium_ref (GenBank records for complete genomes of 10 multidrug-resistant *S*. Typhimurium isolates).

**Additional file 3 – Supplementary results**: An Excel spreadsheet for large tables of results.

