## Supplementary figures, tables, and methods for "GeneMates: an R package for Detecting Horizontal Gene Co-transfer between Bacteria Using Gene-gene Associations Controlled for Population Structure"

February 2020

GeneMates: an R Package for Detecting Horizontal Gene Co-transfer between Bacteria Using Gene-gene Associations Controlled for Population Structure

### 1 Supplementary figures

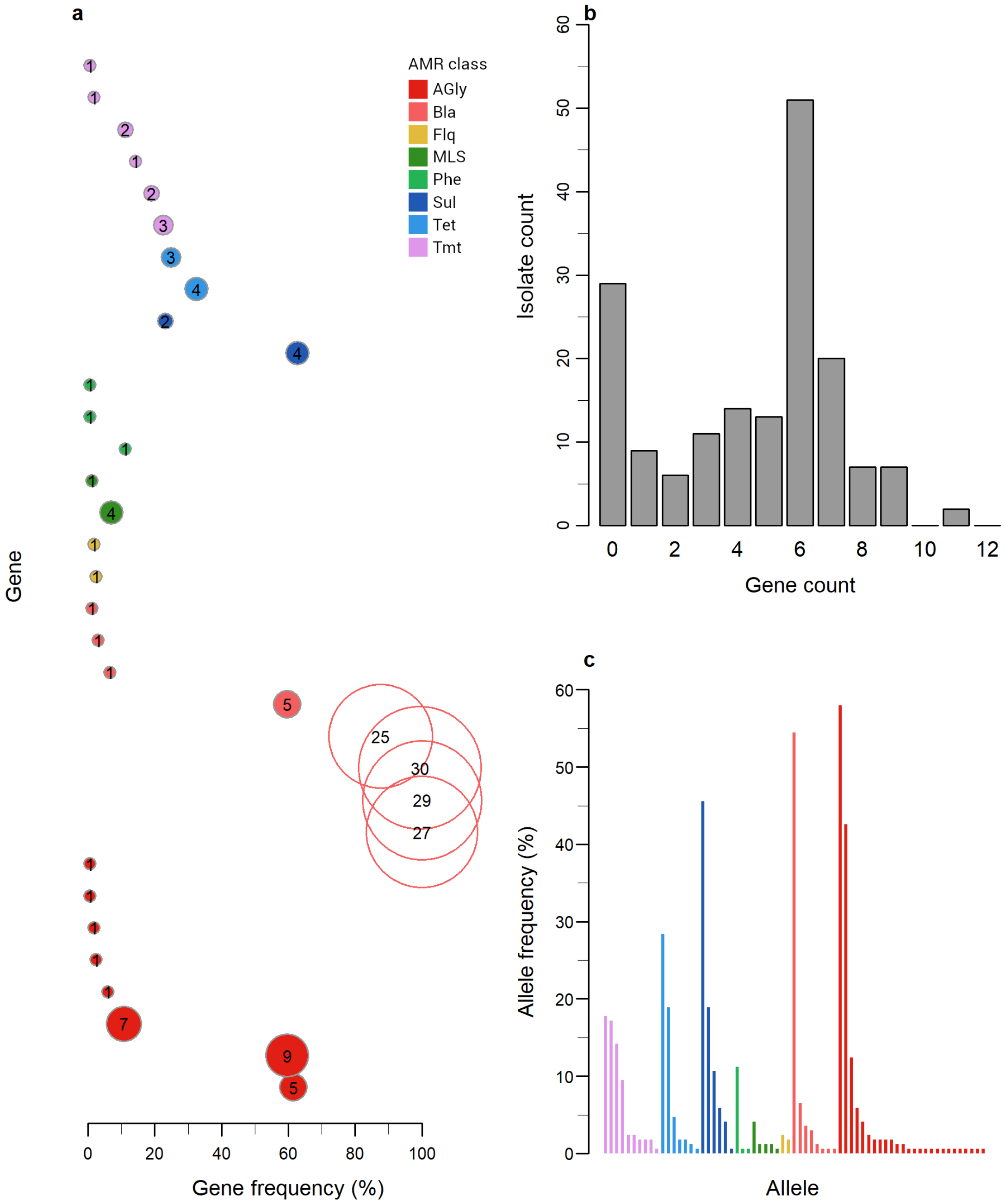

**Figure s1: Antimicrobial resistance (AMR) genes detected in the 169 *E. coli* genomes**. (**a**) Frequencies and allele numbers of AMR genes sorted by AMR classes. Each gene is represented by a circle, either shaded (accessory) or unshaded (intrinsic), and is coloured by the associated AMR class. The diameter of each circle is proportional to the allele number labelled on the circle. (**b**) The number of genomes harbouring a particular number of accessory AMR genes followed a bimodal distribution. (**c**) Frequencies of 178 alleles of accessory AMR genes arranged in a descending order within each AMR class. AMR classes: AGly, aminoglycosides; Bla, beta-lactams; Flq, fluoroquinolones; MLS, macrolides/lincosamides/streptogramins; Phe, phenicols; Sul, sulfonamides; Tet, tetracyclines; Tmt, trimethoprim.

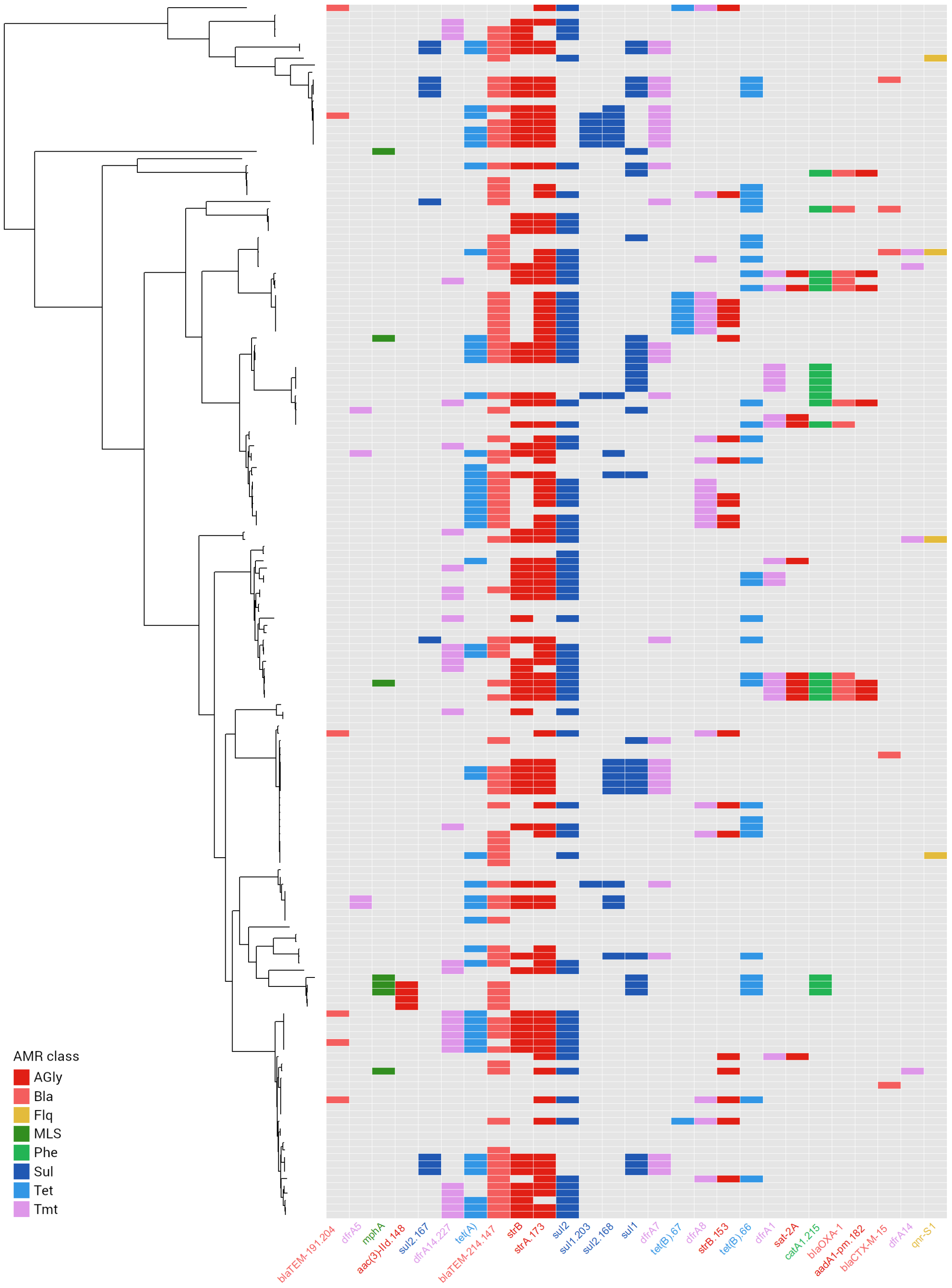

**Figure s2: A binary heat map showing presence-absence of 27 alleles of accessory AMR genes, each occurred at least four times in 169 *E. coli* genomes**. Alleles of lower frequencies are excluded from this heat map for conciseness. In the heat map, rows represent genomes whose relationships are indicated in the midpoint-rooted core-genome ML phylogeny; columns represent alleles, and are clustered using a single-linkage method based on binary distances between columns. In this heat map, each grey box indicates absence of an allele in a genome, and each coloured box indicates presence of an allele in a given genome, with the colour linked to an AMR class. AMR classes: AGly, aminoglycosides; Bla, beta-lactams; Flq, fluoroquinolones; MLS, macrolides/lincosamides/streptogramins; Phe, phenicols; Sul, sulfonamides; Tet, tetracyclines; Tmt, trimethoprim. Original data underlying this figure can be interactively visualised and downloaded from Microreact [1] following the link microreact.org/project/ifrfPqG_u.

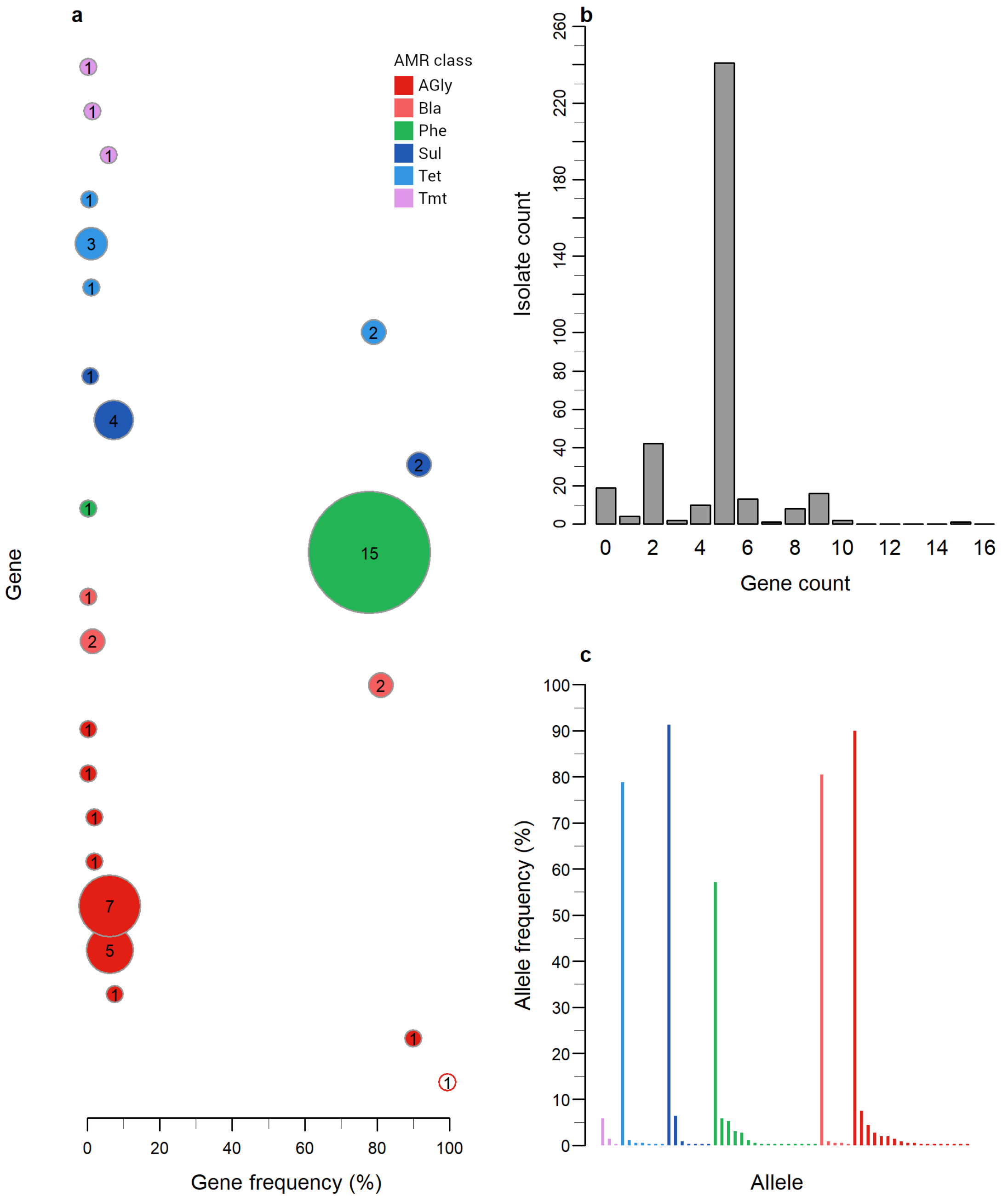

**Figure s3: ARG content in 359 *Salmonella* genomes**. (**a**) Frequencies and allele numbers of AMR genes sorted by AMR classes. Each gene is represented by a circle, either shaded (accessory) or unshaded (intrinsic), and is coloured by the corresponding AMR class. (**b**) The number of genomes each harbouring a particular number of accessory AMR genes. (**c**) Frequencies of 56 alleles of 23 accessory AMR genes arranged in a descending order within each AMR class. AMR classes: AGly, aminoglycosides; Bla, beta-lactams; Phe, phenicols; Sul, sulfonamides; Tet, tetracyclines; Tmt, trimethoprim.

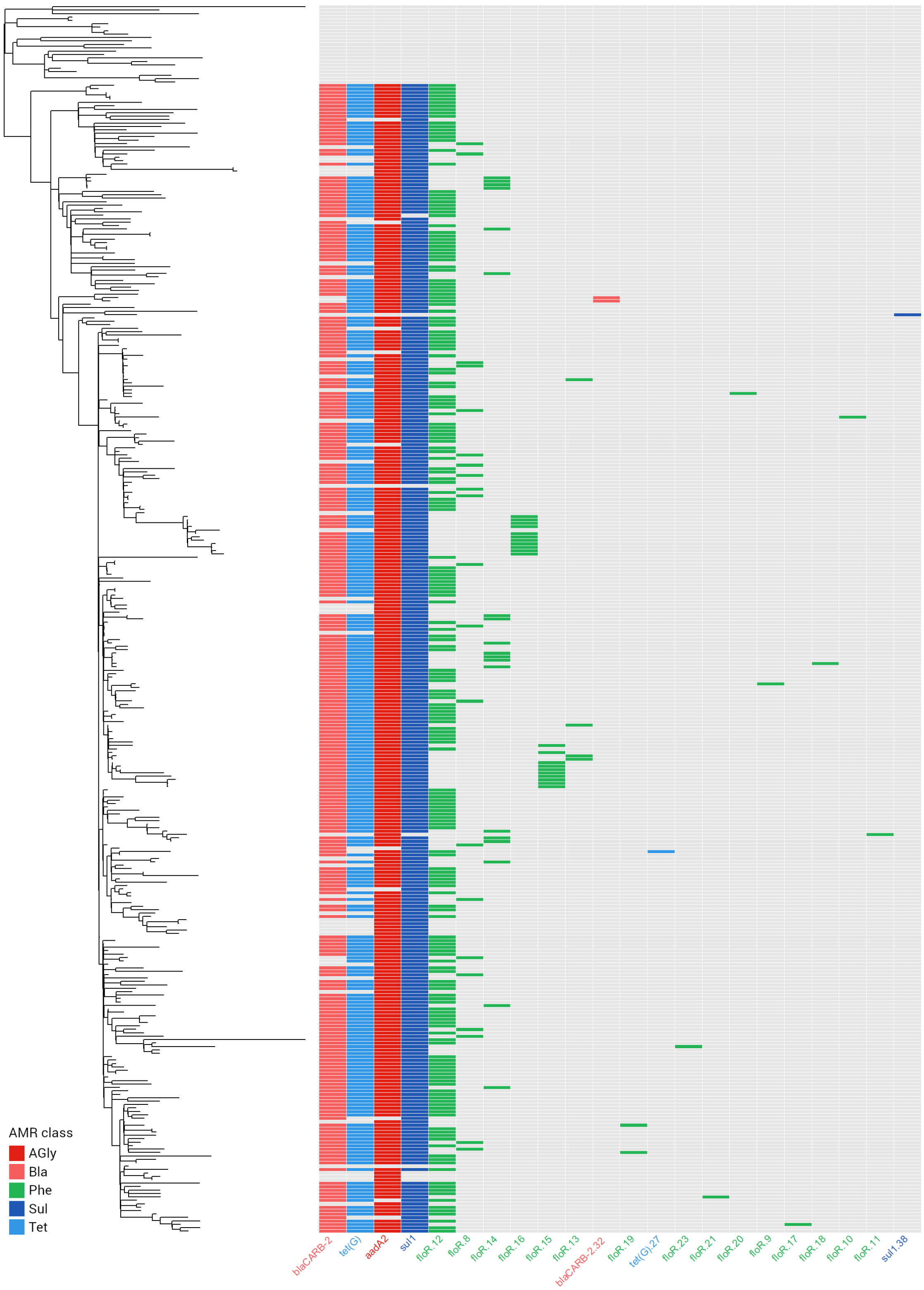

**Figure s4: A binary heat map showing presence-absence of 22 alleles of SGI1-borne AMR genes in 359 *Salmonella* genomes**. Rows represent genomes whose relationships are indicated in the midpoint-rooted core-genome ML phylogeny; columns represent alleles of five AMR genes. The columns are clustered using a single-linkage method based on binary distances between columns. Each grey box indicates absence of an allele in a given genome. AMR classes: AGly, aminoglycosides; Bla, beta-lactams; Phe, phenicols; Sul, sulfonamides; Tet, tetracyclines. Original data underlying this figure can be interactively visualised and downloaded from Microreact following the link microreact.org/project/0bGpp3D-y.

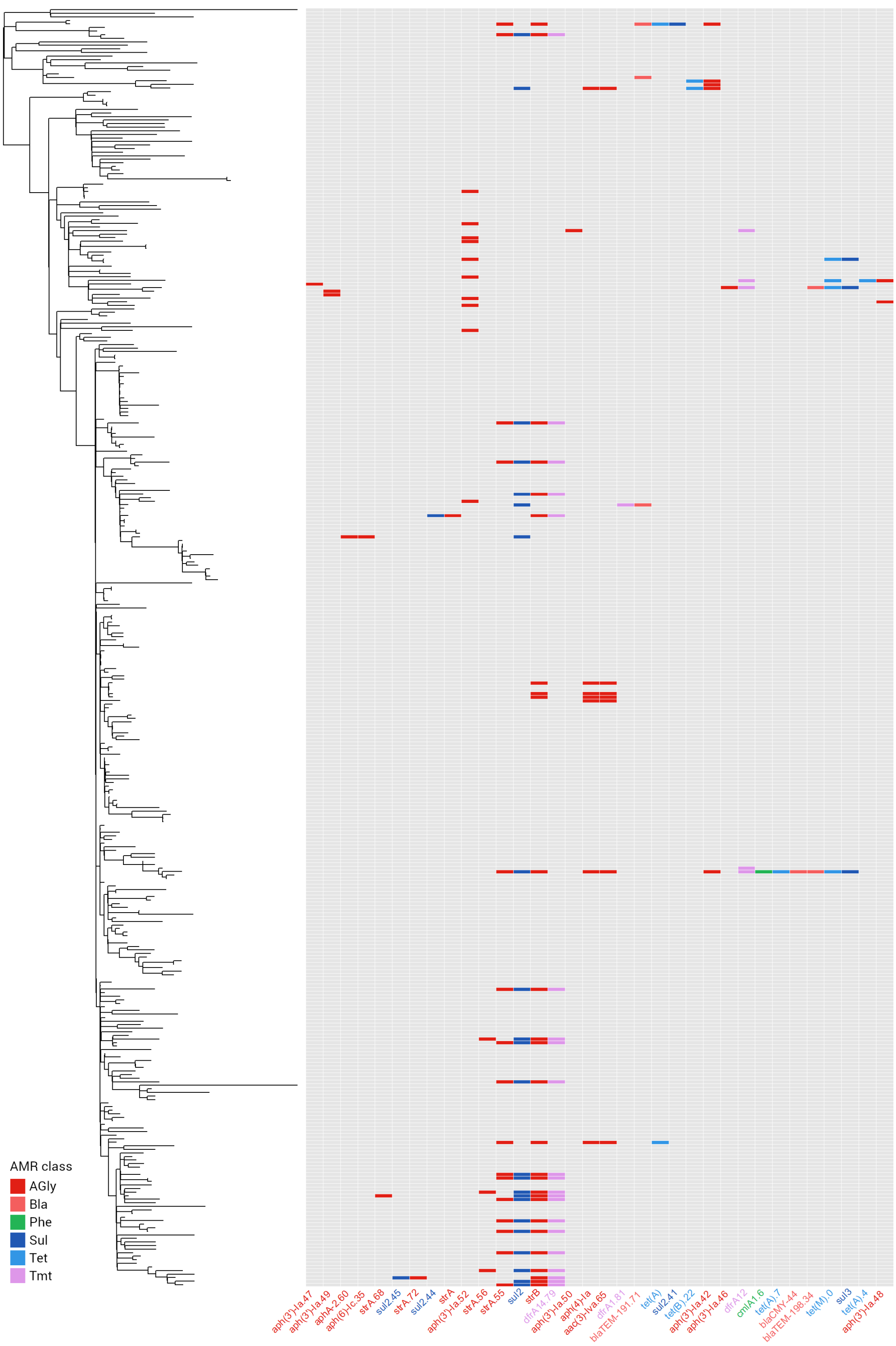

**Figure s5: A binary heat map showing presence-absence of 34 alleles of AMR genes that were not carried by SGI1 in 359 *Salmonella* genomes**. Rows represent genomes whose relationships are indicated in the midpoint-rooted core-genome ML phylogeny; columns represent alleles, which belonged to 18 AMR genes. The columns are clustered using a single-linkage method based on binary distances between the columns. Each grey box in the heat map indicates absence of an allele in a given genome. AMR classes: AGly, aminoglycosides; Bla, beta-lactams; Phe, phenicols; Sul, sulfonamides; Tet, tetracyclines; Tmt, trimethoprim.

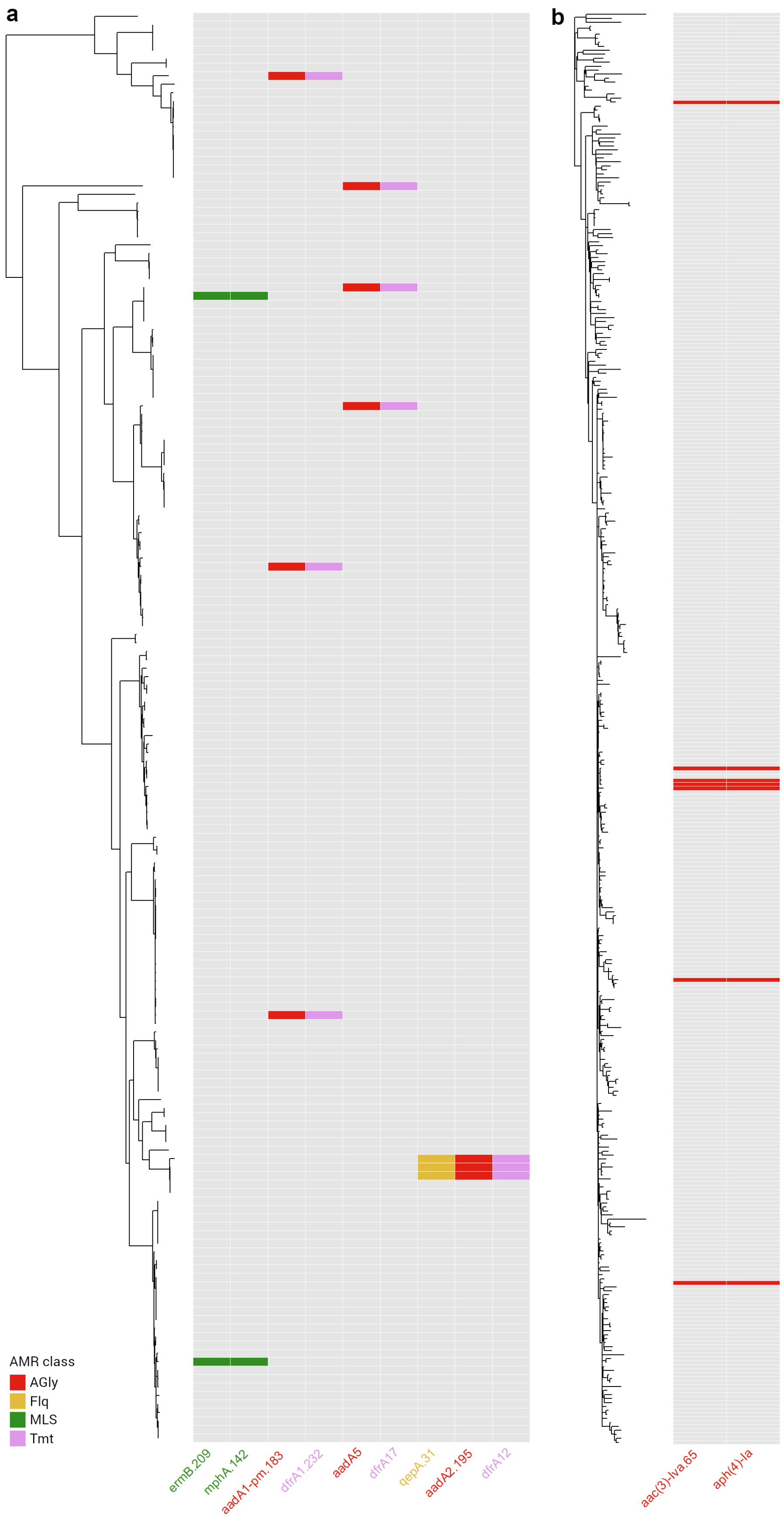

**Figure s6: Binary heat maps showing distributions of identically distributed alleles of accessory AMR genes in (a) *E. coli* or (b) *Salmonella*.** For each species, an ML phylogeny is shown on the left side of each heat map. AMR classes: AGly, aminoglycosides; Flq, fluoroquinolones; MLS, macrolides, lincosamides, and streptogramins; Tmt, trimethoprim.

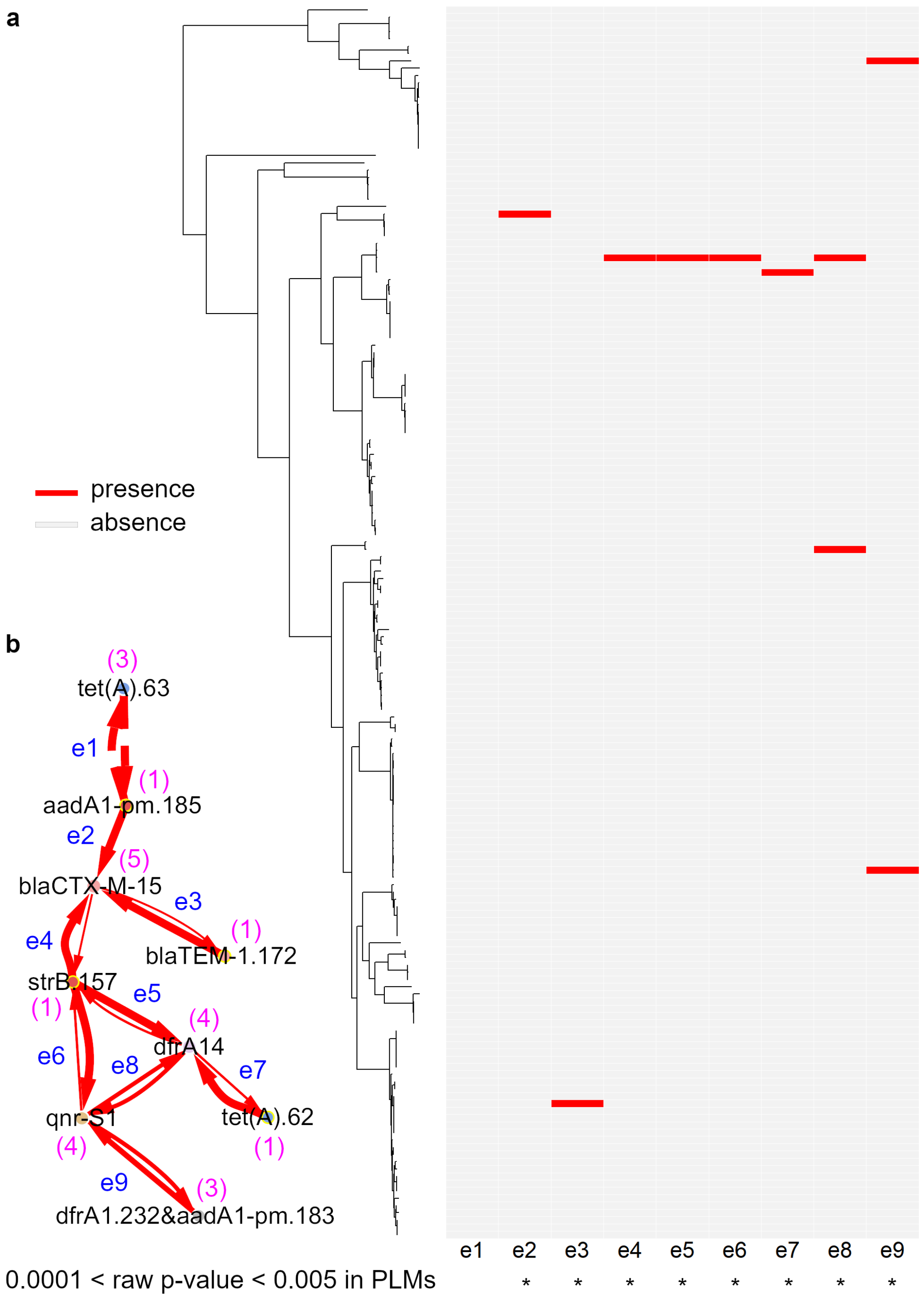

**Figure s7: A separate sub-network in the association network of *E. coli* (a) and co-occurrence of its alleles in 169 *E. coli* genomes (b).** In (**a**), a heat map showing co-occurrence events between nodes in the sub-network (**b**) and an alignment of these events against the midpoint-rooted ML core-genome phylogeny of *E. coli* genomes. Column names of the heat map denote node pairs labelled in (*b*). Counts of alleles in the 169 genomes are displayed as digits between parentheses next to node labels in (**b**). Asterisks beneath column names denote raw p-values (that is, without Bonferroni correction) based on PLMs. Note that PLM-based p-values were the same for each pair of these alleles.

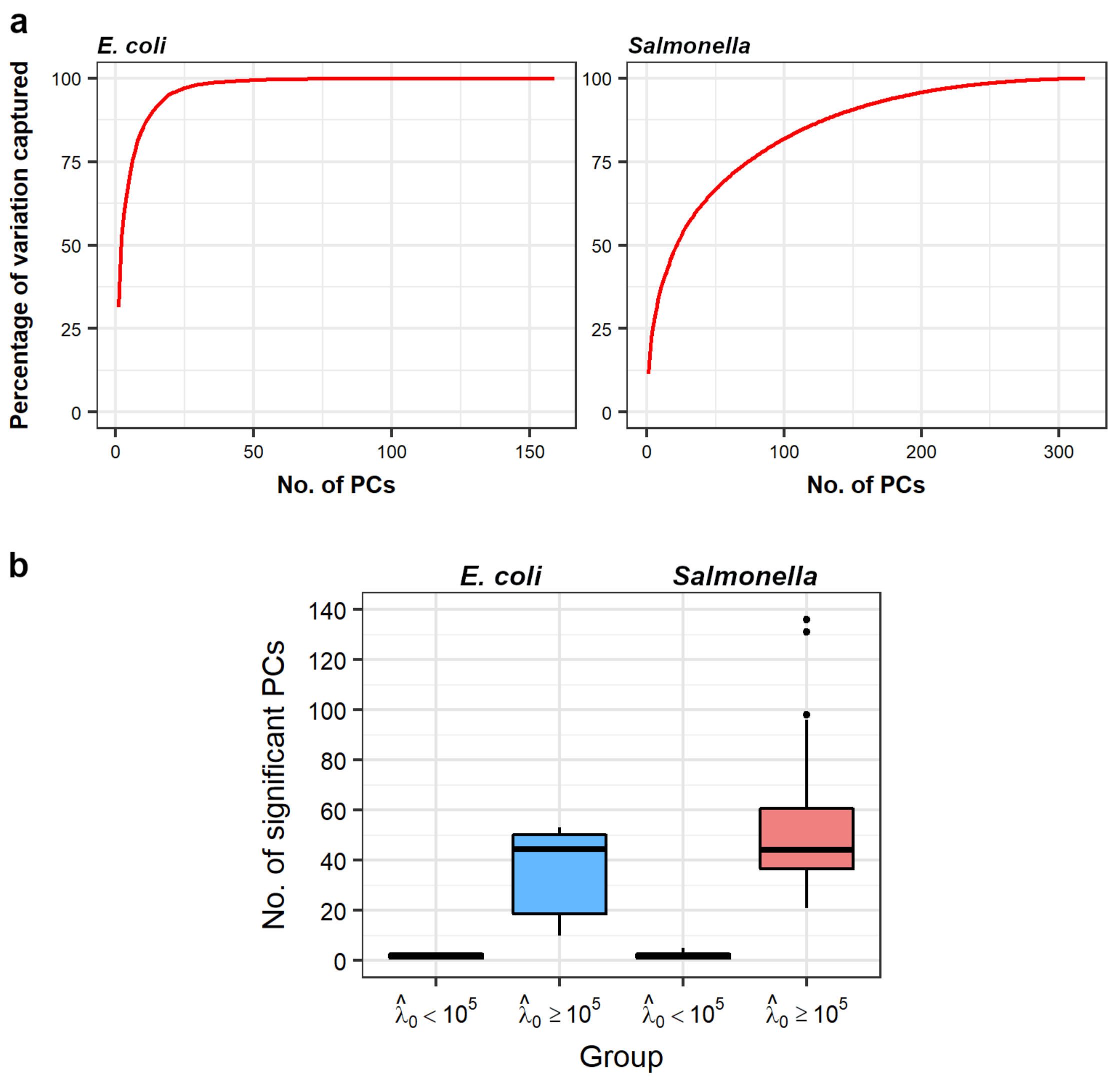

**Figure s8: A summary of PCs obtained from core-genome relatedness matrices**. (**a**) Cumulative percentage of total genetic variation captured by PCs of each species. According to Equation 23 in Section 3.1.7, when PCs are arranged in a descending order of their corresponding eigenvalues, the percentage of genetic variations captured by the first $k$ PCs is calculated by the formula $\sum_{i=1}^{k} \lambda_i / \sum_{i=1}^{n} \lambda_i \times 100\%$, where $\lambda_i$ is the $i$-th PC and $n$ is the total number of PCs. Specifically, all genetic variation of the 169 *E. coli* genomes are captured by 159 PCs, where the first PC, first six PCs and the first 20 PCs capture 31.36%, $>75\%$ and $>95\%$ of total variation, respectively; total genetic variation of the 359 *Salmonella* genomes are captured by 319 PCs, where the first PC, first 74 PCs and the first 191 PCs capture 11.29%, $>75\%$ and $>95\%$ of the total variation, respectively. (**b**) Number of significant PCs contributing to presence-absence of the response pattern in an LMM. Significant PCs are determined based on a maximum of 0.05 for Bonferroni-corrected p-values. For each species, the number of PCs per response is grouped by the REML estimate of $\lambda_0$. Considering both species, for patterns whose $\hat{\lambda}_0 < 10^5$, the count of significant PCs varies between one and five, while for patterns whose $\hat{\lambda}_0 \geq 10^5$, this count varies between 10 and 136.

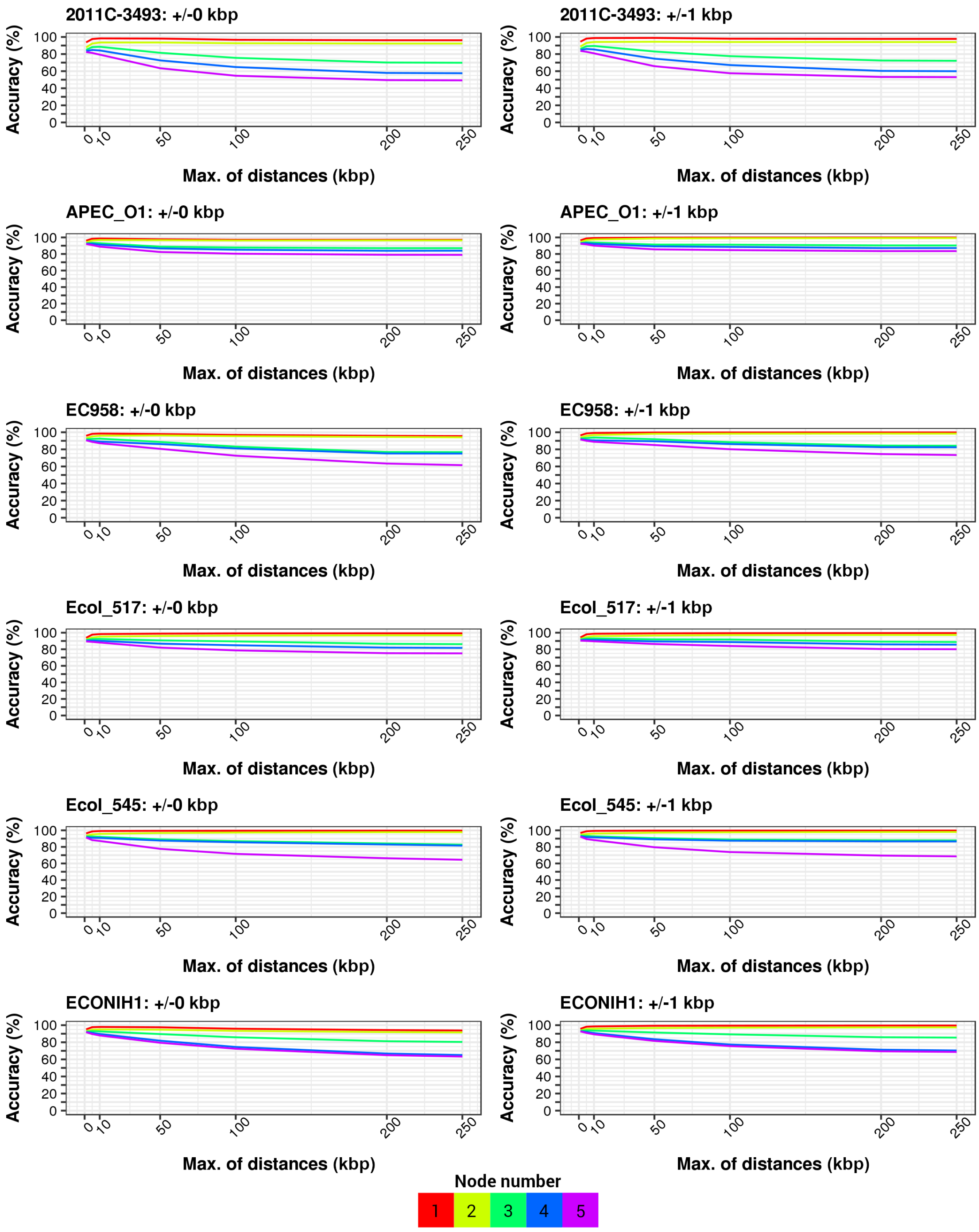

**Figure s9: Accuracy of SPDs measured across at most five nodes in assembly graphs of *E. coli* genomes under two levels of error tolerance (0 or 1 kbp)**. The assembly graph was generated from imperfect simulated reads. A genome name and error tolerance are printed in each panel. Only accuracy rates for six out of the ten genomes are shown in this figure for clarity. APD: allelic physical distance.

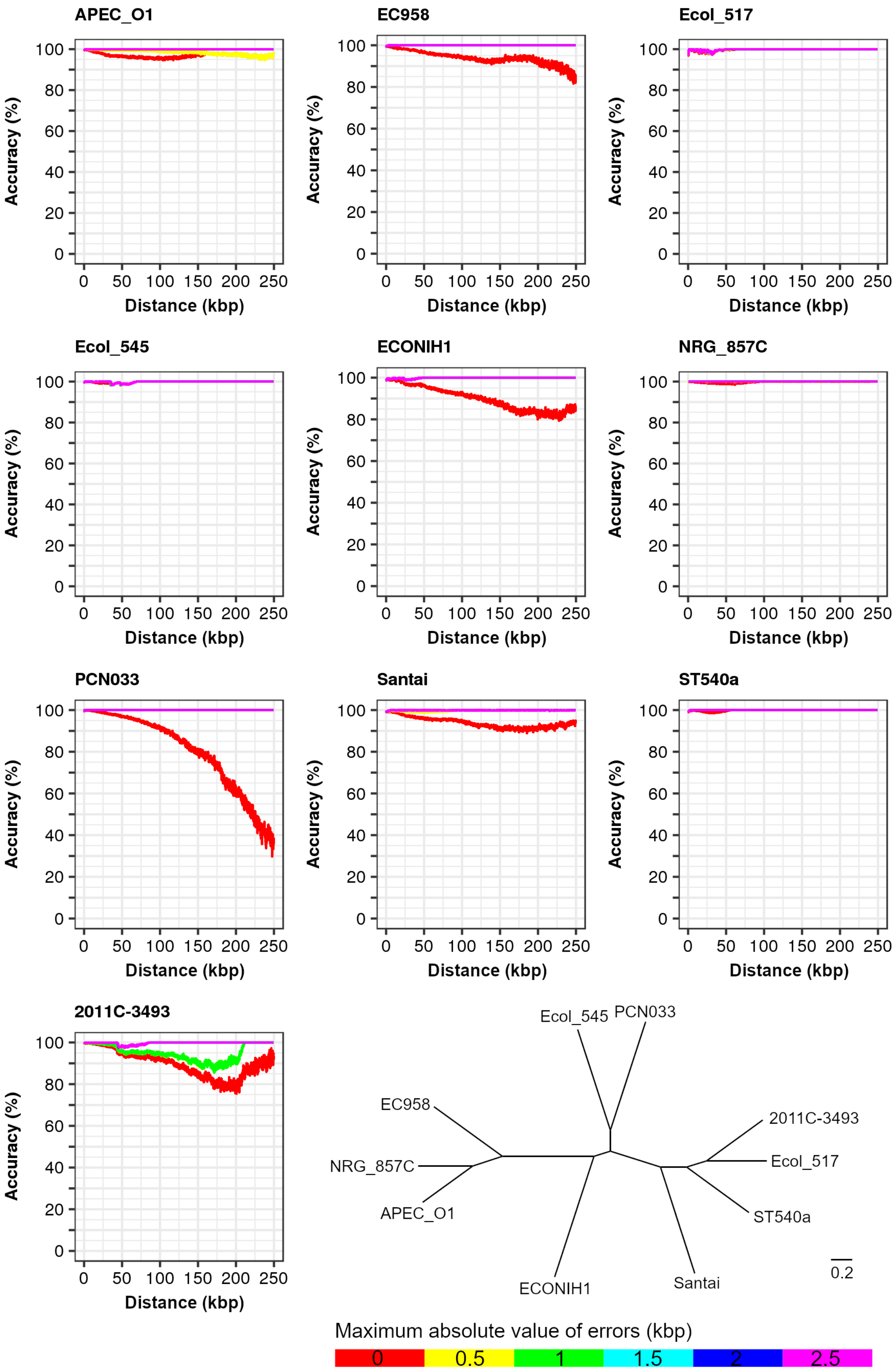

**Figure s10: Accuracy of SPDs in contigs under six error tolerance levels for 10 *E. coli* genomes**. A genome name is shown at the top of each panel. Distance: the SPD between two alleles in a contig. Accuracy: percentage of SPDs differing from true distances by no more than a given error tolerance level. The neighbour-joining tree shows the mean whole-genome average nucleotide identity (ANI) calculated with FastANI v1.0 for each pair of genomes [2].

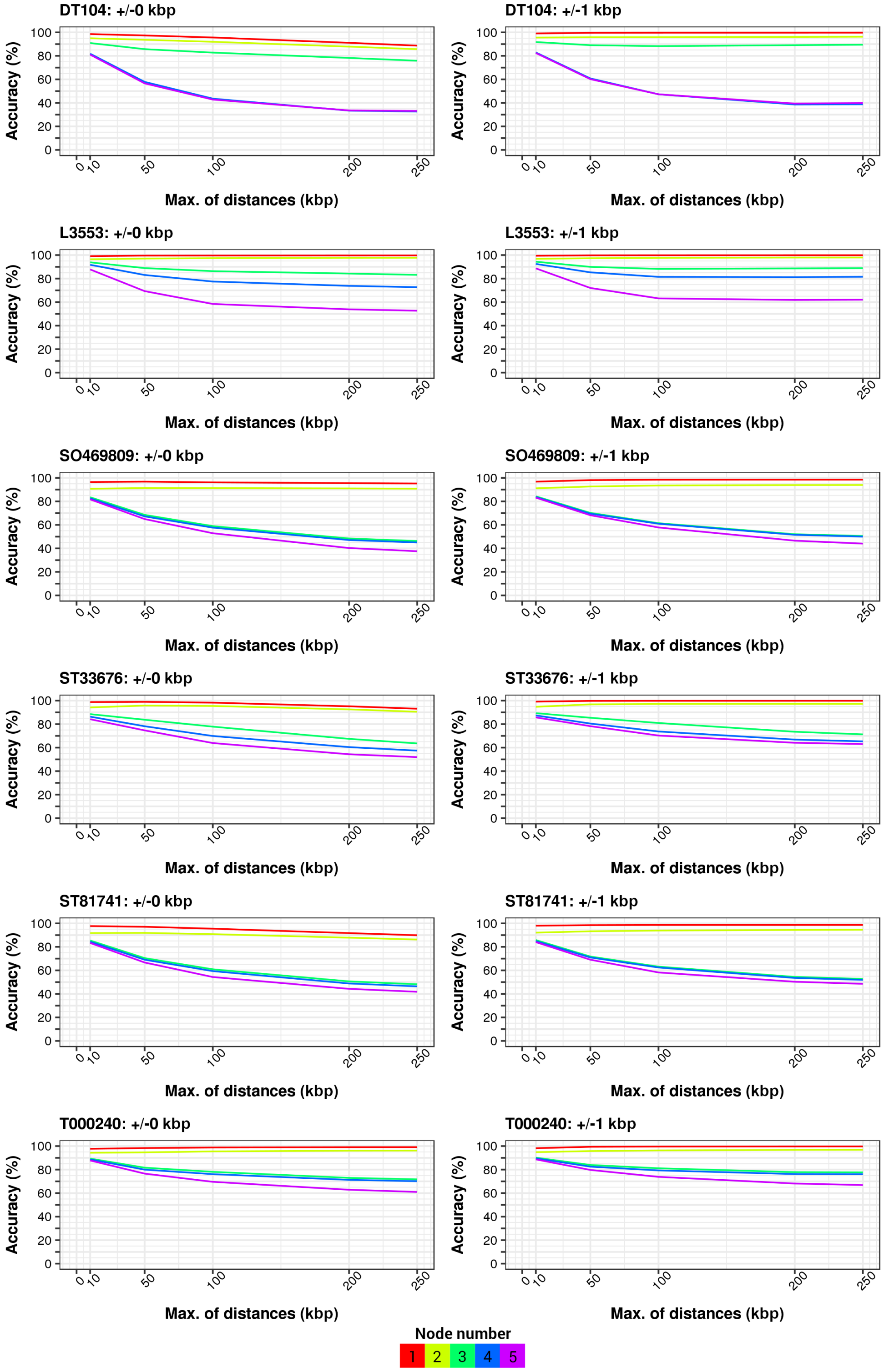

**Figure s11: Accuracy of SPDs measured across at most five nodes in assembly graphs of *Salmonella* genomes when tolerating no or at most 1 kbp errors**. The assemblies were generated from imperfect simulated reads. A genome name and error tolerance are printed in each panel. Only results for six out of the ten genomes are shown here for clarity.

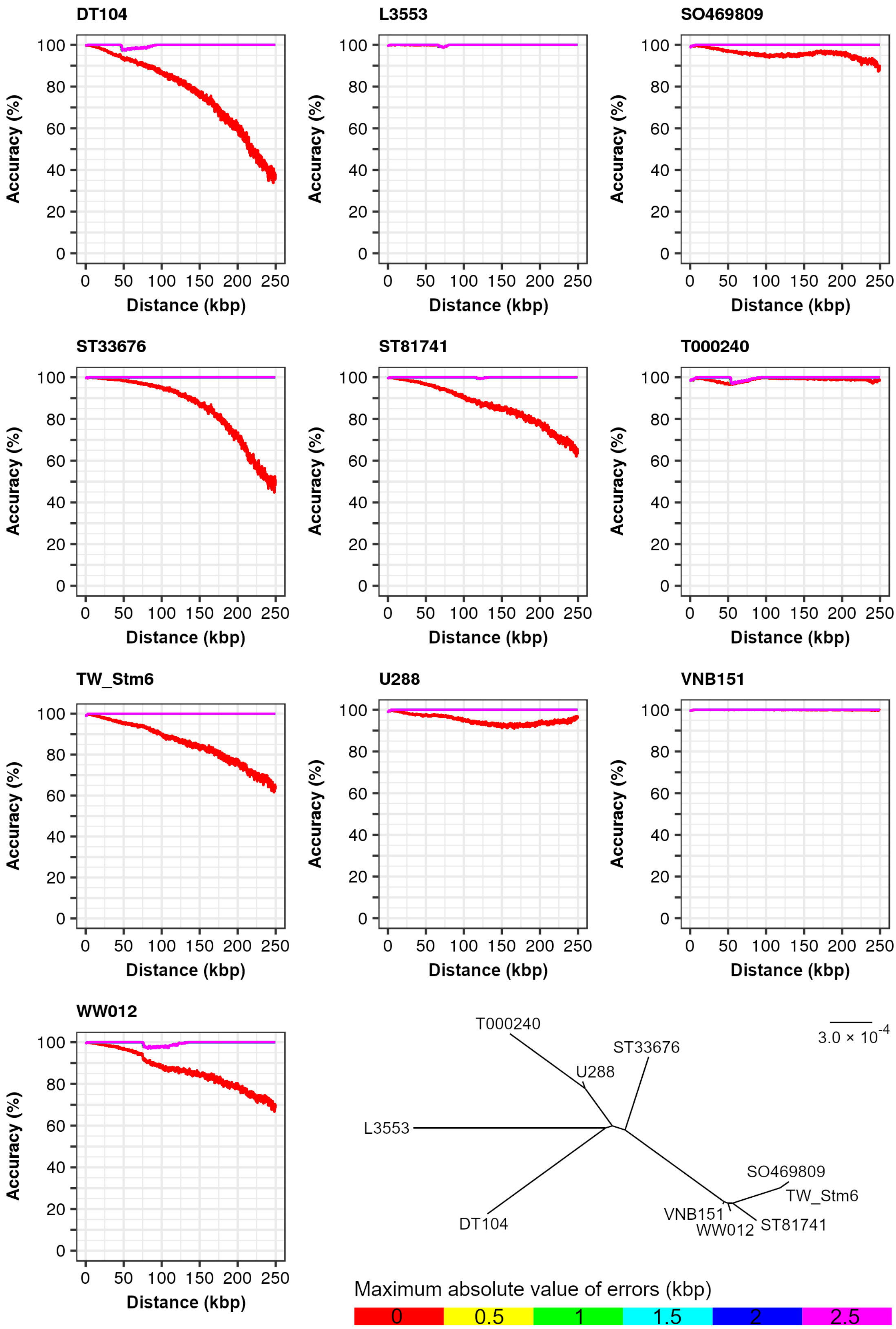

**Figure s12: Accuracy of SPDs measured in contigs under different levels of error tolerance for 10 *S.* Typhimurium genomes**. A genome name is printed to the top of each panel. We analysed all distances without a random sampling because the number of measurable distances in contigs is much smaller than that in assembly graphs. Distance: the shortest distance between two loci in an assembly graph. The neighbour-joining tree shows the mean whole-genome ANI calculated with FastANI v1.0 for each pair of genomes.

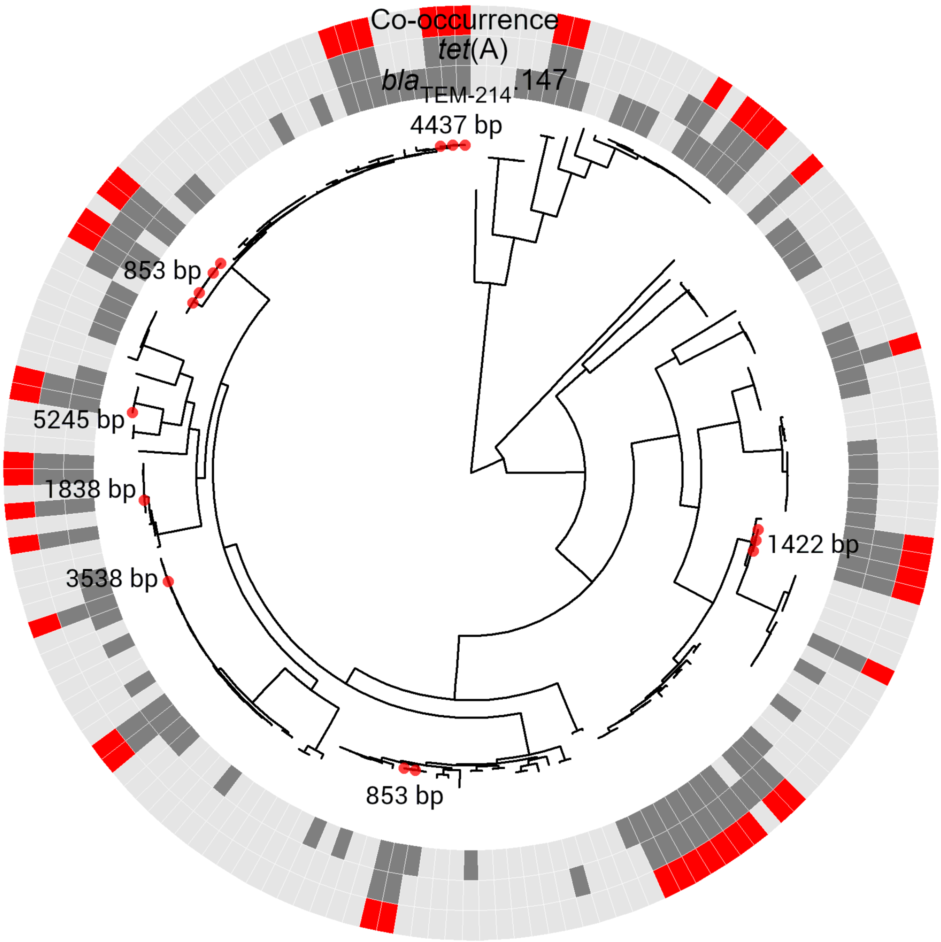

**Figure s13: Distribution of six reliable SPDs measured between positively associated alleles *bla*$_{\text{TEM-214}}$.147 and *tet*(A) in 15 *E. coli* genomes**. Genome assemblies in which reliable SPDs were obtained are highlighted as red circles on tips of the midpoint-rooted ML phylogeny, and are labelled by the distance values. Abbreviation: Co, co-occurrence of the two alleles.

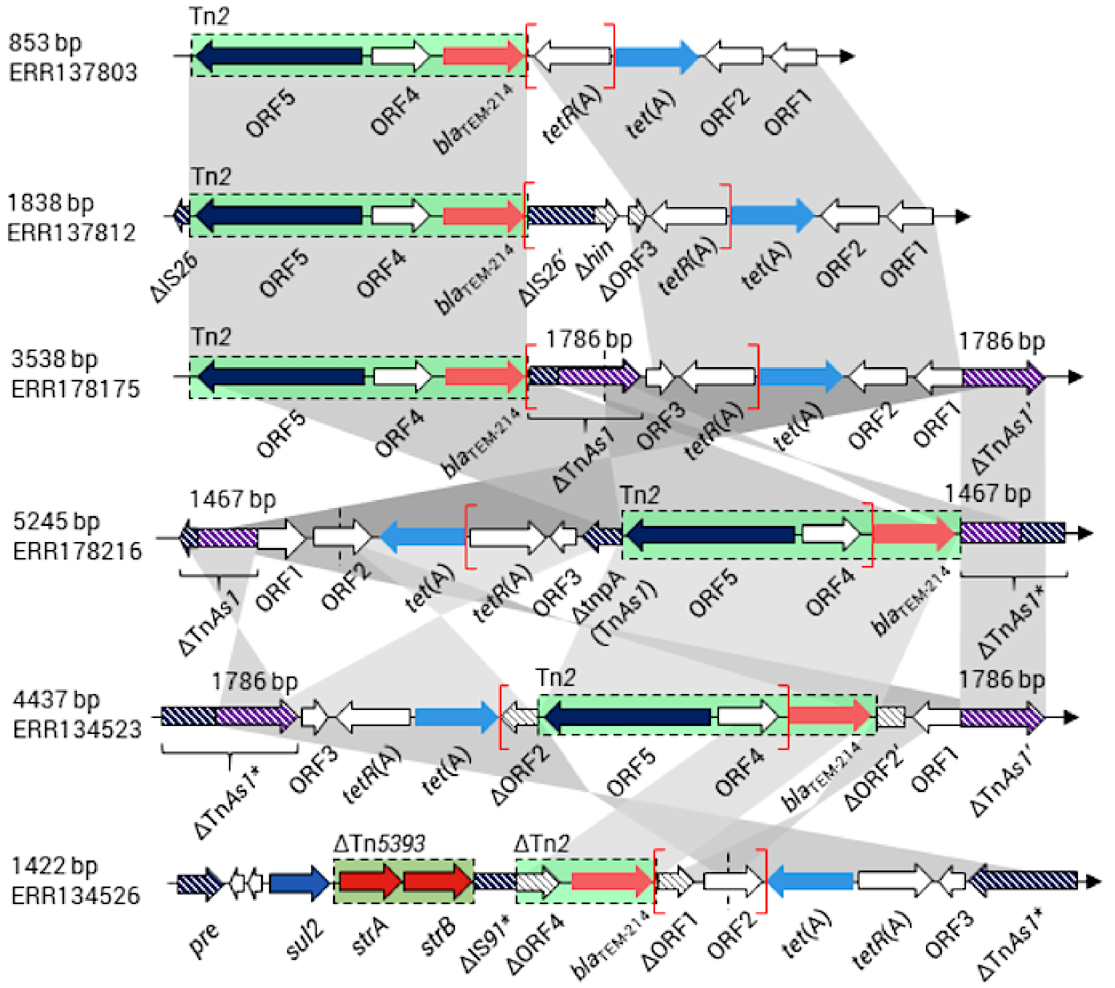

**Figure s14: Resolved structures of regions comprising alleles *bla*$_{TEM-214}$.147 and *tet*(A) in *E. coli* genomes**. The alleles are denoted by their gene names in this figure. The SPD between these two alleles and the genome name are displayed on the left of each structure. The distribution of SPDs in *E. coli* genomes is illustrated in Figure s13. We annotated these regions via searching their nucleotide sequences against *Enterobacteriaceae* genomes in GenBank with megaBLAST. For each structure, a pair of red brackets show the region in which the SPD was measured. Grey shades between structures indicate homologous regions showing $> 99\%$ nucleotide identity. Green boxes with dashed borders represent two transposon sequences. Arrows and boxes filled with colour patterns denote pseudo genes or MGEs. All the six genetic structures contained a 4,950 bp Tn*3*-family transposon Tn*2* (GenBank accession: KT002541, coordinates: 1–4,917), either complete or truncated (1,573 bp, in a single genome), showing a 100% nucleotide identity to each other. Repeats of partial Tn*As1* sequences are highlighted using a purple-white filling pattern and had their lengths labelled nearby. The sign Δ denotes a truncated gene or a genetic element. The asterisks besides an MGE name indicates a variant of the corresponding MGE. Annotations for open reading frames (ORFs): ORF1, cysteine hydrolase (NCBI protein ID: AWA37038) gene; ORF2, a gene encoding an *EamA* family transporter (NCBI protein ID: AYD32134); ORF3, a 243 bp relaxase (NCBI protein ID: AXE60424) gene, which is associated with insertion sequences and transposons; ORF4, a gene encoding a recombinase-family protein (NCBI protein ID: AXS38585); ORF5, a gene encoding a Tn*3*-family transposase (NCBI protein ID: AXS38584). Δ*tnpA*: the 2,964 bp transposase gene of Tn*As1* (GenBank accession: CP022426, at coordinates 4,991,027–4,993,990); Δ*hin*, a 174 bp truncated gene encoding a DNA-invertase *Hin* (GenBank accession: MG692690, at coordinates 2,971–3,387).

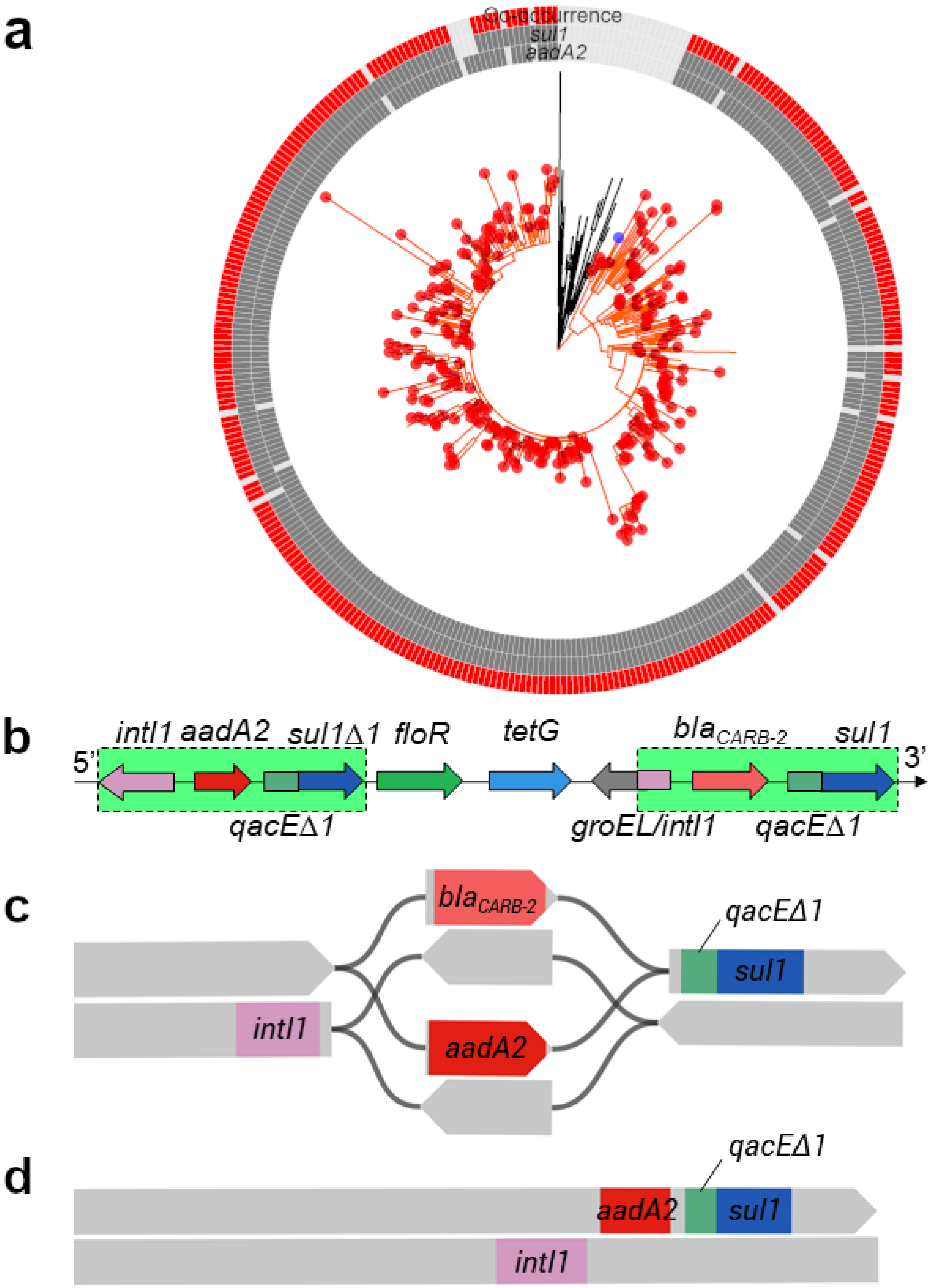

**Figure s15: Reconstructed genetic structures for alleles *sul1* and *aadA2* in *Salmonella* genomes**. (**a**) Distribution of the alleles in 359 *Salmonella* genomes. In the midpoint-rooted ML phylogenetic tree shown in the centre, red and blue circles highlight tips representing genomes from which SPDs between the two alleles were obtained. Particularly, the blue circle denotes the strain DT104, whose complete genome is available in GenBank. A single lineage from which all SPDs were obtained is coloured in orange. (**b**) A diagram showing genetic structure of the MDR region, which was created based on Figure 2 by Boyd, et al. [3]. Other genes within this region are omitted for simplicity. (**c**) An assembly graph (genome DRR006262) of double DNA strands in which the SPD between the alleles was 504 bp. (**d**) Double DNA strands of a single contig (genome ERR170653) harbouring both alleles, which were 504 bp apart in the contig.

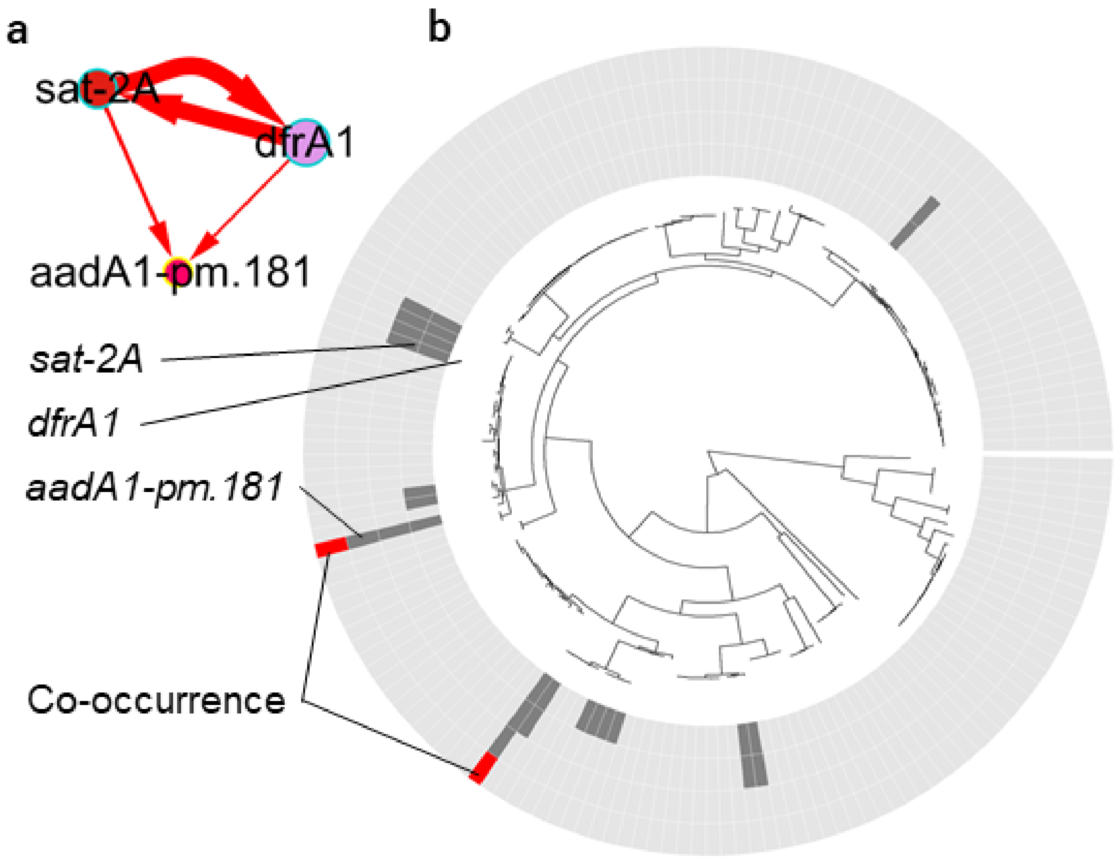

**Figure s16: A maximal clique of three alleles of AMR genes extracted from the linkage network of *E. coli* and distribution of its alleles in genomes**. (**a**) Presence-absence of these alleles were positively associated, as determined using LMMs, and SPDs between these alleles were always measurable and consistent. (**b**) A ring plot created for this clique using GeneMates, which illustrates co-occurrence of all three alleles (red tiles in the outer most ring) and presence-absence of individual alleles (tiles coloured in dark grey in inner rings). A midpoint-rooted phylogeny of *E. coli* genomes is shown at the middle.

### 2 Supplementary tables

**Table s1: A summary of 10 MDR *E. coli* genomes used for determining reliability criteria for APDs**. An accession number in the NCBI nucleotide database is provided for each nucleotide sequence. The sequence length is measured in base pairs (bp). Abbreviations: AMR, antimicrobial resistance; NA, not detected.

| Strain | Sequence | Accession | Length | AMR genes |
|---|---|---|---|---|
| 2011C-3493 | chromosome | CP003289 | 5,273,097 | *ampH*, *ampC1*, *ampC2*, *mrdA*, *drfA7*, *strA*, *strB*, *sul1*, *sul2*, *tet*(A) |
| | plasmid | CP003291 | 74,217 | NA |
| | plasmid | CP003290 | 88,544 | $bla_{\text{CTX-M-15}}$, $bla_{\text{TEM-105}}$ |
| | plasmid | CP003292 | 1,549 | NA |
| NRG_857C | chromosome | CP001855 | 4,747,819 | *ampH*, *ampC2*, *mrdA* |
| | plasmid | CP001856 | 147,060 | *aadA1-pm*, *catA1*, *drfA1*, *mphB*, *strA*, *strB*, *sul1*, *sul2*, *tet*(A), $bla_{\text{TEM-105}}$ |
| Ecol_517 | chromosome | CP018965 | 4,794,957 | *ampH*, *ampC1*, *ampC2*, *mrdA* |
| | plasmid | CP018964 | 118,495 | *aac6-Ib*, *aadA5*, $bla_{\text{CTX-M-15}}$, *catB4*, *dfrA7*, *mphA*, $bla_{\text{OXA-1}}$, *sul1*, *tet*(A) |
| | plasmid | CP018963 | 54,644 | $bla_{\text{KPC-2}}$ |
| Ecol_545 | chromosome | CP018976 | 5,031,843 | *ampH*, *ampC1*, *ampC2*, *mrdA*, $bla_{\text{CTX-M-15}}$, *qnr-S1* |
| | plasmid | CP018975 | 95,926 | NA |
| | plasmid | CP018974 | 70,876 | *aac6-Ib*, *catB3*, $bla_{\text{KPC-2}}$, $bla_{\text{OXA-1}}$, $bla_{\text{TEM-105}}$ |
| | plasmid | CP018973 | 70,152 | $bla_{\text{CTX-M-27}}$ |
| | plasmid | CP018972 | 4,073 | NA |
| | plasmid | CP018971 | 3,164 | NA |
| APEC_O1 | chromosome | CP000468 | 5,082,025 | *ampH*, *ampC1*, *ampC2*, *mrdA* |
| | plasmid | DQ381420 | 174,241 | NA |
| | plasmid | DQ517526 | 241,387 | *aac3-VIa*, *aadA1-pm*, *sul1*, *tet*(C) |
| PCN033 | chromosome | CP006632 | 4,987,957 | *ampH*, *ampC2*, *mrdA*, *aac3-IId*, $bla_{\text{TEM-105}}$ |
| | plasmid | CP006633 | 3,319 | NA |
| | plasmid | CP006634 | 4,086 | NA |
| | plasmid | CP006635 | 161,511 | *aph3-Ia*, *dfrA17*, *oqxA*, *oqxB*, *strA*, *strB*, *sul2*, $bla_{\text{TEM-105}}$, *tet*(B) |
| ST540a | chromosome | CP007390 | 4,807,977 | *ampH*, *ampC1*, *ampC2*, *mrdA*, *aph3-Ia*, *strA*, *strB*, *sul2*, $bla_{\text{TEM-150}}$, *tet*(A) |

| | | | | |
|---|---|---|---|---|
| Santai | chromosome | NZ_CP007592 | 5,104,557 | *ampH*, *ampC1*, *ampC2*, *mrdA*, *aac3-IId*, *aac6-Ib*, *aadA2*, *armA*, *arr3*, *catA1*, *catB3*, *dfrA12*, *floR*, *fosA*, *mphA*, *mphE*, *msrE*, *bla*$_{\text{OXA-1}}$, *strA*, *strB*, *sul1*, *sul2*, *bla*$_{\text{TEM-105}}$, *tet*(A) |
| ECONIH1 | chromosome | CP009859 | 5,310,511 | *ampH*, *ampC1*, *ampC2*, *mrdA*, *bla*$_{\text{CTX-M-15}}$ |
| | plasmid | CP009860 | 121,385 | *aadA5*, *dfrA17*, *ermB*, *mphA*, *sul1* |
| | plasmid | CP009861 | 47,560 | NA |
| | plasmid | CP009862 | 80,186 | *aac6-Ib*, *aadA1-pm*, *dfrA14*, *bla*$_{\text{KPC-2}}$, *bla*$_{\text{OXA-9}}$, *strA*, *strB*, *sul2*, *bla*$_{\text{TEM-150}}$ |
| EC958 | chromosome | HG941718 | 5,109,767 | *ampH*, *ampC2*, *mrdA*, *bla*$_{\text{CMY-23}}$ |
| | plasmid | HG941719 | 135,602 | *aac6-Ib*, *aadA5*, *bla*$_{\text{CTX-M-15}}$, *catB4*, *dfrA17*, *mphA*, *bla*$_{\text{OXA-1}}$, *sul1*, *bla*$_{\text{TEM-105}}$, *tet*(A) |
| | plasmid | HG941720 | 4,080 | NA |

**Table s2: A summary of the 10 MDR *S*. Typhimurium genomes used for determining reliability criteria for APDs**. An accession number in the NCBI nucleotide database is provided for each nucleotide sequence. The sequence length is measured in base pairs (bp). Abbreviations: AMR, antimicrobial resistance; NA, not detected.

| Strain | Sequence | Accession | Length | AMR genes |
|---|---|---|---|---|
| DT104 | chromosome | HF937208 | 4,933,631 | *aac6-Iaa*, *aadA2*, *bla*$_{\text{CARB-2}}$, *floR*, *sul1*, *tet*(G) |
| | plasmid | HF937209 | 94,034 | NA |
| TW_Stm6 | chromosome | CP019649 | 4,999,862 | *aac6-Iaa*, *strA*, *strB*, *sul2*, *bla*$_{\text{TEM-105}}$, *tet*(B) |
| | plasmid | CP019647 | 275,801 | *aadA2*, *aadA1-pm*, *aphA2*, *cmlA1*, *dfrA12*, *strA*, *strB*, *sul3*, *bla*$_{\text{TEM-105}}$, *tet*(A) |
| | plasmid | CP019648 | 4,083 | NA |
| ST33676 | chromosome | CP012681 | 4,809,574 | *aac6-Iaa* |
| | plasmid | CP012683 | 112,639 | *cmy-17* |
| | plasmid | CP012684 | 4,512 | NA |
| | plasmid | CP012682 | 161,461 | *aac3-IId*, *aadA2*, *dfrA12*, *floR*, *oqxA*, *oqxB*, *strA*, *strB*, *sul2*, *sul3*, *tet*(A) |
| T000240 | chromosome | AP011957 | 4,954,814 | *aac6-Iaa*, *aadA1-pm*, *catA1*, *bla*$_{\text{OXA-1}}$, *sul1*, *tet*(B) |
| | plasmid | AP011958 | 106,510 | *aac3-IId*, *aadA2*, *dfrA12*, *sul1* |
| | plasmid | AP011959 | 8,670 | *strA*, *strB*, *sul2* |
| | chromosome | CP003836 | 4,852,606 | *aac6-Iaa* |

U288

| | | | | |
|---|---|---|---|---|
| | plasmid | CP004058 | 148,711 | *aadA2*, *aadA1-pm*, *cmlA1*, *dfrA12*, *sul3*, $bla_{\text{TEM-105}}$ |
| | plasmid | CP004059 | 11,067 | *strA*, *strB*, *sul2*, *tet*(A) |
| | plasmid | CP004060 | 4,675 | NA |
| ST81741 | chromosome | CP019442 | 4,974,856 | *aac6-Iaa*, *tet*(B) |
| | plasmid | CP019443 | 233,802 | *aac3-IId*, *aadA17*, $bla_{\text{CTX-M-65}}$, *floR*, *lunF*, *sul2*, $bla_{\text{TEM-105}}$, *tet*(M) |
| | plasmid | CP019444 | 84,565 | *mphA*, $bla_{\text{NDM-5}}$, $bla_{\text{TEM-105}}$ |
| L3553 | chromosome | AP014565 | 5,051,841 | *aac6-Iaa*, *aada2*, *cmy-17*, *dfrA12*, *floR*, *strA*, *strB*, *sul1*, *sul2*, *tet*(A) |
| | plasmid | AP014566 | 132,611 | *aph3-Ia*, *sul1*, $bla_{\text{TEM-105}}$, *tet*(A) |
| SO469809 | chromosome | NZ_LN999997 | 5,037,238 | *aac6-Iaa*, *strA*, *strB*, *sul2*, $bla_{\text{TEM-105}}$, *tet*(B) |
| WW012 | chromosome | NZ_CP022168 | 4,991,167 | *aac6-Iaa*, *strA*, *strB*, *sul2*, *tet*(B) |
| | plasmid | NZ_CP022169 | 151,609 | *aadA2*, *aadA1-pm*, *cmlA1*, *dfrA12*, *mcr-1*, *sul3* |
| VNB151 | chromosome | NZ_LT795114 | 4,985,374 | *aac6-Iaa*, *tet*(B) |
| | plasmid | NZ_LT795115 | 246,444 | *aac3-Iva*, *aac6-Ib*, *aadA2*, *aadA1-pm*, *aph3-Ia*, *aph4-Ia*, *arr3*, *catB3*, *cmlA1*, *floR*, $bla_{\text{OXA-1}}$, *oqxA*, *oqxB*, *sul1*, *sul2*, *sul3* |
| | plasmid | NZ_LT795116 | 4,239 | NA |

**Table s3: Number of queries unidentified in every assembly graph and contig file of *E. coli* genomes**. Each query is a random CDS extracted from a complete genome for the distance measurement. Bandage runs the nucleotide BLAST to locate queries in each file so as to measure the physical distances. For specificity of our analyses, we accepted the hit that covered at least 95% of a query path under a minimum nucleotide identity 95% and a maximum e-value $1 \times 10^{-5}$.

| Strain | No. of queries | Missing hits in contig file | Missing hits in graph file |
|---|---|---|---|
| 2011C-3493 | 5,150 | 229 | 43 |
| NRG_857C | 4,582 | 40 | 7 |
| Ecol_517 | 4,932 | 103 | 37 |
| Ecol_545 | 5,214 | 148 | 28 |
| APEC_O1 | 4,891 | 164 | 29 |
| PCN033 | 5,076 | 74 | 20 |
| ST540a | 4,562 | 121 | 19 |
| Santai | 4,838 | 73 | 14 |
| ECONIH1 | 5,322 | 143 | 18 |
| EC958 | 5,100 | 113 | 25 |

**Table s4: Number of queries unidentified in every assembly graph and contig file of DT104 genomes**. Each query is a random CDS extracted from a complete genome for the distance measurement. Bandage runs the nucleotide BLAST to locate queries in each file so as to measure the physical distances. For specificity, we accepted the hit that covered at least 95% of a query path under a minimal nucleotide identity of 95% and a maximum e-value of $1 \times 10^{-5}$.

| Strain | No. of queries | Missing hits in contig file | Missing hits in graph file |
|---|---|---|---|
| DT104 | 4,656 | 68 | 9 |
| TW_Stm6 | 5,062 | 110 | 13 |
| ST33676 | 4,767 | 58 | 11 |
| T000240 | 4,871 | 78 | 19 |
| U288 | 4,798 | 58 | 19 |
| ST81741 | 5,172 | 85 | 18 |
| L3553 | 5,106 | 69 | 15 |
| SO469809 | 4,950 | 105 | 11 |
| WW012 | 5,009 | 68 | 12 |
| VNB151 | 5,133 | 79 | 17 |

**Table s5: Accuracy of prioritised SPDs measured between alleles of accessory AMR genes in contigs and assembly graphs of *E. coli***. Since there may be $\geq 2$ copies of an allele at different loci in a genome, the reference distance to be compared with between two alleles was defined as the shortest one among all distances. The actual absolute value of errors given $\leq 2$ nodes did not exceed 19 bp. $N_i$ ($i = 1, .., 5$): accuracy, number of accurate/all distances given $\leq i$ nodes.

| Strain | $N_1$ | $N_2$ | $N_3$ | $N_4$ | $N_5$ |
|---|---|---|---|---|---|
| 2011C-3493 | 100% (5/5) | 100% (5/5) | 100% (5/5) | 100% (5/5) | 75.00% (6/8) |
| APEC_O1 | 100% (6/6) | 100% (6/6) | 100% (6/6) | 100% (6/6) | 100% (6/6) |
| EC958 | 100% (9/9) | 100% (9/9) | 81.82% (9/11) | 46.15% (12/26) | 46.15% (12/26) |
| Ecol_517 | 100% (9/9) | 100% (9/9) | 39.29% (11/28) | 39.29% (11/28) | 36.11% (13/36) |
| Ecol_545 | 100% (3/3) | 100% (4/4) | 100% (4/4) | 100% (5/5) | 100% (5/5) |
| ECONIH1 | 100% (10/10) | 93.33% (14/15) | 90.00% (18/20) | 90.00% (18/20) | 83.33% (25/30) |
| NRG_857C | 100% (16/16) | 100% (16/16) | 100% (22/22) | 91.67% (22/24) | 91.67% (22/24) |
| PCN033 | 100% (5/5) | 100% (5/5) | 33.33% (11/33) | 33.33% (11/33) | 27.50% (11/40) |
| Santai | 100% (23/23) | 100% (41/41) | 86.54% (45/52) | 86.54% (45/52) | 60.49% (49/81) |
| ST540a | 100% (3/3) | 100% (3/3) | 100% (3/3) | 100% (3/3) | 57.14% (4/7) |
| Margin | 100% (89/89) | 99.12% (112/113) | 72.83% (134/184) | 68.32% (138/202) | 58.17% (153/263) |

**Table s6: Accuracy of prioritised SPDs measured between alleles of accessory AMR genes in contigs and assembly graphs of DT104 genomes**. We filtered BLAST hits for a nucleotide identity and query coverage of **95%**. The distances were prioritised based on our empirical discovery that the distance measurements were more accurate in contigs than were in assembly graphs. Since there may be $\geq 2$ copies of an allele in different genomic loci of a genome, the reference distance to be compared with between two alleles was defined as the shortest one among all distances for this table. Error tolerance: $\pm 1$ kb; $N_i$ ($i = 1, .., 5$): accuracy, number of accurate/all distances under a node number $\leq i$.

| Strain | $N_1$ | $N_2$ | $N_3$ | $N_4$ | $N_5$ |
|---|---|---|---|---|---|
| DT104 | 100% (1/1) | 66.67% (2/3) | 71.43% (5/7) | 77.78% (7/9) | 80% (8/10) |
| L3553 | 100% (6/6) | 100% (16/16) | 92.00% (23/25) | 80% (24/30) | 72.73% (24/33) |
| SO469809 | 100% (3/3) | 100% (3/3) | 70.00% (7/10) | 70% (7/10) | 70% (7/10) |
| ST33676 | 100% (12/12) | 100% (16/16) | 59.26% (16/27) | 47.06% (16/34) | 38.10% (16/42) |
| ST81741 | 100% (3/3) | 100% (5/5) | 35.71% (10/28) | 38.71% (12/31) | 34.15% (14/41) |
| T000240 | 100% (10/10) | 100% (14/14) | 100% (16/16) | 100% (16/16) | 100% (16/16) |
| TW_Stm6 | 81.82% (9/11) | 77.78% (14/18) | 70% (14/20) | 57.14% (24/42) | 48.28% (28/58) |
| U288 | 100% (16/16) | 100% (16/16) | 76.19% (16/21) | 76.19% (16/21) | 76.19% (16/21) |
| VNB151 | 100% (36/36) | 100% (42/42) | 62.50% (60/96) | 56.14% (64/114) | 48.48% (64/132) |
| WW012 | 88.89% (8/9) | 88.89% (8/9) | 56.25% (9/16) | 63.16% (12/19) | 60% (12/20) |
| **Margin** | 97.20% (104/107) | 95.77% (136/142) | 66.17% (176/266) | 60.74% (198/326) | 53.52% (205/383) |

**Table s7: Accuracy of prioritised SPDs measured between alleles of accessory AMR genes in contigs and assembly graphs of DT104 genomes**. We filtered BLAST hits for a nucleotide identity and query coverage of **99%**. The distances were prioritised based on our empirical discovery that the distance measurements were more accurate in contigs than were in assembly graphs. Since there may be $\geq 2$ copies of an allele in different genomic loci of a genome, the reference distance to be compared with between two alleles was defined as the shortest one among all distances for this table. Error tolerance: $\pm 1$ kb; $N_i$ ($i = 1, .., 5$): accuracy, number of accurate/all distances under a node number $\leq i$.

| Strain | $N_1$ | $N_2$ | $N_3$ | $N_4$ | $N_5$ |
|---|---|---|---|---|---|
| DT104 | 100% (1/1) | 66.67% (2/3) | 71.43% (5/7) | 77.78% (7/9) | 80% (8/10) |
| L3553 | 100% (6/6) | 100% (16/16) | 92.00% (23/25) | 80% (24/30) | 72.73% (24/33) |
| SO469809 | 100% (3/3) | 100% (3/3) | 70.00% (7/10) | 70% (7/10) | 70% (7/10) |
| ST33676 | 100% (12/12) | 100% (16/16) | 59.26% (16/27) | 47.06% (16/34) | 38.10% (16/42) |
| ST81741 | 100% (3/3) | 100% (5/5) | 35.71% (10/28) | 38.71% (12/31) | 34.15% (14/41) |
| T000240 | 100% (10/10) | 100% (14/14) | 100% (16/16) | 100% (16/16) | 100% (16/16) |
| TW_Stm6 | 100% (8/8) | 100% (11/11) | 84.62% (11/13) | 60% (21/35) | 58.14% (25/43) |
| U288 | 100% (16/16) | 100% (16/16) | 76.19% (16/21) | 76.19% (16/21) | 76.19% (16/21) |
| VNB151 | 100% (36/36) | 100% (42/42) | 62.50% (60/96) | 56.14% (64/114) | 48.48% (64/132) |
| WW012 | 100% (6/6) | 100% (6/6) | 53.85% (7/13) | 60% (9/15) | 60% (9/15) |
| **Margin** | 100% (101/101) | 99.24% (131/132) | 66.80% (171/256) | 60.95% (192/315) | 54.82% (199/363) |

**Table s8: SPDs measured between five alleles of SGI1-borne AMR genes in *Salmonella*.** SPDs were measured in complete genomes, contigs and assembly graphs. Rows are sorted by values of $N_r$ in a descending order. Column names: LMM, whether an LMM-based significant association is identified between two alleles (●, yes; ○, no); $w_d$, the weighted distance score; N, number of all SPDs; SPD, range of all SPDs; $N_{node}$, numbers of nodes across which the SPDs were measured; $N_r$, number of reliable SPDs; $SPD_r$, the range of reliable SPDs. For each pair of alleles, the percentage of reliable SPDs is calculated by the formula $N_r/N \times 100\%$.

| Allele pair | LMM | $w_d$ | N | SPD (bp) | $N_{node}$ | $N_r$ | $SPD_r$ (bp) |
|---|---|---|---|---|---|---|---|
| *aadA2*, *sul1* | ● | 0 | 295 | 504–9,964 | 1–3 | 294 | 504–9,964 |
| $bla_{CARB-2}$, *sul1* | ● | 0.92 | 266 | 557–557 | 1–3 | 265 | 557–557 |
| *floR*.12, *tet*(G) | ● | 0.99 | 202 | 937–937 | 1–1 | 202 | 937–937 |
| $bla_{CARB-2}$, *tet*(G) | ● | 0.04 | 258 | 3,521–4,673 | 1–7 | 13 | 3,521–3,896 |
| *aadA2*, *tet*(G) | ● | 0.04 | 254 | 3,473–4,620 | 1–7 | 11 | 3,473–3,843 |
| $bla_{CARB-2}$, *floR*.12 | ● | 0.04 | 186 | 1,745–5,634 | 1–3 | 10 | 1,745–5,634 |
| *sul1*, *tet*(G) | ● | 0.03 | 250 | 3,185–710,475 | 1–44 | 8 | 4,757–4,945 |
| *floR*.12, *sul1* | ○ | 0.03 | 181 | 6,301–7,058 | 1–13 | 7 | 6,870–7,058 |
| *aadA2*, *floR*.12 | ● | 0.02 | 181 | 1,692–5,586 | 1–3 | 6 | 1,692–5,586 |
| *aadA2*, $bla_{CARB-2}$ | ○ | 0 | 249 | 5,221–704,968 | 1–57 | 1 | 8,540–8,540 |

**Table s9: Physical distances measured between 45 pairs of positively associated alleles in 169 *E. coli* genomes**. A minimum of two reliable SPDs were obtained for each pair. Abbreviations of column names: Co, co-occurrence count; M, measurability of all SPDs; $M_r$, measurability of reliable SPDs; $P_r$, $M_r/M \times 100\%$ – percentage of reliable SPDs in all the SPDs; $P_{nongraph}$, percentage of reliable SPDs not measured in assembly graphs; C, consistency scores of SPDs.

| Allele_1 | Allele_2 | Co | M | $M_r$ | $P_r$ | $P_{nongraph}$ | C |
|---|---|---|---|---|---|---|---|
| *dfrA7* | *sul1* | 19 | 100.00% | 100.00% | 100.00% | 100.00% | 1 |
| *strA*.173 | *sul2*.168 | 18 | 100.00% | 100.00% | 100.00% | 100.00% | 1 |
| *strB* | *sul2*.168 | 18 | 100.00% | 100.00% | 100.00% | 100.00% | 1 |
| *catA1*.215 | $bla_{OXA-1}$ | 11 | 100.00% | 100.00% | 100.00% | 100.00% | 1 |
| *dfrA1* | *sat-2A* | 10 | 100.00% | 100.00% | 100.00% | 100.00% | 1 |
| *dfrA7* | *sul1*.203 | 7 | 100.00% | 100.00% | 100.00% | 100.00% | 1 |
| *aadA1-pm*.182 | *catA1*.215 | 7 | 100.00% | 100.00% | 100.00% | 100.00% | 1 |
| *aadA1-pm*.182 | $bla_{OXA-1}$ | 7 | 100.00% | 100.00% | 100.00% | 100.00% | 1 |
| *mphA* | *sul1* | 5 | 100.00% | 100.00% | 100.00% | 100.00% | 1 |
| *aadA2*.195 | *sul1* | 3 | 100.00% | 100.00% | 100.00% | 100.00% | 1 |
| *aadA2*.195 | *catA1*.215 | 3 | 100.00% | 100.00% | 100.00% | 100.00% | 1 |
| *aadA2*.195 | *mphA* | 3 | 100.00% | 100.00% | 100.00% | 100.00% | 1 |
| *dfrA12* | *sul1* | 3 | 100.00% | 100.00% | 100.00% | 100.00% | 1 |
| *catA1*.215 | *dfrA12* | 3 | 100.00% | 100.00% | 100.00% | 100.00% | 1 |
| *dfrA12* | *mphA* | 3 | 100.00% | 100.00% | 100.00% | 100.00% | 1 |
| *qepA*.31 | *sul1* | 3 | 100.00% | 100.00% | 100.00% | 100.00% | 1 |
| *catA1*.215 | *qepA*.31 | 3 | 100.00% | 100.00% | 100.00% | 100.00% | 1 |
| *mphA* | *qepA*.31 | 3 | 100.00% | 100.00% | 100.00% | 100.00% | 1 |
| *aadA5* | *sul1* | 3 | 100.00% | 100.00% | 100.00% | 100.00% | 1 |
| *dfrA17* | *sul1* | 3 | 100.00% | 100.00% | 100.00% | 100.00% | 1 |
| *aadA1-pm*.181 | *dfrA1* | 2 | 100.00% | 100.00% | 100.00% | 100.00% | 1 |
| *aadA1-pm*.181 | *sat-2A* | 2 | 100.00% | 100.00% | 100.00% | 100.00% | 1 |
| *aac(3)-IId*.148 | *aadA2*.195 | 2 | 100.00% | 100.00% | 100.00% | 100.00% | 0 |
| *aac(3)-IId*.148 | *dfrA12* | 2 | 100.00% | 100.00% | 100.00% | 100.00% | 0 |
| *aac(3)-IId*.148 | *qepA*.31 | 2 | 100.00% | 100.00% | 100.00% | 100.00% | 0 |
| *aadA5* | *mphA* | 2 | 100.00% | 100.00% | 100.00% | 100.00% | 1 |
| *dfrA17* | *mphA* | 2 | 100.00% | 100.00% | 100.00% | 100.00% | 1 |
| *dfrA14*.227 | *sul2* | 30 | 96.67% | 96.67% | 100.00% | 82.76% | 1 |
| *strB* | *sul2* | 45 | 95.56% | 95.56% | 100.00% | 81.40% | 1 |
| *dfrA14*.227 | *strB* | 27 | 92.59% | 92.59% | 100.00% | 84.00% | 1 |
| *catA1*.215 | *mphA* | 4 | 75.00% | 75.00% | 100.00% | 100.00% | 1 |
| *strA*.173 | *strB* | 66 | 68.18% | 68.18% | 100.00% | 97.78% | 1 |
| *strB*.153 | *sul2* | 20 | 65.00% | 65.00% | 100.00% | 92.31% | 1 |
| *strA*.173 | *sul2* | 68 | 64.71% | 63.24% | 97.73% | 90.70% | 1 |
| *catA1*.215 | *sul1* | 8 | 100.00% | 62.50% | 62.50% | 100.00% | 0 |
| *strA*.173 | $bla_{TEM-214}$.147 | 70 | 78.57% | 37.14% | 47.27% | 100.00% | 1 |
| $bla_{TEM-214}$.147 | *tet*(A) | 43 | 100.00% | 34.88% | 34.88% | 93.33% | 0 |
| *dfrA8* | *strB*.153 | 18 | 61.11% | 33.34% | 54.55% | 100.00% | 1 |
| *dfrA8* | *sul2* | 22 | 100.00% | 22.73% | 22.73% | 100.00% | 1 |
| *strB* | *tet*(A) | 32 | 87.50% | 9.37% | 10.71% | 100.00% | 0 |
| *strA*.173 | *tet*(A) | 43 | 69.77% | 9.30% | 13.33% | 100.00% | 0 |
| *catA1*.215 | *dfrA1* | 11 | 100.00% | 9.09% | 9.09% | 100.00% | 0 |
| *dfrA7* | $bla_{TEM-214}$.147 | 27 | 100.00% | 7.41% | 7.41% | 100.00% | 0 |
| *dfrA7* | *strA*.173 | 27 | 100.00% | 3.70% | 3.70% | 100.00% | 0 |
| *dfrA7* | *strB* | 27 | 100.00% | 3.70% | 3.70% | 100.00% | 0 |

**Table s10: Physical distances measured between 15 pairs of positively associated alleles in 359 *Salmonella* genomes**. A minimum of two reliable SPDs were obtained for each pair. Abbreviations of column names: Co, co-occurrence count; M: measurability of all SPDs; $M_r$: measurability of reliable SPDs in all the SPDs; $P_r$, $M_r/M \times 100\%$ – percentage of reliable SPDs in all the SPDs; $P_{nongraph}$: percentage of reliable SPDs not measured in assembly graphs; C, consistency score of SPDs.

| Allele_1 | Allele_2 | Co | M | $M_r$ | $P_r$ | $P_{nongraph}$ | $S_d$ |
|---|---|---|---|---|---|---|---|
| *dfrA14*.79 | *strB* | 21 | 100.00% | 100.00% | 100.00% | 85.71% | 1 |
| *floR*.12 | *tet*(G) | 204 | 99.02% | 99.02% | 100.00% | 100.00% | 1 |
| *strB* | *sul2* | 20 | 95.00% | 95.00% | 100.00% | 73.68% | 1 |
| *dfrA14*.79 | *sul2* | 19 | 94.74% | 94.74% | 100.00% | 72.22% | 1 |
| *aadA2* | *sul1* | 318 | 92.77% | 92.45% | 99.66% | 14.63% | 0 |
| $bla_{CARB-2}$ | *sul1* | 288 | 92.36% | 92.01% | 99.62% | 4.53% | 1 |
| *aac(3)-Iva*.65 | *strB* | 5 | 80.00% | 60.00% | 75.00% | 66.67% | 1 |
| *aph(4)-Ia* | *strB* | 5 | 80.00% | 60.00% | 75.00% | 66.67% | 1 |
| *strA*.55 | *strB* | 16 | 25.00% | 25.00% | 100.00% | 75.00% | 1 |
| *strA*.55 | *sul2* | 14 | 14.29% | 14.29% | 100.00% | 50.00% | 1 |
| $bla_{CARB-2}$ | *floR*.12 | 202 | 92.08% | 4.95% | 5.38% | 30.00% | 1 |
| $bla_{CARB-2}$ | *tet*(G) | 279 | 92.47% | 4.66% | 5.04% | 23.08% | 1 |
| *aadA2* | *tet*(G) | 282 | 90.07% | 3.90% | 4.33% | 27.27% | 1 |
| *aadA2* | *floR*.12 | 204 | 88.73% | 2.94% | 3.31% | 33.33% | 1 |
| *sul1* | *tet*(G) | 282 | 88.65% | 2.84% | 3.20% | 50.00% | 1 |

**Table s11: SPDs measured in *E. coli* genomes between three alleles of the clique shown in Figure s16**. All SPDs were obtained from single nodes in assembly graphs.

| Allele1 | Allele2 | Distance (bp) | No. of distances |
|---|---|---|---|
| *aadA1-pm*.181 | *sat-2A* | 46 | 2 |
| *sat-2A* | *dfrA1* | 94 | 10 |
| *aadA1-pm*.181 | *dfrA1* | 665 | 2 |

**Table s12: Exact matches of the 3,084 bp MDR region in the genome assembly of the *Salmonella* genome ERR026101 to the NCBI nucleotide database**. All of these hits displayed the same bit score. The database was accessed in April, 2018.

| Species | Strain | Plasmid | Size (bp) | Accession | Coordinates |
|---|---|---|---|---|---|
| *Escherichia coli* | MS7163 | pMS7163B | 84,078 | CP026855 | 61,778–64,861 |
| *Escherichia coli* | 1283 | p7 | 6,800 | CP023375 | 2,487–5,570 |
| *Escherichia coli* | S1.2.T2R | pCERC1 | 6,790 | JN012467 | 97–3,180 |
| *Salmonella enterica* | SA20084699 | unnamed2 | 38,945 | CP022499 | 6,380–9,463 |
| *Shigella sonnei* | c8225 | pABC-3 | 6,779 | KT988306 | 97–3,180 |
| *Yersinia ruckeri* | 1521 | pYR1521 | 5,021 | HG423538 | 924–4,007 |

### 3 Supplementary details of implementation

In Section Implementation of the main article, we have outlined our network approach that identifies horizontally co-transferred alleles of AMR genes in bacterial genomes. Herein we provide a full mathematical justification of this approach. This section explains our association analysis and the APD assessment for network construction, and tests for significant structural random effects contributing to the allelic presence-absence of AMR genes. By convention, we use boldface upper-case letters to represent matrices, boldface lower-case letters for column vectors, and regular letters for scalars. All mathematical expressions are italicised. Abbreviations are the same as the main article.

#### 3.1 Association analysis controlling for population structure

In order to determine edges in the linkage network, we test for fixed effect of an explanatory allele on presence-absence of a response allele for each LMM that takes bacterial population structure and environmental randomness into account. Herein we derive a stringent procedure from existing methods for network construction.

##### 3.1.1 Representing allelic presence-absence status

The first step in our association analysis is to represent the presence-absence of alleles in all bacterial genomes using a matrix. Assuming that $m$ alleles of $M$ genes ($m \geqslant M$) are identified in $n$ genomes, let an $n \times m$ binary matrix $\boldsymbol{A} = (a_{ij})$ represent the presence-absence of every allele across genomes, where the $(i, j)$-th element $a_{ij}$ of $\boldsymbol{A}$ equals one if the $j$-th allele is present in the $i$-th genome, and equals zero otherwise (see the manual of GEMMA [4]). This designation of one and zero to presence-absence status makes the explanation of results more straightforward, although it is merely arbitrary and does not change any conclusions. Following this designation, the matrix is $\boldsymbol{A}$ becomes an allelic PAM, where rows represent genomes and columns represent alleles. In particular, we do not include any allele that does not show variation in its distribution, namely, any allele showing a frequency of zero or one is excluded from our analysis in order to observe a fundamental assumption for linear models – variables must be random. Problems arise when this assumption is violated. For example, a perfect fit of an explanatory variable to a constant response is seen in a linear model, where the coefficient of the explanatory variable equals zero as expected.

##### 3.1.2 Identifying presence-absence patterns

In practice, it is not unusual to see several alleles sharing the same distribution in samples. For instance, the allelic co-transfer of the tetracycline resistance gene *tet*(G) and its regulatory gene *tetR*(G) between *S.* Typhimurium has been reported [5]. Mathematically, identically distributed alleles are interchangeable in association tests and produce the same result. As a result, these duplicated tests lead to an excessively rigorous adjustment of p-values for controlling false positives as they enlarge the number of tests. Consequently, the power of tests is compromised. To retain the power, we can learn from the R package BugWAS [6] and take a single allele from each group of identically distributed alleles as a representative for all relevant association tests and call this representative as a presence-absence pattern.

Assuming there are $p$ patterns representing $m$ alleles, where $p \leqslant m$, we can compress the $n \times m$ allelic PAM $\boldsymbol{A}$ into an $n \times p$ binary matrix $\boldsymbol{B} = (b_{ij})$, whose rows denote samples and columns denote patterns. We call $\boldsymbol{B}$ a pattern matrix. In the following example, we merge the first and fourth columns, the third and fifth columns of $\boldsymbol{A}$, respectively, into two columns to make a pattern matrix $\boldsymbol{B}$. Note that neither rows nor columns of $\boldsymbol{A}$ and $\boldsymbol{B}$ have to be sorted.

$$\boldsymbol{A} = \begin{bmatrix} 1 & 1 & 1 & 1 & 1 \\ 0 & 1 & 1 & 0 & 1 \\ 0 & 0 & 1 & 0 & 1 \end{bmatrix} \Rightarrow \boldsymbol{B} = \begin{bmatrix} 1 & 1 & 1 \\ 0 & 1 & 1 \\ 0 & 0 & 1 \end{bmatrix} \tag{1}$$

##### 3.1.3 Column-wise zero-centring of the pattern matrix

Zero-centring random variables of the same population by their arithmetic means is a common technique for simplifying algebra without changing the distribution of data points or affecting results. Herein, we treat each pattern as a column vector of $n$ dichotomous variables representing presence-absence of the same allele in $n$ genomes. Accordingly, we define an $n \times p$ column-wisely zero-centred pattern matrix $\boldsymbol{X} = (x_{ij})$ as follows:

$$x_{ij} = b_{ij} - \frac{1}{n}\sum_{k=1}^{n} b_{kj} = b_{ij} - \bar{b}_{.j}, \text{ where } 1 \leqslant i \leqslant n \text{ and } 1 \leqslant j \leqslant p \tag{2}$$

Accordingly, the presence-absence status of an allele belonging to the $j$-th pattern in the $i$-th genome appears as following in the centred pattern matrix $\boldsymbol{X}$:

$$x_{ij} = \begin{cases} 1 - \bar{b}_{.j} > 0, & \text{presence} \\ -\bar{b}_{.j} < 0, & \text{absence} \end{cases} \tag{3}$$

Note that the column mean $\bar{b}_{.j}$ actually equals the frequency of each allele represented by the $j$-th pattern in $n$ genomes. It is known that every column of $\boldsymbol{X}$ sums to zero:

$$\sum_{k=1}^{n} x_{kj} = (1 - \bar{b}_{.j})(\bar{b}_{.j}n) + (-\bar{b}_{.j})(n - \bar{b}_{.j}n) = 0 \tag{4}$$

This property applies to other zero-centred binary matrices.

##### 3.1.4 Genotype matrix of biallelic core-genome SNPs

The construction of a genotype matrix for biallelic core-genome SNPs (cgSNPs) brings in genetic variations for estimating population structure of bacterial genomes. In our approach, a cgSNP is strictly defined as a single-nucleotide polymorphic site that is present in all genomes. This constraint is a limitation of current methods that incorporate population structure into linear models using principal components (PCs) [4, 7].

Assuming there are $L$ biallelic cgSNPs identified in $n$ genomes and $n < L$, we define an $n \times L$ binary genotype matrix $\boldsymbol{G} = (g_{ij})$, where $1 \leqslant i \leqslant n$, $1 \leqslant j \leqslant L$, and

$$g_{ij} = \begin{cases} 0, & \text{major allele} \\ 1, & \text{minor allele} \end{cases} \tag{5}$$

We treat each SNP as a dichotomous random variable observed in $n$ genomes. Accordingly, we can also zero-centre columns of $\boldsymbol{G}$ by column means to simplify algebra, creating an $n \times L$ column-wise zero-centred genotype matrix $\boldsymbol{S} = (s_{ij})$:

$$s_{ij} = g_{ij} - \frac{1}{n}\sum_{k=1}^{n} g_{kj} = g_{ij} - \bar{g}_{.j} = \begin{cases} -\bar{g}_{.j} < 0, & \text{major allele} \\ 1 - \bar{g}_{.j} > 0, & \text{minor allele} \end{cases} \tag{6}$$

Note that the column mean $\bar{g}_{.j}$ equals the minor allele frequency (MAF) of the $j$-th cgSNP in $n$ genomes. According to Equation 4, we know that every column of $\boldsymbol{S}$ sums up to zero as well. Furthermore, the maximum rank of $\boldsymbol{S}$ reduces by 1 from $n$ as its columns have been zero-centred [8]. Hence we have:

$$\text{rank}(\boldsymbol{S}) \leqslant n - 1 \tag{7}$$

More generally, we have $\text{rank}(\boldsymbol{S}) \leqslant \min\{n-1, L\}$ when removing the assumption that $n < L$ for the SNP matrix.

##### 3.1.5 Calculation of a relatedness matrix

A relatedness matrix captures population structure and plays a pivotal role in introducing the population structure into linear models. In our implementation of GeneMates, the function *findPhysLink* calls GEMMA to calculate this relatedness matrix [4]. Following the manual of GEMMA (github.com/genetics-statistics/GEMMA), we calculate an $n \times$

$n$ relatedness matrix $\boldsymbol{K} = (k_{ij})$ from the centred SNP matrix $\boldsymbol{S}$ (Note that GEMMA performs column-wise zero-centring on $\boldsymbol{G}$ before calculating $\boldsymbol{K}$) with formula

$$\boldsymbol{K} = \frac{\boldsymbol{S}\boldsymbol{S}^T}{L} \tag{8}$$

where the superscript T denotes a matrix transpose and this notation will be used throughout this article. The relatedness matrix $\boldsymbol{K}$ reveals all-to-all relationships between the $n$ genomes. It is a symmetric matrix because

$$k_{ij} = \frac{1}{L}\sum_{r=1}^{L} s_{ir}s_{jr} = \frac{1}{L}\sum_{r=1}^{L} s_{jr}s_{ir} = k_{ji} \tag{9}$$

where $1 \leqslant i, j \leqslant n$. As such, both rows and columns of $\boldsymbol{K}$ denote the samples. Moreover, given the inequality (Formula 7), the relatedness matrix $\boldsymbol{K}$ is positive semidefinite and $\text{rank}(\boldsymbol{K}) = \text{rank}(\boldsymbol{S})$ (Theorems 2.6D and 2.4A in the book by A.C. Rencher [9]).

##### 3.1.6 Singular-value decomposition of the SNP matrix

This is a critical step for converting the population structure into an orthogonal form, which can be incorporated into an LMM afterwards for term of structural random effects. Let $r = \text{rank}(\boldsymbol{K})$. Since $\boldsymbol{K}$ is a symmetric matrix of order $n$, we can perform eigen-decomposition on it, which returns $n$ real eigenvalues (cf. Theorem 2.12C in [9]) and $n$ accompanying linearly independent column vectors, even though some eigenvalues may be the same. Moreover, the eigenvalues must not be negative but may equal zero, because $\boldsymbol{K}$ is a positive semidefinite matrix. Let $\lambda_1 \geqslant \lambda_2 \geqslant \cdots \geqslant \lambda_n \geqslant 0$ represent these eigenvalues sorted in a descending order. Note that there must be positive eigenvalues because the square matrix $\boldsymbol{K}$ is positive semidefinite (To put it simple, there must be positive eigenvalues of $\boldsymbol{K}$ because the sum of its eigenvalues equals its trace and the trace must be positive as the entries on the main diagonal of the relatedness matrix, i.e., entries representing self-relatedness, must be positive). To avoid confusions, we refer an eigenvector (in a narrow sense) of $\boldsymbol{K}$ to an orthonormal vector obtained from the linearly independent vectors aforementioned through the Gram–Schmidt process and subsequent normalisation, although in a broad sense, all of these untransformed vectors are also eigenvectors of $\boldsymbol{K}$ (linearly independent, but are not necessarily orthogonal). Note that the Gram–Schmidt process itself shows that it retains the link between eigenvalues and broad-sense eigenvectors when it is applied.

Therefore, we obtain and can only obtain $n$ eigenvectors $\boldsymbol{e}_1, \cdots, \boldsymbol{e}_n$ corresponding to the non-negative eigenvalues $\lambda_1, \cdots, \lambda_n$ of $\boldsymbol{K}$. By definition, these eigenvectors are orthonormal bases of an $n$ dimensional real Euclidean space $V^n \subset \mathbb{R}^n$, in which each genome is a data point pinned down by $n$ coordinates. Note that the orientation of each base (hence that of the axis) is merely arbitrary and relies on the corresponding eigenvalue. As a result, reversing one eigenvector has no impact on the orthonormality of bases. Using the $n$ eigenvectors, we can construct an $n \times n$ matrix $\boldsymbol{E} = [\boldsymbol{e}_1 \cdots \boldsymbol{e}_n]$. This is an orthonormal matrix as $\boldsymbol{E}^T\boldsymbol{E} = \boldsymbol{E}\boldsymbol{E}^T = \boldsymbol{I}_n$ (an identity matrix of order $n$) and we can immediately know that $\boldsymbol{E}^{-1} = \boldsymbol{E}^T$. Since $\boldsymbol{K}$ is a symmetric matrix of real numbers and $\boldsymbol{E}$ is invertible, we have $\boldsymbol{E}^{-1}\boldsymbol{K}\boldsymbol{E} = \mathrm{diag}(\lambda_1, \lambda_2, \cdots, \lambda_n)$ and $r = \mathrm{rank}(\boldsymbol{K}) = \mathrm{rank}(\boldsymbol{E}^{-1}\boldsymbol{K}\boldsymbol{E})$. Therefore, $r$ is the number of non-zero (hence positive) eigenvalues of $\boldsymbol{K}$, and $n - r$ equals the number of its zero eigenvalues (cf. Chapter 2.12.5 in [9]). As we will be demonstrating in the following algebra, this is an important property for obtaining correct transformation of population structure, however, it has not been taken into account in literature so far to our knowledge.

Further, since a singular value of $\boldsymbol{K}$ is defined as the non-negative square root of one of its eigenvalues, there is always an equal number of singular values and eigenvalues of the same relatedness matrix, regardless whether there are duplicated values or not. Using SVD on real matrices, we can decompose the biallelic cgSNP matrix $\boldsymbol{S}$ into a product of matrices:

$$\boldsymbol{S}_{n \times L} = \boldsymbol{P}_{n \times n} \boldsymbol{\Sigma}_{n \times L} \boldsymbol{Q}^T_{L \times L} \tag{10}$$

where the matrices

$$\boldsymbol{P} = \begin{bmatrix} \boldsymbol{U}_{n \times r} & \boldsymbol{N}_{n \times (n-r)} \end{bmatrix} \tag{11}$$

$$\boldsymbol{\Sigma} = \begin{bmatrix} \boldsymbol{D}_{r \times r} & \boldsymbol{O}_{r \times (L-r)} \\ \boldsymbol{O}_{(n-r) \times r} & \boldsymbol{O}_{(n-r) \times (L-r)} \end{bmatrix} \tag{12}$$

$$\boldsymbol{Q}^T = \begin{bmatrix} \boldsymbol{V}_{L \times r} & \boldsymbol{W}_{L \times (L-r)} \end{bmatrix}^T \tag{13}$$

To be more specific, columns of $\boldsymbol{P}$ are eigenvectors (also known as the left singular vectors) of $\boldsymbol{S}\boldsymbol{S}^T = L\boldsymbol{K}$, which correspond to the $r$ positive eigenvalues and $n - r$ zero eigenvalues (notice eigenvectors of $L\boldsymbol{K}$ are the same as $\boldsymbol{K}$ but eigenvalues are $L$ times those of $\boldsymbol{K}$); columns of $\boldsymbol{Q}$ are eigenvectors (right singular vectors) of $\boldsymbol{M} = (m_{ij}) = \boldsymbol{S}^T\boldsymbol{S}$ (called a scatter matrix), which correspond to the same $r$ positive eigenvalues and $L - r$ zero eigenvalues of $L\boldsymbol{K}$; $\boldsymbol{D}$ is a diagonal square matrix of $r$ positive singular values of both $\boldsymbol{S}\boldsymbol{S}^T$ and $\boldsymbol{S}^T\boldsymbol{S}$; and $\boldsymbol{O}$ denotes a zero matrix of a given size. Both matrices $\boldsymbol{P}$ and $\boldsymbol{Q}$ are orthonormal. In addition, the scatter matrix $\boldsymbol{M}$ equals $n - 1$ times the genome variance-covariance matrix of un-centred cgSNP genotypes because

$$
\begin{aligned}
m_{ij} &= \sum_{k=1}^{n} s_{ki}s_{kj} = (n-1)\sum_{k=1}^{n}\frac{s_{ki}s_{kj}}{n-1} = (n-1)\sum_{k=1}^{n}\frac{(g_{ki}-\bar{g}_{.i})(g_{kj}-\bar{g}_{.j})}{n-1} \\
&= (n-1)\,\mathrm{Cov}(\boldsymbol{g}_i,\boldsymbol{g}_j)
\end{aligned} \tag{14}
$$

where $g_i$, $g_j$ denote the $i$-th and $j$-th column of the un-centred cgSNP matrix $\boldsymbol{G}$, respectively.

For conciseness, singular values are arranged in a descending order. Therefore, each of the matrices $\boldsymbol{U}$ and $\boldsymbol{V}$ is comprised of $r$ eigenvectors corresponding to the $r$ positive eigenvalues, and the matrices $\boldsymbol{N}$ and $\boldsymbol{W}$ are comprised of $n-r$ and $L-r$ eigenvectors corresponding to zero eigenvalues, respectively. Now we show that

$$
\begin{aligned}
\boldsymbol{S}_{n\times L} &= \begin{bmatrix}\boldsymbol{U}_{n\times r} & \boldsymbol{N}_{n\times(n-r)}\end{bmatrix}\begin{bmatrix}\boldsymbol{D}_{r\times r} & \boldsymbol{O}_{r\times(L-r)} \\ \boldsymbol{O}_{(n-r)\times r} & \boldsymbol{O}_{(n-r)\times(L-r)}\end{bmatrix}\begin{bmatrix}\boldsymbol{V}^T_{L\times r} \\ \boldsymbol{W}^T_{L\times(L-r)}\end{bmatrix} \\
&= \begin{bmatrix}\boldsymbol{UD} & \boldsymbol{O}_{n\times(L-r)}\end{bmatrix}\begin{bmatrix}\boldsymbol{V}^T_{L\times r} \\ \boldsymbol{W}^T_{L\times(L-r)}\end{bmatrix} = \boldsymbol{UDV}^T
\end{aligned} \tag{15}
$$

Accordingly, we can deduce that $\mathrm{rank}(\boldsymbol{S}) = \mathrm{rank}(\boldsymbol{P\Sigma Q}^T) = \mathrm{rank}(\boldsymbol{\Sigma})$ because $\mathrm{rank}(\boldsymbol{\Sigma}) = \mathrm{rank}(\boldsymbol{D}) = r$ and columns of $\boldsymbol{P}$ and $\boldsymbol{Q}$ are orthonormal (hence both matrices are non-singular and invertible). Notice neither $\boldsymbol{U}$ nor $\boldsymbol{V}$ is invertible when $r < n$ because they are not square matrices under this condition, and then we can only have $\boldsymbol{U}^T\boldsymbol{U} = \boldsymbol{V}^T\boldsymbol{V} = \boldsymbol{I}_r$.

Since matrices $\boldsymbol{N}$ and $\boldsymbol{W}$ always get cancelled out in Equation 15, we call Equation $\boldsymbol{S} = \boldsymbol{UDV}^T$ the reduced form of SVD, which is equivalent to the full form, $\boldsymbol{S} = \boldsymbol{P\Sigma Q}^T$. Consequently, we can completely recover $\boldsymbol{S}$ only with $r$ eigenvectors in $\boldsymbol{U}$ and $\boldsymbol{V}$ corresponding to the $r$ positive singular values in $\boldsymbol{D}$ instead of using all eigenvectors in $\boldsymbol{P}$ and $\boldsymbol{Q}$. This substitution reduces computational expense. Nonetheless, as we will demonstrate later, we can benefit from the orthonormal matrices in the full form of SVD in simplifying some equations.

##### 3.1.7 Projecting data points on axes defined by eigenvectors

**Projections can be acquired through both the full and reduced forms of SVD** We consider every bacterial genome as a data point in an $L$ dimensional real Euclidean space $\boldsymbol{V}^L \subset \mathbb{R}^L$ using genotypes of $L$ biallelic cgSNPs as coordinates. These coordinates may not be linearly independent because of homoplasy, parallel evolution, linkage disequilibrium, SNP-call errors, and so forth. Noticing $\boldsymbol{Q}^T\boldsymbol{Q} = \boldsymbol{I}_L$, we obtain an orthogonal transformation of rows (that is, coordinate vectors of genomes) in $\boldsymbol{S}$ with the orthonormal matrix $\boldsymbol{Q}$ by the equation

$$\boldsymbol{S} = \boldsymbol{P\Sigma Q}^T \Leftrightarrow \boldsymbol{SQ} = \boldsymbol{P\Sigma} \tag{16}$$

Let an $n \times L$ matrix $\boldsymbol{C}_L = \boldsymbol{SQ} = \boldsymbol{P\Sigma}$, where $\boldsymbol{C}_L = [\boldsymbol{c}_1 \cdots \boldsymbol{c}_L]$ and the length-$n$ column vector $\boldsymbol{c}_i$ $(1 \leqslant i \leqslant L)$ is the $i$-th column of $\boldsymbol{C}_L$. Similarly, we define the $j$-th $(1 \leqslant j \leqslant n)$ column of $\boldsymbol{P}$ and the $j$-th singular value in $\boldsymbol{\Sigma}$ as $\boldsymbol{p}_j$ and $\sigma_j$, respectively. Then we have

$$\begin{aligned} \boldsymbol{C}_L = \boldsymbol{P\Sigma} &= \begin{bmatrix} \boldsymbol{p}_1 & \cdots & \boldsymbol{p}_r & \boldsymbol{p}_{r+1} & \cdots & \boldsymbol{p}_n \end{bmatrix} \begin{bmatrix} \mathrm{diag}(\sigma_1, \cdots, \sigma_r, 0, \cdots, 0) & \boldsymbol{O}_{n\times(L-n)} \end{bmatrix} \\ &= \begin{bmatrix} \sigma_1 \boldsymbol{p}_1 & \cdots & \sigma_r \boldsymbol{p}_r & 0\boldsymbol{p}_{r+1} & \cdots 0\boldsymbol{p}_n & \boldsymbol{0} & \cdots & \boldsymbol{0} \end{bmatrix} \\ &= \begin{bmatrix} \sigma_1 \boldsymbol{p}_1 & \cdots & \sigma_r \boldsymbol{p}_r & \boldsymbol{0} & \cdots & \boldsymbol{0} \end{bmatrix}_{n\times L} \end{aligned} \tag{17}$$

Hence $\boldsymbol{C}_L = [\sigma_1 \boldsymbol{p}_1 \quad \cdots \quad \sigma_r \boldsymbol{p}_r \quad \boldsymbol{0} \quad \cdots \quad \boldsymbol{0}]$, which consists of $L - r$ zero column vectors. Since $\boldsymbol{c}_i = \sigma_i \boldsymbol{p}_i$, $1 \leqslant i \leqslant n$, we know that $\boldsymbol{c}_i$ equals the $i$-th eigenvector of the scatter matrix $\boldsymbol{S}^T\boldsymbol{S}$ (positive semidefinite, of the same rank as $\boldsymbol{S}$) scaled by its $i$-th singular value $\sigma_i$. We notice the non-zero partition of $\boldsymbol{C}_L$ in (17) can be acquired via the reduced form of SVD:

$$\boldsymbol{S} = \boldsymbol{UDV}^T \Leftrightarrow \boldsymbol{SV} = \boldsymbol{UD} = \begin{bmatrix} \boldsymbol{p}_1 & \cdots & \boldsymbol{p}_r \end{bmatrix} \begin{bmatrix} \sigma_1 & \cdots & 0 \\ \vdots & \ddots & \vdots \\ 0 & \cdots & \sigma_r \end{bmatrix} = \begin{bmatrix} \sigma_1 \boldsymbol{p}_1 & \cdots & \sigma_r \boldsymbol{p}_r \end{bmatrix} \tag{18}$$

which gives $\boldsymbol{C}_r = [\boldsymbol{c}_1 \quad \cdots \quad \boldsymbol{c}_r] = [\sigma_1 \boldsymbol{p}_1 \quad \cdots \quad \sigma_r \boldsymbol{p}_r]$, where the notation $\boldsymbol{C}_r$ represents a sub-matrix comprised of the first $r$ columns of $\boldsymbol{C}_L$. As elucidated in the following paragraphs, vectors $\boldsymbol{c}_1, \cdots, \boldsymbol{c}_n$ are projections of genomes onto axes defined by eigenvectors $\boldsymbol{p}_1, \cdots, \boldsymbol{p}_n$.

**The projections remain in the same Euclidean space** Since the inner product $\boldsymbol{c}_i^T \boldsymbol{c}_j = \sigma_i \boldsymbol{p}_i^T \sigma_j \boldsymbol{p}_j = 0$, Equation (17) also illustrates that vectors $\boldsymbol{c}_1$, ..., $\boldsymbol{c}_n$ are orthogonal and in parallel with the orthonormal bases $\boldsymbol{p}_1, \ldots, \boldsymbol{p}_n$ of the $n$ dimensional Euclidean space $\boldsymbol{V}^n$ described previously.

Similarly, we expand the $n \times L$ matrix product $\boldsymbol{SQ}$ in Equation 16 into a matrix comprised of inner products of vectors:

$$\boldsymbol{SQ} = \begin{bmatrix} \boldsymbol{s}_1 \\ \vdots \\ \boldsymbol{s}_n \end{bmatrix} \begin{bmatrix} \boldsymbol{q}_1 & \cdots & \boldsymbol{q}_L \end{bmatrix} = \begin{bmatrix} \boldsymbol{s}_1\boldsymbol{q}_1 & \cdots & \boldsymbol{s}_1\boldsymbol{q}_L \\ \vdots & \ddots & \vdots \\ \boldsymbol{s}_n\boldsymbol{q}_1 & \cdots & \boldsymbol{s}_n\boldsymbol{q}_L \end{bmatrix} = \begin{bmatrix} \boldsymbol{c}_1 & \cdots & \boldsymbol{c}_L \end{bmatrix} \tag{19}$$

where $\boldsymbol{s}_i$ ($i = 1, \cdots, n$) is the $i$-th row vector $[s_{i1}, \ldots, s_{iL}]$ of $\boldsymbol{S}$ and $\boldsymbol{c}_j$ ($j = 1, \cdots, L$) is a column vector of length $n$. The vector $\boldsymbol{c}_j$ shows "coordinates" (strictly specaking, scaling coefficients of bases that defining axes) on $L$ "axes" (namely, SNP genotype codes taking values of either one or zero and usually are mutually correlated between genomes and hence are not genuine bases of a linear space) that locate the $i$-th genome in the space $\boldsymbol{V}^L$. Therefore, $\boldsymbol{c}_j = [\boldsymbol{s}_1\boldsymbol{q}_j \ \cdots \ \boldsymbol{s}_n\boldsymbol{q}_j]^T$ where $j = 1, \cdots, L$. Note that every one of $\boldsymbol{c}_{r+1}, \cdots, \boldsymbol{c}_L$ equals $\boldsymbol{0}$ following Equation 17. We can also write the $i$-th row of $\boldsymbol{C}_L$ in the form:

$$\begin{bmatrix} c_{i1} & \cdots & c_{iL} \end{bmatrix} = \begin{bmatrix} \boldsymbol{s}_i\boldsymbol{q}_1 & \cdots & \boldsymbol{s}_i\boldsymbol{q}_L \end{bmatrix} = \boldsymbol{s}_i\boldsymbol{Q} \tag{20}$$

which represents an orthonormal transformation of the $\boldsymbol{V}^L$ itself (denoted by $\boldsymbol{V}^L \to \boldsymbol{V}^L$) and illustrates that $c_{i1}, \cdots, c_{iL}$ are projections of coordinates $s_{i1}, \cdots, s_{iL}$ via the orthonormal matrix $\boldsymbol{Q}$. Since $\boldsymbol{c}_i$ is orthogonal to $\boldsymbol{c}_j$ when $i \neq j$, we transform correlated vectors $[s_{1i}, \ldots, s_{ni}]^T$ and $[s_{1j}, \ldots, s_{nj}]^T$ ("coordinates" of the $n$ genomes on the $i$-th and $j$-th axes of SNPs) into orthogonal coordinates $\boldsymbol{c}_i$ and $\boldsymbol{c}_j$ with Equation 20 as a benefit of SVD. This equation is known as a rotation transformation, which preserves the distribution of data points but establishes a set of orthogonal axes going through the same origin (i.e., builds another coordinate system). Therefore, projections on every new axis, namely, elements in the vector $\boldsymbol{c}_i$, are zero-centred as well. In addition, the matrix $\boldsymbol{Q}$ is also known as the rotation matrix.

**The projections and matrices show profound interconnections** As for the $i$-th element in $\boldsymbol{c}_j$, it follows

$$c_{ij} = \boldsymbol{s}_i\boldsymbol{q}_j = \sum_{k=1}^{L} q_{kj}s_{ik} \tag{21}$$

which means the coordinate of the $i$-th genome on the $j$-th axis (either defined by $\boldsymbol{p}_j$ when $j \leqslant n$ or being $\boldsymbol{0}$ with any directions when $n < j \leqslant L$) is a linear combination of all of its SNP "coordinates" using elements in $\boldsymbol{q}_j$ as weights. This equation reveals profound connections between the SNP matrix $\boldsymbol{S}$, the relatedness matrix $\boldsymbol{K}$ of genomes, the variance-covariance matrix $\boldsymbol{M}$ of biallelic cgSNPs, and the projections of genomes on a group of orthogonal axes through SVD.

**Principal components are bases establishing $r$ orthogonal axes where data points are projected onto** According to the reduced form of SVD (Equation 18) and Equation 20, the $r$ eigenvectors $\boldsymbol{p}_1, \cdots, \boldsymbol{p}_r$, which are columns of $\boldsymbol{U}$ and bases of an $r$ dimensional Euclidean space $\boldsymbol{V}^r$, are called PCs of $\boldsymbol{S}$ [7, 8]. As shown in Equation 18, projections $\boldsymbol{c}_1, \cdots, c_r$ of the $n$ data points on the $i$-th axis are coordinates or the length of the scaled $i$-th PC, which are hence referred to as scores in literature. Since the PCs are ranked by their accompanying eigenvalues in a descending order, people often use the first a few PCs to capture the majority of variance in data for an approximation in their analysis (for example, the principal component regression [10]), which is usually more computationally efficient than use all PCs. For GeneMates, however, we use all PCs in order to capture all variance in the SNP data (Figure s8).

**Each eigenvalue measures the percentage of genetic variation captured by the corresponding principal component** Considering the variance of data projections on the $j$-th ($j \leqslant n$) axis, namely, the variance of the length of the $j$-th PC, since the projections remain zero-centred, their arithmetic mean equals zero. Accordingly, we can determine the sample variance of $c_{1j}, \cdots, c_{nj}$ through the equation

$$\mathrm{Var}(\boldsymbol{c}_j) = \mathrm{Var}(c_{1j}, \cdots, c_{nj}) = \frac{1}{n-1} \sum_{i=1}^{n} \left(c_{ij} - \bar{c}_{.j}\right)^2 = \frac{1}{n-1} \sum_{i=1}^{n} c_{ij}^2 = \frac{\boldsymbol{c}_j^T \boldsymbol{c}_j}{n-1} \tag{22}$$

According to Equation 17, since $\boldsymbol{c}_j^T \boldsymbol{c}_j = (\sigma_j \boldsymbol{p}_j)^T (\sigma_j \boldsymbol{p}_j) = \sigma_j^2 \boldsymbol{p}_j^T \boldsymbol{p}_j = \lambda_j \cdot 1$, we have

$$\mathrm{Var}(\boldsymbol{c}_j) = \frac{\lambda_j}{n-1} \propto \lambda_j, (j \leqslant n) \tag{23}$$

which shows that the projections of data points $1, \cdots, n$ on the $j$-th axis has a variance proportional to the $j$-th eigenvalue. This equation also shows that the singular value $\sigma_j$ can be obtained from $\sigma(\boldsymbol{c}_j)$, the standard deviation of observed lengths of the $j$-th PC:

$$\sigma_j = \sqrt{\lambda_j} = \sqrt{n-1}\,\sigma(\boldsymbol{c}_j) \tag{24}$$

**The rank of the relatedness matrix $\boldsymbol{K}$ determines the minimum number of principal components required for capturing all genetic variation in sample genomes** Equation 23 shows that projections on the first $r$ axes (in parallel with orthonormal bases $\boldsymbol{p}_1, \cdots, \boldsymbol{p}_r$) always have variances greater than zero (hence are informative), while projections on the other $n-r$ axes (in parallel with orthonormal bases $\boldsymbol{p}_{r+1}, \cdots, \boldsymbol{p}_n$) all fall into the origin and hence do not show any variance (uninformative). Moreover, we do not consider projections on the rest of $L-n$ axes because these axes are always $\mathbf{0}$ of any directions and all the projections do not diverge from the origin either. Consequently, all variances in the distribution of data points are captured by the first $r$ axes, and the proportion of total variance captured by the $i$-th ($i \leqslant r$) axis equals $\lambda_i$ divided by the sum of all $r$ positive eigenvalues.

##### 3.1.8 Univariate linear mixed models and parameter estimation

For genes of interest, we use LMMs to explain the presence-absence of a response allele with a fixed effect of the presence-absence of an explanatory allele, additive random effects of population structure, and environmental random effects. Specifically, alleles are represented with patterns as described in Section 3.1.2. For any two of $p$ columns $\boldsymbol{x}$ and $\boldsymbol{y}$ in the zero-centred pattern matrix $\boldsymbol{X}$, where $\boldsymbol{x} \neq \boldsymbol{y}$, we consider $\boldsymbol{y}$ as a sum of a fixed effect of $\boldsymbol{x}$, additive random effects of population structure and random environmental errors. In addition, the effects of population structure can be called structural random effects regarding to the bacterial population structure. In literature, they are also known as background effects and lineage effects [6]. Following our notations, we construct a univariate LMM with four parameters to explain observations in the vector $\boldsymbol{y}$:

$$\boldsymbol{y} = \mathbf{1}\alpha + \boldsymbol{x}\beta + \boldsymbol{C}_L\boldsymbol{\gamma} + \boldsymbol{\varepsilon} \tag{25}$$

$$\boldsymbol{\gamma} \sim MVN_L\left(\mathbf{0}, \lambda\tau^{-1}L^{-1}\boldsymbol{I}_L\right) \tag{26}$$

$$\boldsymbol{\varepsilon} \sim MVN_n\left(\mathbf{0}, \tau^{-1}\boldsymbol{I}_n\right) \tag{27}$$

where $\alpha$ is the coefficient for the intercept term and $\beta$ is the fixed effect size of the explanatory pattern $\boldsymbol{x}$; $\boldsymbol{\gamma}$ is a column vector of length $L$, which represents sizes of structural random effects of $\boldsymbol{c}_1, \cdots, \boldsymbol{c}_L$ on the response vector $\boldsymbol{y}$; finally, the error term $\boldsymbol{\varepsilon}$ of length $n$ represents residuals between data points and the mean of $\boldsymbol{y}$ under the model. Four parameters $\alpha$, $\beta$, $\lambda$ and $\tau$ of the model will be estimated based on observations. Note that there is only a constant term $\mathbf{1}\alpha$ in the model for covariates as we do not take other variables into account at present.

Particularly, authors of GEMMA defined two components that constitute the total variance in random effects in an LMM. Specifically, $\sigma_e^2 = \tau^{-1}$ is the environmental variance component and $\sigma_g^2 = \lambda\sigma_e^2 = \lambda\tau^{-1}$ is the structural variance component (In the GEMMA paper, they are called environmental effect and genetic effect, respectively [4]). Therefore, $\lambda = \sigma_g^2/\sigma_e^2$, which measures the dominance of population structure over environmental randomness in random effects. Accordingly, we can rewrite the assumed distributions of random effects in the LMM (Equation 25) as

$$\boldsymbol{\gamma} \sim MVN_L\left(\mathbf{0}, L^{-1}\sigma_g^2\boldsymbol{I}_L\right) \tag{28}$$

$$\boldsymbol{\varepsilon} \sim MVN_n\left(\mathbf{0}, \sigma_e^2\boldsymbol{I}_n\right) \tag{29}$$

As already shown in Equation 17, there are only $r \leqslant n-1$ orthogonal axes having projections diverging from the origin. Accordingly, the total effect of population structure reduces to the form

$$\boldsymbol{C}_L\boldsymbol{\gamma} = \sum_{j=1}^{r} \gamma_j \boldsymbol{c}_j + \sum_{j=L-r}^{L} \gamma_j \boldsymbol{0} = \boldsymbol{C}_r\boldsymbol{\gamma}_r \tag{30}$$

where the column vector $\boldsymbol{\gamma}_r$ is comprised of the leading $r$ elements of $\gamma$ without changing their order. Therefore, we can simplify the model defined in Equation 25 into an equivalent form:

$$\boldsymbol{y} = \boldsymbol{1}\alpha + \boldsymbol{x}\beta + \boldsymbol{C}_r\boldsymbol{\gamma}_r + \boldsymbol{\varepsilon} \tag{31}$$

$$\boldsymbol{\gamma}_r \sim MVN_r\left(\boldsymbol{0}, \lambda\tau^{-1}L^{-1}\boldsymbol{I}_r\right) \tag{32}$$

and the error term $\boldsymbol{\varepsilon}$ follows the same distribution as in Equation 27.

##### 3.1.9 Parameter estimation

We use GEMMA to estimate the four parameters in Model 25. In this section, we only outline key algebra for the estimates to demonstrate their forms specifically in our model. Readers may read the original paper and manual of GEMMA for more details [4]. Herein, for allelic presence-absence status in $n$ genomes, we use the notation $\boldsymbol{x}$ to denote a column vector for an explanatory pattern, and use $\boldsymbol{y}$ to denote the other column vector for a response pattern. Both vectors have already been zero-centred by their genome means. Note that the designation of response and explanatory vectors is arbitrary and in practice, patterns are iterated for both roles in the LMM to make all-to-all contrasts. Assuming $\boldsymbol{x} \neq \boldsymbol{y}$ and $n \gg 2$, we specify GEMMA to estimate the parameters using a residual maximum-likelihood (REML) approach and obtain unbiased parameter estimates for random effects. The target function for our model to optimise is

$$\begin{aligned} l_{r1}(\lambda, \tau; \boldsymbol{y}, \boldsymbol{x}, \boldsymbol{K}) = {} & \frac{n-2}{2}\log\tau - \frac{n-2}{2}\log(2\pi) + \frac{1}{2}\log\det\left(\begin{bmatrix}\boldsymbol{1} & \boldsymbol{x}\end{bmatrix}^T \begin{bmatrix}\boldsymbol{1} & \boldsymbol{x}\end{bmatrix}\right) \\ & -\frac{1}{2}\log\det\boldsymbol{H} - \frac{1}{2}\log\det\left(\begin{bmatrix}\boldsymbol{1} & \boldsymbol{x}\end{bmatrix}^T \boldsymbol{H}^{-1} \begin{bmatrix}\boldsymbol{1} & \boldsymbol{x}\end{bmatrix}\right) - \frac{1}{2}\tau\boldsymbol{y}^T\boldsymbol{W}_x\boldsymbol{y} \end{aligned} \tag{33}$$

where

$$\boldsymbol{H} = \lambda\boldsymbol{K} + \boldsymbol{I}_n \tag{34}$$

$$\boldsymbol{W}_x = \boldsymbol{H}^{-1} - \boldsymbol{H}^{-1}\begin{bmatrix}\boldsymbol{1} & \boldsymbol{x}\end{bmatrix}\left(\begin{bmatrix}\boldsymbol{1} & \boldsymbol{x}\end{bmatrix}^T \boldsymbol{H}^{-1}\begin{bmatrix}\boldsymbol{1} & \boldsymbol{x}\end{bmatrix}\right)^{-1}\begin{bmatrix}\boldsymbol{1} & \boldsymbol{x}\end{bmatrix}^T\boldsymbol{H}^{-1} \tag{35}$$

The term $\boldsymbol{W}_x$ is an $n \times n$ matrix. Noticing $\boldsymbol{x}$ is a zero-centred vector, we deduced that the following term in Equation 33 involves the genetic variance of $\boldsymbol{x}$.

$$\begin{aligned} \det\left(\begin{bmatrix}\mathbf{1} & \boldsymbol{x}\end{bmatrix}^T \begin{bmatrix}\mathbf{1} & \boldsymbol{x}\end{bmatrix}\right) &= \begin{vmatrix} n & \sum_{i=1}^n x_i \\ \sum_{i=1}^n x_i & \sum_{i=1}^n x_i^2 \end{vmatrix} = \begin{vmatrix} n & 0 \\ 0 & \sum_{i=1}^n x_i^2 \end{vmatrix} \\ &= n(n-1)\sum_{i=1}^n \frac{x_i^2}{n-1} = n(n-1)\,\mathrm{Var}(\boldsymbol{x}) \end{aligned} \tag{36}$$

Therefore, the target function (Equation 33) does not exist when $\boldsymbol{x}$ is a constant vector [$\mathrm{Var}(\boldsymbol{x}) = 0$, when the explanatory allele is present or absent in all genomes] because the function takes a logarithm of the determinant (Equation 36). This is a limitation in our LMMs: REML parameter estimates only exist when the explanatory allele has a frequency of neither zero nor one.

Provided existence of the target function, Zhou and Stephens pointed out that this function is maximised at the scalar [4]

$$\hat{\tau} = \frac{n-2}{\boldsymbol{y}^T \boldsymbol{W}_x \boldsymbol{y}} \tag{37}$$

assuming the parameter $\lambda$ is known. This equation immediately indicates another limitation in our method – $\hat{\tau}$ does not exist when the response allele is absent in all genomes (that is, $\boldsymbol{y} = \mathbf{0}$).

Putting Equations 37 and 36 back into Equation 33, we show the residual target function for REML estimates:

$$\begin{aligned} l_{r1}(\lambda; \boldsymbol{y}, \boldsymbol{x}, \boldsymbol{K}) &= \frac{1}{2}\left[(n-2)\log\frac{n-2}{2\pi} + \log\frac{n(n-1)}{\det \boldsymbol{H}} - (n-2)\right] \\ &+ \frac{1}{2}\left[\log\frac{\mathrm{Var}(\boldsymbol{x})}{\det\left(\begin{bmatrix}\mathbf{1} & \boldsymbol{x}\end{bmatrix}^T \boldsymbol{H}^{-1}\begin{bmatrix}\mathbf{1} & \boldsymbol{x}\end{bmatrix}\right)} - (n-2)\log\left(\boldsymbol{y}^T \boldsymbol{W}_x \boldsymbol{y}\right)\right] \end{aligned} \tag{38}$$

which consists of a constant term (within the first pair of square braces) and a variant term (within the second pair of square braces). Then we can determine the REML estimate of $\lambda$ for the Model 25 through

$$\hat{\lambda}_{r1} = \operatorname{argmax} l_{r1}(\lambda; \boldsymbol{y}, \boldsymbol{x}, \boldsymbol{K}) \tag{39}$$

Next, GEMMA uses a generalised least-square (GLS) approach to estimate parameters of fixed effects in the LMM [11]. Following the algebra by authors of GEMMA, we have derived the following GLS estimator and variance of $\beta$ given REML estimates of both variance components (i.e., structural and environmental) explaining the response pattern $\boldsymbol{y}$ [4].

$$\hat{\beta} = \left(\boldsymbol{x}^T \boldsymbol{W}_1 \boldsymbol{x}\right)^{-1} \boldsymbol{x}^T \boldsymbol{W}_1 \boldsymbol{y} \tag{40}$$

$$\mathrm{Var}(\hat{\beta}) = \frac{1}{n-2} \cdot \frac{\boldsymbol{y}^T \boldsymbol{W}_x \boldsymbol{y}}{\boldsymbol{x}^T \boldsymbol{W}_1 \boldsymbol{x}} \tag{41}$$

where $\boldsymbol{W}_1 = \boldsymbol{H}^{-1} - \boldsymbol{H}^{-1}\mathbf{1}\left(\mathbf{1}^T \boldsymbol{H}^{-1}\mathbf{1}\right)^{-1}\mathbf{1}^T \boldsymbol{H}^{-1}$ and it is an $n \times n$ matrix.

##### 3.1.10 Hypothesis tests for the fixed effect

The null hypothesis for our LMMs to be tested for is $\beta = 0$ while the alternative hypothesis is $\beta \neq 0$. The LMM defined in Equation 25 becomes $\boldsymbol{y} = \mathbf{1}\alpha + \boldsymbol{C}_L\boldsymbol{\gamma} + \boldsymbol{\varepsilon}$ under the null hypothesis.

Likelihood-ratio tests are invalid in our approach for comparing LMMs under the null and alternative hypotheses using the logarithms of their residual likelihood functions because they differ in fixed effects, that is, with or without the term $\boldsymbol{x}\beta$ besides the constant fixed effect $\mathbf{1}\alpha$. Instead, a Wald test is implemented in GEMMA to test for the null hypothesis. Specifically, the test statistic follows an F distribution when the null hypothesis is true and thereby a p-value is calculated [4].

$$F = \frac{\hat{\beta}^2}{\mathrm{Var}(\hat{\beta})} \sim F(1, n-2) \tag{42}$$

#### 3.2 Assessment of structural random effects

This step determines whether sample projections on an axis can explain the presence-absence of an allele as a structural random effect under a given significance level (namely, a maximum type-1 error rate or false-positive rate). According to the assumption of structural random effects for the Model 25, the effects follow a multivariate normal distribution (Equation 26). However, being different to the fixed effects $\mathbf{1}\alpha + \boldsymbol{x}\beta$, explicit element values in the vector $\boldsymbol{\gamma} = [\gamma_1, \cdots, \gamma_n, \cdots, \gamma_L]^T$ are unobservable in an LMM, even though we can estimate the relevant parameters $\lambda$ and $\tau$. Earle, Wu, et al showed that we can use the posterior distribution of $\boldsymbol{\gamma}$ under the null LMM to test for if $\gamma_i$ $(1 \leqslant i \leqslant L)$ significantly differs from zero [6]. In this section, we revise their algebra for higher stringency and accuracy.

##### 3.2.1 Posterior distribution of structural random effects

Given the null LMM,

$$\boldsymbol{y} = \mathbf{1}\alpha + \boldsymbol{C}_L\boldsymbol{\gamma} + \boldsymbol{\varepsilon} \tag{43}$$

we can rewrite it into an equivalent form

$$\boldsymbol{y} - \mathbf{1}\alpha = \boldsymbol{C}_L\boldsymbol{\gamma} + \boldsymbol{\varepsilon} \tag{44}$$

where $\boldsymbol{\gamma} \sim MVN_L\left(\mathbf{0}, \lambda\tau^{-1}L^{-1}\boldsymbol{I}_L\right)$ and $\boldsymbol{\varepsilon} \sim MVN_n\left(\mathbf{0}, \tau^{-1}\boldsymbol{I}_n\right)$. Let $\boldsymbol{z} = \boldsymbol{y} - \mathbf{1}\alpha$, we have an ordinary model of multiple regression:

$$\boldsymbol{z} = \boldsymbol{C}_L\boldsymbol{\gamma} + \boldsymbol{\varepsilon} \tag{45}$$

Given REML estimates of $\lambda$ and $\tau$ under the null model (Equation 43), prior distributions of $\boldsymbol{\gamma}$ and $\boldsymbol{\varepsilon}$ are determined:

$$\boldsymbol{\gamma} \sim MVN_L\left(\mathbf{0}, \hat{\lambda}\hat{\tau}^{-1}L^{-1}\boldsymbol{I}_L\right) \tag{46}$$

$$\boldsymbol{\varepsilon} \sim MVN_n\left(\mathbf{0}, \hat{\tau}^{-1}\boldsymbol{I}_n\right) \tag{47}$$

Supposing that the true variance-covariance matrix of the residual error $\boldsymbol{\varepsilon}$ is known and it equals $\hat{\tau}^{-1}\boldsymbol{I}_n$, we can deduce that the posterior distribution of $\boldsymbol{\gamma}$ is a multivariate normal distribution with a mean vector $\boldsymbol{\mu}$ and a covariance matrix $\boldsymbol{\Sigma}$ determined by the following procedure (cf. Theorem 11.45, Equations 11.60 and 11.61 on page 327 of Kendall's book [12], when $\sigma^2 = \tau^{-1}$ for both equations).

Let $\boldsymbol{w} = \hat{\lambda}\hat{\tau}^{-1}L^{-1}\boldsymbol{I}_L$. Note that $\boldsymbol{I}_L^{-1} = \boldsymbol{I}_L$, then $\boldsymbol{w}^{-1} = \hat{\lambda}^{-1}\hat{\tau}L\boldsymbol{I}_L$. According to Kendall's equation 11.60 [12], we derived that

$$\begin{aligned}
\boldsymbol{\mu} = E(\boldsymbol{\gamma}|\boldsymbol{z}) &= \left(\boldsymbol{w}^{-1} + \hat{\tau}\boldsymbol{C}_L^T\boldsymbol{C}_L\right)^{-1}\left(\boldsymbol{w}^{-1}\cdot\mathbf{0} + \hat{\tau}\boldsymbol{C}_L^T\boldsymbol{z}\right) = \hat{\tau}\left(\boldsymbol{w}^{-1} + \hat{\tau}\boldsymbol{C}_L^T\boldsymbol{C}_L\right)^{-1}\boldsymbol{C}_L^T\boldsymbol{z} \\
&= \left(L\hat{\lambda}^{-1}\boldsymbol{I}_L + \boldsymbol{C}_L^T\boldsymbol{C}_L\right)^{-1}\boldsymbol{C}_L^T(\boldsymbol{y} - \mathbf{1}\alpha) \\
&= \left(L\hat{\lambda}^{-1}\boldsymbol{I}_L + \boldsymbol{C}_L^T\boldsymbol{C}_L\right)^{-1}\boldsymbol{C}_L^T\boldsymbol{y} - \alpha\left(L\hat{\lambda}^{-1}\boldsymbol{I}_L + \boldsymbol{C}_L^T\boldsymbol{C}_L\right)^{-1}\boldsymbol{C}_L^T\mathbf{1}
\end{aligned} \tag{48}$$

On one hand, we have shown a rotation transformation $\boldsymbol{C}_L = \boldsymbol{S}\boldsymbol{Q}$ in Section 3.1.7, then

$$\boldsymbol{C}_L^T\mathbf{1} = (\boldsymbol{S}\boldsymbol{Q})^T\mathbf{1} = \boldsymbol{Q}^T\boldsymbol{S}^T\mathbf{1} \tag{49}$$

because column sums of the $n \times L$ centred genotype matrix $\boldsymbol{S}$ are zeros, we have

$$\boldsymbol{S}^T\mathbf{1} = \begin{bmatrix} s_{\cdot 1} \\ \vdots \\ s_{\cdot L} \end{bmatrix} = \mathbf{0}_{L\times 1} \tag{50}$$

where $s_{\cdot j} = \sum_{i=1}^{n} s_{ij}, 1 \leqslant j \leqslant L$, is the sum of elements in the $j$-th column. Therefore, the second term of Equation 48 is cancelled and we have the posterior mean

$$\boldsymbol{\mu} = \left(\boldsymbol{C}_L^T\boldsymbol{C}_L + L\hat{\lambda}^{-1}\boldsymbol{I}_L\right)^{-1}\boldsymbol{C}_L^T\boldsymbol{y} \tag{51}$$

which equals the ridge estimator of the coefficient vector $\boldsymbol{\eta}$ in the linear model

$$\boldsymbol{y} = \boldsymbol{C}_L\boldsymbol{\eta} + \boldsymbol{e} \text{ where } \boldsymbol{e} \sim MVN_n\left(\mathbf{0}, \sigma^2\boldsymbol{I}_n\right) \tag{52}$$

given the ridge parameter $k = L/\hat{\lambda}$. Consequently, the intercept term $\mathbf{1}\alpha$ in our LMM does not affect the posterior mean $\boldsymbol{\mu} = E(\boldsymbol{\gamma}|\boldsymbol{z})$ at all, which is reasonable as it only reveals the relative scale of observations.

On the other hand, since $\boldsymbol{C}_L = \boldsymbol{P\Sigma}$, $\boldsymbol{P}^T\boldsymbol{P} = \boldsymbol{I}_n$ and the diagonal matrix $\boldsymbol{\Sigma}^T = \boldsymbol{\Sigma}$, we show

$$\begin{aligned} \boldsymbol{C}_L^T\boldsymbol{C}_L &= (\boldsymbol{P\Sigma})^T\boldsymbol{P\Sigma} = \boldsymbol{\Sigma}^T\boldsymbol{P}^T\boldsymbol{P\Sigma} = \boldsymbol{\Sigma}^T\boldsymbol{\Sigma} \\ &= \begin{bmatrix} \boldsymbol{D}_{r\times r} & \boldsymbol{O}_{r\times(n-r)} \\ \boldsymbol{O}_{(L-r)\times r} & \boldsymbol{O}_{(L-r)\times(n-r)} \end{bmatrix} \begin{bmatrix} \boldsymbol{D}_{r\times r} & \boldsymbol{O}_{r\times(L-r)} \\ \boldsymbol{O}_{(n-r)\times r} & \boldsymbol{O}_{(n-r)\times(L-r)} \end{bmatrix} \\ &= \begin{bmatrix} \boldsymbol{D}_{r\times r}^2 & \boldsymbol{O}_{r\times(L-r)} \\ \boldsymbol{O}_{(L-r)\times r} & \boldsymbol{O}_{(L-r)\times(L-r)} \end{bmatrix}_{L\times L} = \operatorname{diag}\left(\lambda_1, \cdots, \lambda_r, 0, \cdots, 0\right) \end{aligned} \tag{53}$$

Let $\boldsymbol{\Lambda}_{L\times L}$ be the diagonal matrix $= \operatorname{diag}\left(\lambda_1, \cdots, \lambda_r, 0, \cdots, 0\right)$ of eigenvalues, we can simplify the Formula 51 into

$$
\begin{aligned}
\boldsymbol{\mu} &= \left(L\hat{\lambda}^{-1}\boldsymbol{I}_L + \boldsymbol{C}_L^T\boldsymbol{C}_L\right)^{-1}\boldsymbol{C}_L^T\boldsymbol{y} \\
&= \left(L\hat{\lambda}^{-1}\boldsymbol{I}_L + \boldsymbol{\Lambda}\right)^{-1}\boldsymbol{C}_L^T\boldsymbol{y} \\
&= \text{diag}\left[\left(L\hat{\lambda}^{-1}+\lambda_1\right)^{-1},\cdots,\left(L\hat{\lambda}^{-1}+\lambda_r\right)^{-1},0,\cdots,0\right]\boldsymbol{C}_L^T\boldsymbol{y} \\
&= \begin{bmatrix} \left(L\hat{\lambda}^{-1}+\lambda_1\right)^{-1}\boldsymbol{c}_1^T \\ \vdots \\ \left(L\hat{\lambda}^{-1}+\lambda_r\right)^{-1}\boldsymbol{c}_r^T \\ \mathbf{0} \\ \vdots \\ \mathbf{0} \end{bmatrix}\boldsymbol{y}
\end{aligned} \tag{54}
$$

Hence, the $i$-th ($i = 1,\cdots,r$) element of $\boldsymbol{\mu}$ is

$$
\boldsymbol{\mu}_i = \left(L\hat{\lambda}^{-1}+\lambda_i\right)^{-1}\boldsymbol{c}_i^T\boldsymbol{y} = \left(L\hat{\lambda}^{-1}+\lambda_i\right)^{-1}\sum_{j=1}^{n} c_{ji}y_j, 1 \leqslant i \leqslant r \tag{55}
$$

which is the $i$-th posterior mean of $\gamma_i$ given $\boldsymbol{z}$ (or $\boldsymbol{y}$) and $\boldsymbol{C}_L$. This equation illustrates that we can use $\boldsymbol{C}_r$ instead of $\boldsymbol{C}_L$ to capture all genetic variations underlying population structure.

Similarly, we can deduce the variance-covariance matrix of $\boldsymbol{\gamma}|\boldsymbol{z}$ through Kendall's equation 11.61 [12].

$$
\boldsymbol{\Delta} = \text{Cov}(\boldsymbol{\gamma}|\boldsymbol{z}) = \left(\boldsymbol{w}^{-1} + \hat{\tau}\boldsymbol{C}_L^T\boldsymbol{C}_L\right)^{-1} = \hat{\tau}^{-1}\left(L\hat{\lambda}^{-1}\boldsymbol{I}_L + \boldsymbol{C}_L^T\boldsymbol{C}_L\right)^{-1} \tag{56}
$$

Knowing Equation 53, we show that this $\boldsymbol{\Delta}$ is an $L \times L$ diagonal matrix:

$$
\begin{aligned}
\boldsymbol{\Delta} &= \hat{\tau}^{-1}\left(L\hat{\lambda}^{-1}\boldsymbol{I}_L + \boldsymbol{C}_L^T\boldsymbol{C}_L\right)^{-1} = \hat{\tau}^{-1}\left(L\hat{\lambda}^{-1}\boldsymbol{I}_L + \boldsymbol{\Lambda}\right)^{-1} \\
&= \hat{\tau}^{-1}\,\text{diag}\left[\left(L\hat{\lambda}^{-1}+\lambda_1\right),\cdots,\left(L\hat{\lambda}^{-1}+\lambda_r\right),\left(L\hat{\lambda}^{-1}+0\right),\cdots,\left(L\hat{\lambda}^{-1}+0\right)\right]^{-1} \\
&= \hat{\tau}^{-1}\,\text{diag}\left[\left(L\hat{\lambda}^{-1}+\lambda_1\right)^{-1},\cdots,\left(L\hat{\lambda}^{-1}+\lambda_r\right)^{-1},\hat{\lambda}/L,\cdots,\hat{\lambda}/L\right]
\end{aligned} \tag{57}
$$

This is what we can expect for genome projections, which are mutually independent, that their effect sizes to the response $\boldsymbol{y}$ are independent as well. Put Equations 51 and 56 together, we have the posterior distribution of $\boldsymbol{\gamma}$ based on the full form of SVD.

$$
\begin{aligned}
&\boldsymbol{\gamma}|\boldsymbol{z} = \boldsymbol{\gamma}|\boldsymbol{y} \sim MVN_n(\boldsymbol{\mu}, \boldsymbol{\Delta}) \\
&\text{where } \boldsymbol{\mu} = \left(\boldsymbol{C}_L^T\boldsymbol{C}_L + L\hat{\lambda}^{-1}\boldsymbol{I}_L\right)^{-1}\boldsymbol{C}_L^T\boldsymbol{y} \\
&\text{and } \boldsymbol{\Delta} = \hat{\tau}^{-1}\left(L\hat{\lambda}^{-1}\boldsymbol{I}_L + \boldsymbol{C}_L^T\boldsymbol{C}_L\right)^{-1}
\end{aligned}
\tag{58}
$$

##### 3.2.2 Bayesian chi-square tests of structural random effects

Let $\varphi_i = \gamma_i|\boldsymbol{y}$ for Equation 58, where $i = 1, \cdots, L$, we are interested in testing for the null hypothesis $H_0 : \varphi_i = 0$ versus an alternative hypothesis $H_1 : \varphi_i \neq 0$. We consider genome projections along the $i$-th PC contributes to the presence-absence status in $\boldsymbol{y}$ if $H_0$ is rejected under a given significance level.

Since $\varphi_i$ is a member variable participating in the multivariate normal distribution (Equation 58), it follows a univariate normal distribution:

$$
\gamma_i|\boldsymbol{y} = \varphi_i \sim \mathrm{N}(\mu_i, \Delta_{ii}) \tag{59}
$$

where $\Delta_{ii}$ is the $i$-th diagonal element of the matrix $\boldsymbol{\Sigma}$. Therefore, we can construct a random variable (denoted as $w_i$) that follows a chi-square distribution of one degree of freedom from the distribution of $\varphi_i$ using the connection between a normal distribution and a chi-square distribution.

$$
w_i = \left(\frac{\varphi_i - \mu_i}{\sqrt{\Delta_{ii}}}\right)^2 = \frac{(\varphi_i - \mu_i)^2}{\Delta_{ii}} \sim \chi^2(1) \tag{60}
$$

Hence the null hypothesis for the posterior distribution (Equation 58), $\gamma_i|\boldsymbol{y} = \varphi_i = 0$, is equivalently converted into an observation of the chi-square distribution:

$$
w_i|_{\varphi_i=0} = \frac{(0-\mu_i)^2}{\Delta_{ii}} = \frac{\mu_i^2}{\Delta_{ii}} \tag{61}
$$

This is the statistic drawn from the population of $\chi^2(1)$ to test for the null hypothesis $\varphi_i = 0$ versus the alternative $\varphi_i \neq 0$, and it relies on parameter estimates $\hat{\lambda}$ and $\hat{\tau}$ of the null LMM as well as genome projections. Assuming a confidence level of $p_0$ ($0 < p_0 < 1$), we can consider the event of observing $w_i|_{\varphi_i=0} \geqslant \mu_i^2/\Delta_{ii}$ as "impossible" and reject the null hypothesis if the upper-tail probability $P(w_i|_{\varphi_i=0} \geqslant \mu_i^2/\Delta_{ii}) \leqslant p_0$ of $\chi^2(1)$ – it is unlikely the null distribution of $w_i$ holds. By convention, the threshold $p_0$ for confidence may be 0.05, 0.01, and so forth.

Furthermore, it is worth noting that this hypothesis test determines whether projections of samples along a specific PC explain allelic presence-absence status in the response vector $\boldsymbol{y}$. Hence it may report a PC that does not significantly contribute to the presence of alleles but the absence. In other words, a PC may show a significant positive association or a negative association with $\boldsymbol{y}$.

#### 3.3 Scoring evidence of physical linkage

In this section, we describe a scoring scheme for enriching allele pairs that are physically linked in HGT. The identification of co-localised alleles is a particular utility of the association result. The final score applies to each pair of explanatory allele and response allele. In other words, it weights each directed edge given an association network of alleles. We have developed this scheme via taking into account the direction of each significant association and characteristics in measurements of pairwise physical distances between alleles. As such, the overall score (denoted by $s$ hereafter) is comprised of two components, which are explained in the following contents in details.

##### 3.3.1 Score for association status

For each pair of explanatory and response alleles, the first component of the overall score is a score for association status in terms of its orientation (positive or negative) and statistical significance. Let $s_a$ denote the association score, then $s_a$ is determined using a decision tree shown in Figure s17.

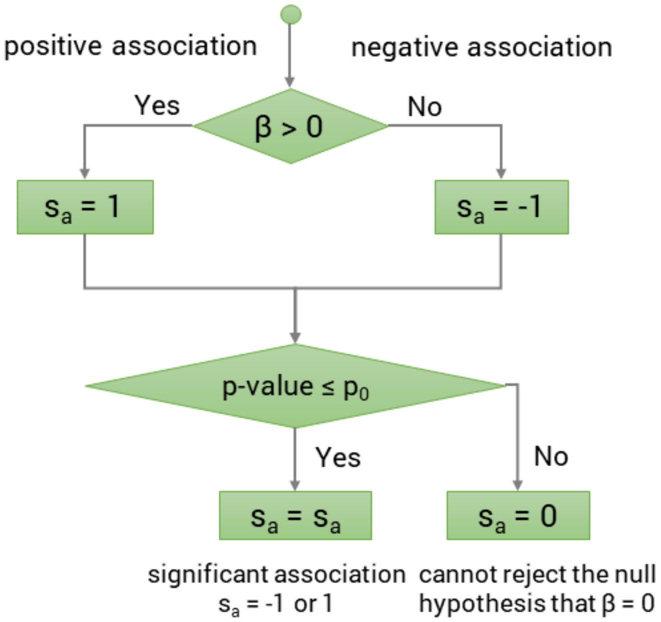

**Figure s17: A decision tree determining the association score $s_a$.** The parameter $p_0$ is the threshold of p-values for significance. By convention, it may be 0.05 in practice, corresponding to a type-1 error rate of 5%.

Specifically, the association score $s_a$ is a discrete variable equalling -1 (evidence against physical linkage), 0 (insufficient evidence for a decision), or 1 (evidence supporting physical linkage). The tree is designed under the consideration that a significant positive association is strong evidence for the co-localisation of alleles, and a significant negative association is evidence against the presence of allelic co-localisation. Nonetheless, since the association status tested for using LMMs are directed and confounded by the contrast of allele distributions (e.g., how many overlaps and mismatches in distributions of each pair of alleles), there may be significant associations where $\beta < 0$ but alleles are actually co-localised in some genomes. These cases can be considered as co-localisation signals that are too weak to be detected due to a high level of noise caused by co-occurrence of the same alleles which are however physically unlinked in the same bacterial collection.

##### 3.3.2 Score for allelic physical distances

Since physical distances between acquired alleles are unlikely to conserve across bacterial linages when individual alleles are randomly acquired and inserted into bacterial genomes, consistency in the physical distances found in phylogenetically distant bacterial genomes provides us with the second layer of evidence for the inference of physical linkage. In addition, the network of positive associations between alleles may contain edges resulted from transfer dependency (where the horizontal transfer of one MGE relies on the presence of another) or particular combinations of alleles whose distribution contrast leads to a positive association only computationally. Taken together, we leverage physical distances (physical evidence) to filter positive associations (statistical evidence) for those supporting the existence of physical linkage.

Nonetheless, we cannot directly incorporate the distances into an LMM as a covariate by far because it is highly correlated with the presence-absence of the explanatory allele and is confounded by the same population structure. For instance, it is evident that the distance is unmeasurable when the explanatory allele is absent (the same to the response allele as well), and the consistency in distances may be resulted from a phenomenon called identity-by-descent (IBD), where the same genomic structure is passed down to descendants through clonal reproduction. In evolution, the IBD gets lost or weaken in some bacteria in the same clade because of gene-loss events, insertion of MGEs (A common example is the disruption of genomic structures by the acquisition of insertion sequences), genomic rearrangement, and so forth, causing a spurious HGcoT signal in the distribution of allelic co-occurrence events. Therefore, we designed a decision tree to score the consistency in APDs (Figure s18) while taking into account the probability of bacterial genomes to display consistent distances. This method enables us to integrate the association score and make a concise scoring scheme.

Assuming $m(\geqslant 2)$ valid physical distances $d_1, \cdots, d_m$ are measured in $m$ genomes for a pair of alleles X and Y, the decision tree works as follows.

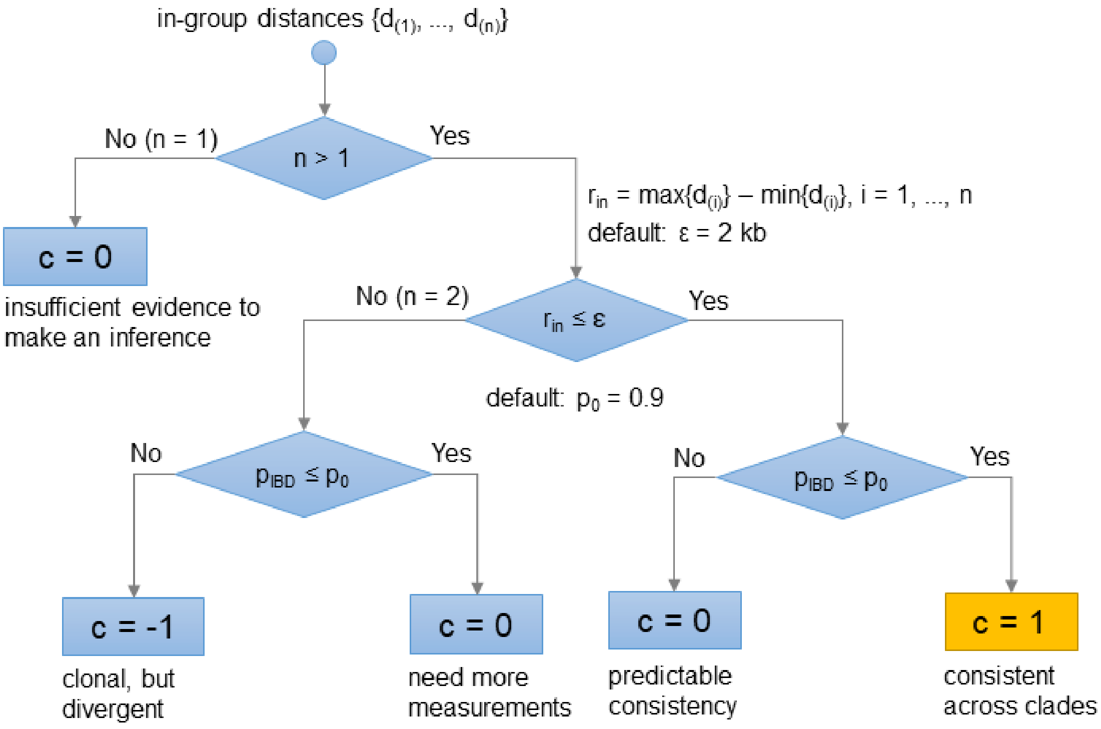

**Figure s18: A decision tree determining the consistency score** $c$. IBD: identity-by-descendent in terms of the presence-absence of consistent physical distances in bacterial genomes. $r_{in}$: range of in-group distances. $\varepsilon$: a threshold of $r_{in}$. The in-group distances are considered as consistent when $r_{in} < \varepsilon$. It may be twice the maximum error (with a unit of bp) to be tolerated for calling the distance measurements accurate (i.e., the error tolerance). $p_{IBD}$: an estimate of the probability that the presence of consistent physical distances in bacterial genomes is due to IBD. $p_0$: an upper bound for $p_{IBD}$, above which the consistency in the distances is considered as IBD.

1. Define in-group distance measurements as those within (inclusive) the range $Q1 - 1.5IQR \leqslant d_i \leqslant Q3 + 1.5IQR(1 \leqslant i \leqslant m)$, where Q1 and Q3 denote the first and third quantiles (namely, the 25th and 75th percentiles) of all the distances, respectively, and IQR is an abbreviation of the interquartile range (equalling $Q3 - Q1$) of distances. Outlier distance measurements are determined accordingly. This grouping of distances enables us to evade erroneous decisions driven by a few outlier distances that may be caused by poor assembly quality, recombination events, etc.

2. Assuming there are $n$ ($n \leqslant m$) in-group physical distances measured between the alleles X and Y in $n$ bacterial genomes (hence there is only a single distance measurement per genome), let $d_{(i)}$ ($1 \leqslant i \leqslant n$) represent the distance measurement in the $i$-th genome and define the range of in-group distances as $r_{in} = max\{d_{(i)}\} - min\{d_{(i)}\}$. Herein, we assume $n \geqslant 2$ for the rest of steps for simplicity of our description.

3. The in-group distances are considered consistent if their range $r_{in} \leqslant \varepsilon$ where $\varepsilon$ (bp) is a user-specified upper bound for distance ranges. We suggest users to set $\varepsilon$

as twice the error tolerance for distance measurements to simplify explanations to results as this is the maximum difference we can expect to see under this accuracy level when the true distances are the same in all of $n$ genomes.

4. To determine if the presence of consistent in-group distances is due to IBD, we obtain a binary vector as a “trait” for all genomes where one is assigned to the $n$ genomes and zero to the others to denote the presence and absence of in-group distances, respectively. Then we reconstruct the presence-absence state of the same/similar distance in the most-recent common ancestor of all the $n$ genomes using the ace function in the R package ape [13] under an all-rates-equal model for state transitions of a discrete trait in the genomes. Based on the outcome of this function, we consider the presence of consistent distances as a result of IBD if the empirical Bayesian posterior probability of the ancestral state of “presence” exceeds a pre-specified threshold $p_0$ (we use 0.9 by default for this parameter). The consistency score $c$ is determined accordingly.

5. The same ancestral state reconstruction applies to inconsistent in-group distances as well. It can be understood as an evaluation of the tendency in the $n$ genomes to show consistent physical distances following clonal reproduction. We consider inconsistency in the distances when the tendency is strong as evidence against physical linkage (hence $c$ equals -1), whereas it is unsurprised to see inconsistency in the distances when the tendency is weak or absent (hence $c$ equals zero).

For this scoring procedure, a user may provide an ML tree estimated using an external program as an input. GeneMates generates a neighbour-joining tree from Euclidean distances between genome projections for ancestral state reconstruction when such a pre-specified tree is absent. Comparing to the ML tree, the projection-based neighbour-joining tree does not depend on models or assumptions, it however may be less accurate than the ML tree as it is built on less SNPs (only biallelic cgSNPs) than is the ML tree (usually constructed from SNPs conserved in 99% of bacterial genomes).

##### 3.3.3 Overall score

The final score for the physical linkage between a specific pair of alleles is an integration of both the association score $s_a$ and the consistency score $c$. In addition, we need to consider the measurability of APDs in genomes where both alleles are co-occurring because it is positively associated with representability of the consistency score, frequency of allelic co-localisation and quality of genome assemblies. Specifically, we define the in-group measurability ($m_{in}$) as the proportion of genomes where a pair of alleles are co-occurring have the APDs reliably measured and the distance is not an outlier. Accordingly, $0 \leqslant m_{in} \leqslant 1$. Then the overall linkage score is defined as

$$s = s_a + m_{in}c \tag{62}$$

where the term $s_a$ is the association component of $s$ and $m_{in}c$ is the distance component. Note that the measurability can be considered as a weight for the consistency score. In particular, we define a distance score $s_d = m_{in}c$ to simplify Equation 62. Evidently, $s$ is a continuous variable and $-2 \leqslant s \leqslant 2$.

Assuming perfect measurability (namely, $m_{in} = 1$), since both $s_a$ and $c$ have three levels (-1, 0, 1) each, there are five levels of $s$ as shown in Table s13:

**Table s13: Overall scores given association scores and distance scores under the assumption of perfect distance measurability ($m_{in} = 1$).**

| Scores | | $s_d$ 1 | 0 | -1 |
|---|---|---|---|---|
| | **1** | 2 | 1 | 0 |
| $s_a$ | **0** | 1 | 0 | -1 |
| | **-1** | 0 | -1 | -2 |

And these overall scores can be interpreted with the following list:

- 2: the physical linkage is well supported by both association analysis and APDs;
- 1: the linkage is supported by either association analysis or APDs;
- 0: we cannot determine whether a pair of alleles are physically linked or not;
- -1: there is weak evidence opposing the presence of a physical linkage;
- -2: there is strong evidence against the presence of a physical linkage.

An advantage in using the product of measurability and the distance score is that it does not apply a hard and often arbitrary cut-off to the measurability for filtering edges in the association network. As a result, the overall scores retain more information for investigation than do the scores filtered for a certain measurability level. In practice, the overall score can be mapped to the colour depth for edges in an association network so that we can identify clusters of co-localised alleles at different levels of measurability. Some studies may use 0.5 as a cut-off for $m_{in}$ to filter out scores that are not sufficiently representative.

#### 3.4 Further discussion

In this section, we show connections between our association analysis and other methods. Furthermore, we explain the fixed effect of the explanatory variable. Finally, we discuss limitations and further improvements of the approach to detection of HGcoT.

##### 3.4.1 Model equivalence

Here, we demonstrate equivalence of our LMM to LMMs fitted by GEMMA, BugWAS and EMMA [14] using affine transformations.

**Equivalence to the standard LMM of GEMMA** Let $\boldsymbol{u} = \boldsymbol{C}_L\boldsymbol{\gamma}$ denote the term of random structural effect, we used an affine transformation of multivariate normal distributions to prove the equivalence between Model 25 and the standard LMM fitted in GEMMA [4]. Specifically, given $\boldsymbol{u} = \boldsymbol{C}_L\boldsymbol{\gamma}$, Equation 26 and $\boldsymbol{C}_L = \boldsymbol{SQ} = \boldsymbol{P\Sigma}$, the affine transformation shows

$$\boldsymbol{u} \sim MVN_n\left[\boldsymbol{0}, \boldsymbol{C}_L\left(\lambda\tau^{-1}L^{-1}\boldsymbol{I}_L\right)\boldsymbol{C}_L^T\right] \tag{63}$$

and

$$\begin{aligned}\boldsymbol{C}_L\left(\lambda\tau^{-1}L^{-1}\boldsymbol{I}_L\right)\boldsymbol{C}_L^T &= \lambda\tau^{-1}\frac{\boldsymbol{C}_L\boldsymbol{C}_L^T}{L} = \lambda\tau^{-1}\frac{(\boldsymbol{P\Sigma})(\boldsymbol{SQ})^T}{L}\\ &= \lambda\tau^{-1}\frac{\left(\boldsymbol{P\Sigma Q}^T\right)\boldsymbol{S}^T}{L} = \lambda\tau^{-1}\frac{\boldsymbol{SS}^T}{L} = \lambda\tau^{-1}\boldsymbol{K}\end{aligned} \tag{64}$$

This equivalence also applies to Model 31, which follows the reduced form of SVD. Specifically, applying an affine transformation on the $n\times 1$ vector $\boldsymbol{u} = \boldsymbol{C}_r\boldsymbol{\gamma}_r$, we can restore the same variance-covariance matrix of the multivariate normal distribution in the GEMMA LMM.

$$\begin{aligned}\boldsymbol{C}_r\left(\lambda\tau^{-1}L^{-1}\boldsymbol{I}_r\right)\boldsymbol{C}_r^T &= \lambda\tau^{-1}\frac{\boldsymbol{C}_r\boldsymbol{C}_r^T}{L} = \lambda\tau^{-1}\frac{(\boldsymbol{UD})(\boldsymbol{SV})^T}{L}\\ &= \lambda\tau^{-1}\frac{\left(\boldsymbol{UDV}^T\right)\boldsymbol{S}^T}{L} = \lambda\tau^{-1}\frac{\boldsymbol{SS}^T}{L} = \lambda\tau^{-1}\boldsymbol{K}\end{aligned} \tag{65}$$

Taken together, our LMM is equivalent to the LMM in GEMMA:

$$\boldsymbol{y} = \boldsymbol{1}\alpha + \boldsymbol{x}\beta + \boldsymbol{C}_L\boldsymbol{\gamma} + \boldsymbol{\varepsilon} = \boldsymbol{1}\alpha + \boldsymbol{x}\beta + \boldsymbol{u} + \boldsymbol{\varepsilon} \tag{66}$$

$$\boldsymbol{u} \sim MVN_n\left(\boldsymbol{0}, \lambda\tau^{-1}\boldsymbol{K}\right) \tag{67}$$

$$\boldsymbol{\varepsilon} \sim MVN_n\left(\boldsymbol{0}, \tau^{-1}\boldsymbol{I}_n\right) \tag{68}$$

Hypotheses for $\beta$ to be tested for are the same between both models.

**Equivalence to the LMM of BugWAS** Our model is also equivalent to the LMM underlying the BugWAS method. More specifically, BugWAS estimates parameters of the LMM

$$\boldsymbol{y} = \mathbf{1}\alpha + \boldsymbol{x}\beta + \boldsymbol{S}\boldsymbol{\delta} + \boldsymbol{\varepsilon} \tag{69}$$

$$\boldsymbol{\delta} \sim MVN_L\left(\mathbf{0}, \lambda'\tau^{-1}\boldsymbol{I}_L\right) \tag{70}$$

$$\boldsymbol{\varepsilon} \sim MVN_n\left(\mathbf{0}, \tau^{-1}\boldsymbol{I}_n\right) \tag{71}$$

Herein the $L \times 1$ vector $\boldsymbol{\delta}$ represents additive background effects of cgSNPs. Let $\boldsymbol{u}' = \boldsymbol{S}\boldsymbol{\delta}$, using an affine transformation of Equation 70, we show

$$\boldsymbol{u}' \sim MVN_n\left[\mathbf{0}, \boldsymbol{S}\left(\lambda'\tau^{-1}\boldsymbol{I}_L\right)\boldsymbol{S}^T\right] \tag{72}$$

and we can calculate that

$$\boldsymbol{S}\left(\lambda'\tau^{-1}\boldsymbol{I}_L\right)\boldsymbol{S}^T = \lambda'\tau^{-1}L\frac{\boldsymbol{S}\boldsymbol{S}^T}{L} = L\lambda'\tau^{-1}\boldsymbol{K} \tag{73}$$

Particularly, when $\lambda' = \lambda/L$, $\boldsymbol{u}'$ follows the same distribution of $\boldsymbol{u}$ in our model.

$$\boldsymbol{u}' \sim MVN_n\left(\mathbf{0}, \lambda\tau^{-1}\boldsymbol{K}\right) \tag{74}$$

As such, the LMM of BugWAS becomes an equivalent form of our model when $\lambda' = \lambda/L$:

$$\boldsymbol{y} = \mathbf{1}\alpha + \boldsymbol{x}\beta + \boldsymbol{S}\boldsymbol{\delta} + \boldsymbol{\varepsilon} = \mathbf{1}\alpha + \boldsymbol{x}\beta + \boldsymbol{u}' + \boldsymbol{\varepsilon} \tag{75}$$

where $\boldsymbol{u}' \sim MVN_n\left(\mathbf{0}, \lambda\tau^{-1}\boldsymbol{K}\right)$. We thereby justified the feasibility in using GEMMA to estimate parameters for the BugWAS model so as to test for the association between a phenotype and the genotype of an explanatory locus.

**Equivalence between LMMs of GEMMA and EMMA** First, we demonstrate the equivalence between two forms of LMMs that are fitted using GEMMA. An equivalent form of Equation 66 in the GEMMA paper is:

$$\begin{aligned} \boldsymbol{y} &= \mathbf{1}\alpha + \boldsymbol{x}\beta + \boldsymbol{Z}\boldsymbol{u}_m + \boldsymbol{\varepsilon} \\ \boldsymbol{u}_m &\sim MVN_n\left(\mathbf{0}, \lambda\tau^{-1}\boldsymbol{K}_m\right) \\ \boldsymbol{\varepsilon} &\sim MVN_n\left(\mathbf{0}, \tau^{-1}\boldsymbol{I}_n\right) \end{aligned} \tag{76}$$

where $\boldsymbol{Z}$ is an $n \times m$ incidence matrix showing the membership of $n$ individuals in $m$ lineages/groups, and $\boldsymbol{K}_m$ is an $m \times m$ relatedness matrix between the lineages. In the simplest scenario of GWAS, $\boldsymbol{Z}$ can be an $n \times n$ identity matrix. According to supplementary materials of the GEMMA paper, there are two matrices playing an important role in parameter estimation:

$$\boldsymbol{G} = \boldsymbol{Z}\boldsymbol{K}_m\boldsymbol{Z}^T \tag{77}$$

$$\boldsymbol{H} = \lambda\boldsymbol{G} + \boldsymbol{I}_n \tag{78}$$

Using an affine transformation and defining $\boldsymbol{u} = \boldsymbol{Z}\boldsymbol{u}_m$ and $\boldsymbol{K} = \boldsymbol{Z}\boldsymbol{K}_m\boldsymbol{Z}^T$, the linear combination of elements in um follows a multivariate normal distribution:

$$\boldsymbol{u} = \boldsymbol{Z}\boldsymbol{u}_m \sim MVN_n\left[\boldsymbol{0}, \boldsymbol{Z}\left(\lambda\tau^{-1}\boldsymbol{K}_m,\right)\boldsymbol{Z}^T\right] \tag{79}$$

$$\boldsymbol{Z}\left(\lambda\tau^{-1}\boldsymbol{K}_m,\right)\boldsymbol{Z}^T = \lambda\tau^{-1}\left(\boldsymbol{Z}\boldsymbol{K}_m\boldsymbol{Z}^T\right) = \lambda\tau^{-1}\boldsymbol{K} \tag{80}$$

By definition, the $n \times n$ matrix $\boldsymbol{K} = \boldsymbol{Z}\boldsymbol{K}_m\boldsymbol{Z}^T$ describes the relatedness between these $n$ individuals. Substituting $\boldsymbol{Z}\boldsymbol{u}_m$ with $\boldsymbol{u}$ in Equation 76, we obtain Equation 66.

$$\boldsymbol{y} = \boldsymbol{1}\alpha + \boldsymbol{x}\beta + \boldsymbol{u} + \boldsymbol{\varepsilon} \tag{81}$$

where $\boldsymbol{u} \sim MVN_n\left(\boldsymbol{0}, \lambda\tau^{-1}\boldsymbol{K}\right)$ and $\boldsymbol{K} = \boldsymbol{Z}\boldsymbol{K}_m\boldsymbol{Z}^T$. Hence $\boldsymbol{G} = \boldsymbol{Z}\boldsymbol{K}_m\boldsymbol{Z}^T = \boldsymbol{K}$ and $\boldsymbol{H} = \lambda\boldsymbol{K} + \boldsymbol{I}_n$. Particularly, $\boldsymbol{K}_m = \boldsymbol{K}$ and both Models 66 and 76 become the same when $m = n$ and $\boldsymbol{Z} = \boldsymbol{I}_n$. However, we usually take the form of Equation 66 for LMMs in practice because $\boldsymbol{K}$ can be easily calculated using the formula $\boldsymbol{K} = (\boldsymbol{S}\boldsymbol{S}^T)/L$ while the calculation of $\boldsymbol{K}_m$ may be difficult.

On the other hand, we know that GEMMA is an improvement of EMMA. Specifically, EMMA works on the LMM [14]:

$$\boldsymbol{y} = \boldsymbol{X}\boldsymbol{\beta} + \boldsymbol{Z}\boldsymbol{g} + \boldsymbol{\varepsilon} \tag{82}$$

$$\boldsymbol{g} \sim MVN_n\left(\boldsymbol{0}, \sigma_g^2\boldsymbol{K}_m\right) \tag{83}$$

$$\boldsymbol{\varepsilon} \sim MVN_n\left(\boldsymbol{0}, \sigma_e^2\boldsymbol{I}_n\right) \tag{84}$$

where $\boldsymbol{X}\boldsymbol{\beta}$ includes an intercept and possible covariates. Herein we am interested in $\sigma_g^2$ and $\sigma_e^2$, which are known as variance components of random effects $\boldsymbol{g}$ and residuals $\boldsymbol{\varepsilon}$, respectively. According to the GEMMA paper, $\lambda$ is defined as the ratio of $\sigma_g^2$ on $\sigma_e^2$ (cf.

Section 3.1.8).

$$\lambda = \frac{\sigma_g^2}{\sigma_e^2} \tag{85}$$

Hence $\sigma_g^2 = \lambda\sigma_e^2$. Following the EMMA paper and Equation 80,

$$\boldsymbol{H}' = \boldsymbol{Z}\boldsymbol{K}_m\boldsymbol{Z}^T + \lambda^{-1}\boldsymbol{I}_n = \boldsymbol{K} + \lambda^{-1}\boldsymbol{I}_n = \lambda^{-1}(\lambda\boldsymbol{K} + \boldsymbol{I}_n) = \lambda^{-1}\boldsymbol{H} \tag{86}$$

Therefore, $\boldsymbol{H}'^{-1} = \lambda\boldsymbol{H}^{-1}$. Furthermore, the authors of EMMA illustrated that the full log-likelihood function of their model (Equation 82) is:

$$\begin{aligned}
l_F(\boldsymbol{y};\boldsymbol{\beta},\sigma_g,\lambda) &= -\frac{1}{2}\left[n\log\left(2\pi\sigma_g^2\right) + \log\det\boldsymbol{H}' + \frac{1}{\sigma_g^2}(\boldsymbol{y}-\boldsymbol{X\beta})^T\left(\boldsymbol{H}'\right)^{-1}(\boldsymbol{y}-\boldsymbol{X\beta})\right] \\
&= -\frac{1}{2}\left[n\log(2\pi) + n\log\left(\lambda\sigma_e^2\right) + \log\det\left(\lambda^{-1}\boldsymbol{H}\right) + \frac{1}{\lambda\sigma_e^2}(\boldsymbol{y}-\boldsymbol{X\beta})^T\left(\lambda^{-1}\boldsymbol{H}\right)^{-1}(\boldsymbol{y}-\boldsymbol{X\beta})\right] \\
&= -\frac{1}{2}\left[n\log(2\pi) + n\log\lambda + n\log\sigma_e^2 + \log\left(\lambda^{-n}\det\boldsymbol{H}\right) + \sigma_e^{-2}(\boldsymbol{y}-\boldsymbol{X\beta})^T\boldsymbol{H}^{-1}(\boldsymbol{y}-\boldsymbol{X\beta})\right] \\
&= -\frac{1}{2}\left[n\log(2\pi) + n\log\lambda + n\log\sigma_e^2 - n\log\lambda + \log\det\boldsymbol{H} + \sigma_e^{-2}(\boldsymbol{y}-\boldsymbol{X\beta})^T\boldsymbol{H}^{-1}(\boldsymbol{y}-\boldsymbol{X\beta})\right] \\
&= -\frac{1}{2}\left[n\log(2\pi) + n\log\sigma_e^2 + \log\det\boldsymbol{H} + \sigma_e^{-2}(\boldsymbol{y}-\boldsymbol{X\beta})^T\boldsymbol{H}^{-1}(\boldsymbol{y}-\boldsymbol{X\beta})\right]
\end{aligned} \tag{87}$$

Let $\tau = \sigma_e^{-2}$, then $\sigma_e^2 = \tau^{-1}$, hence we have

$$\begin{aligned}
l_F(\boldsymbol{y};\boldsymbol{\beta},\lambda,\tau) &= l_F(\boldsymbol{y};\boldsymbol{\beta},\sigma_g,\lambda) \\
&= -\frac{1}{2}\left[n\log(2\pi) - n\log\tau + \log\det\boldsymbol{H} + \tau(\boldsymbol{y}-\boldsymbol{X\beta})^T\boldsymbol{H}^{-1}(\boldsymbol{y}-\boldsymbol{X\beta})\right] \\
&= \frac{1}{2}\left[n\log\tau - n\log(2\pi) - \log\det\boldsymbol{H} - \tau(\boldsymbol{y}-\boldsymbol{X\beta})^T\boldsymbol{H}^{-1}(\boldsymbol{y}-\boldsymbol{X\beta})\right]
\end{aligned} \tag{88}$$

which is exactly the log-likelihood function of the standard LMM in GEMMA. In conclusion, both GEMMA and EMMA use the same LMM when $\lambda = \sigma_g^2/\sigma_e^2$ and $\tau = \sigma_e^{-2}$ (note that we do not require $m = n$). In other words, LMMs in Equations 25, 31, 66, 75, 76, and 82 are equivalent under aforementioned conditions. Moreover, they use the same relatedness matrix $\boldsymbol{K}$ in this case.

##### 3.4.2 Equivalent posterior distributions of structural random effects

In this section, we demonstrate equivalence between algebra calculating posterior distributions of structural random effects based on different LMMs.

**Equivalence to the posterior distribution derived for BugWAS** On one hand, since $\boldsymbol{C}_L^T\boldsymbol{C}_L = (\boldsymbol{SQ})^T\boldsymbol{SQ} = \boldsymbol{Q}^T\boldsymbol{S}^T\boldsymbol{SQ}$, $\boldsymbol{Q}^T\boldsymbol{Q} = \boldsymbol{QQ}^T = \boldsymbol{I}_L$, and $\boldsymbol{Q}^{-1} = \boldsymbol{Q}^T$, we rewrite the posterior mean (Equation 51) using the centred SNP matrix $\boldsymbol{S}$:

$$\begin{aligned}\boldsymbol{\mu} &= \left(L\hat{\lambda}^{-1}\boldsymbol{Q}^T\boldsymbol{Q}+\boldsymbol{Q}^T\boldsymbol{S}^T\boldsymbol{SQ}\right)^{-1}(\boldsymbol{SQ})^T\boldsymbol{y} = \left[\boldsymbol{Q}^T\left(L\hat{\lambda}^{-1}\boldsymbol{I}_L+\boldsymbol{S}^T\boldsymbol{S}\right)\boldsymbol{Q}\right]^{-1}\boldsymbol{Q}^T\boldsymbol{S}^T\boldsymbol{y}\\ &= \left[\boldsymbol{QQ}^T\left(L\hat{\lambda}^{-1}\boldsymbol{I}_L+\boldsymbol{S}^T\boldsymbol{S}\right)\boldsymbol{Q}\right]^{-1}\boldsymbol{S}^T\boldsymbol{y} = \boldsymbol{Q}^{-1}\left(L\hat{\lambda}^{-1}\boldsymbol{I}_L+\boldsymbol{S}^T\boldsymbol{S}\right)^{-1}\boldsymbol{S}^T\boldsymbol{y}\\ &= \boldsymbol{Q}^T\left(L\hat{\lambda}^{-1}\boldsymbol{I}_L+\boldsymbol{S}^T\boldsymbol{S}\right)^{-1}\boldsymbol{S}^T\boldsymbol{y}\end{aligned} \tag{89}$$

Similarly, the variance-covariance matrix (Equation 56) of the posterior distribution can be rewritten as

$$\begin{aligned}\boldsymbol{\Delta} &= \hat{\tau}^{-1}\left(\boldsymbol{C}_L^T\boldsymbol{C}_L+L\hat{\lambda}^{-1}\boldsymbol{I}_L\right)^{-1} = \hat{\tau}^{-1}\left(\boldsymbol{Q}^T\boldsymbol{S}^T\boldsymbol{SQ}+L\hat{\lambda}^{-1}\boldsymbol{Q}^T\boldsymbol{Q}\right)^{-1}\\ &= \hat{\tau}^{-1}\left[\boldsymbol{Q}^T\left(\boldsymbol{S}^T\boldsymbol{S}+L\hat{\lambda}^{-1}\boldsymbol{I}_L\right)\boldsymbol{Q}\right]^{-1} = \hat{\tau}^{-1}\boldsymbol{Q}^{-1}\left(\boldsymbol{S}^T\boldsymbol{S}+L\hat{\lambda}^{-1}\boldsymbol{I}_L\right)^{-1}\boldsymbol{Q}\\ &= \hat{\tau}^{-1}\boldsymbol{Q}^T\left(\boldsymbol{S}^T\boldsymbol{S}+L\hat{\lambda}^{-1}\boldsymbol{I}_L\right)^{-1}\boldsymbol{Q}\end{aligned} \tag{90}$$

On the other hand, based on the same equations of Kendall, S. Earle, C-H. Wu, et al have directly deduced the posterior distribution of $\boldsymbol{\delta}$ for the LMM (Equation 75) used in BugWAS under the null hypothesis where $\beta = 0$ [6]. Specifically, $\boldsymbol{\delta}|\boldsymbol{y}$ follows a multivariate normal distribution that takes as parameters the REML estimates of $\lambda'$ and $\tau$ (cf. source codes of the BugWAS package).

$$\boldsymbol{\delta} \sim MVN_L(\boldsymbol{\mu}_\delta, \boldsymbol{\Delta}_\delta) \tag{91}$$

$$\boldsymbol{\mu}_\delta = \left(\boldsymbol{S}^T\boldsymbol{S}+\frac{1}{\hat{\lambda}'}\boldsymbol{I}_L\right)^{-1}\boldsymbol{S}^T\boldsymbol{y} \tag{92}$$

$$\boldsymbol{\Delta}_\delta = \hat{\tau}^{-1}\left(\boldsymbol{S}^T\boldsymbol{S}+\frac{1}{\hat{\lambda}'}\boldsymbol{I}_L\right)^{-1} \tag{93}$$

Since $\boldsymbol{S} = \boldsymbol{P\Sigma Q}^T = \boldsymbol{C}_L\boldsymbol{Q}^T$, we have $\boldsymbol{S\delta} = \boldsymbol{C}_L\boldsymbol{Q}^T\boldsymbol{\delta}$. Let $\boldsymbol{\gamma}_\delta = \boldsymbol{Q}^T\boldsymbol{\delta}$ and $\hat{\lambda} = L\hat{\lambda}'$, we can obtain an affine transformation of the posterior multivariate normal distribution of the structural random effects $\boldsymbol{\gamma}_\delta$:

$$\boldsymbol{\gamma}_\delta \sim MVN_L\left(\boldsymbol{\mu}'_\delta, \boldsymbol{\Delta}'_\delta\right) \tag{94}$$

$$\boldsymbol{\mu}'_{\delta} = \boldsymbol{Q}^T \boldsymbol{\mu}_{\delta} = \boldsymbol{Q}^T \left( \boldsymbol{S}^T \boldsymbol{S} + \frac{1}{\hat{\lambda}'} \boldsymbol{I}_L \right)^{-1} \boldsymbol{S}^T \boldsymbol{y} = \boldsymbol{Q}^T \left( \boldsymbol{S}^T \boldsymbol{S} + L\hat{\lambda}^{-1} \boldsymbol{I}_L \right)^{-1} \boldsymbol{S}^T \boldsymbol{y} \tag{95}$$

$$\boldsymbol{\Delta}'_{\delta} = \boldsymbol{Q}^T \boldsymbol{\Delta}_{\delta} \boldsymbol{Q} = \hat{\tau}^{-1} \boldsymbol{Q}^T \left( \boldsymbol{S}^T \boldsymbol{S} + L\hat{\lambda}^{-1} \boldsymbol{I}_L \right)^{-1} \boldsymbol{Q} \tag{96}$$

Both $\boldsymbol{\mu}'_{\delta}$ and $\boldsymbol{\Delta}'_{\delta}$ are exactly the same as Equations 89 and 90. As such, the structural random effects $\boldsymbol{\gamma}$ in our model (Equation 25) and the $\boldsymbol{\delta}$ in the BugWAS model are interconnected in terms of their posterior multivariate normal distributions given observations $\boldsymbol{y}$ and $\boldsymbol{S}$.

**Posterior distribution of structural random effects computed using reduced SVD**
We can derive the posterior distribution of structural random effects given $\boldsymbol{y}$ through the reduced form of SVD as well. Using the reduced form of SVD, we have shown the equivalent form of our LMM in Formulae 31 and 32, where $\boldsymbol{S} = \boldsymbol{U}\boldsymbol{D}\boldsymbol{V}^T$ and $\boldsymbol{C}_r = \boldsymbol{U}\boldsymbol{D} = \boldsymbol{S}\boldsymbol{V}$. Since Formulae 25 and 31 only differ in the vector for sizes of structural random effects (that is, $\boldsymbol{\gamma}$ versus $\boldsymbol{\gamma}_r$), substituting $\boldsymbol{C}_L$ with $\boldsymbol{C}_r$ and $\boldsymbol{I}_L$ with $\boldsymbol{r}$ in Equation 58 immediately produces the posterior distribution:

$$\boldsymbol{\gamma}_r | \boldsymbol{y} \sim MVN_n\left(\boldsymbol{\mu}_r, \boldsymbol{\Delta}_r\right) \tag{97}$$

$$\boldsymbol{\mu}_r = \left( \boldsymbol{C}_r^T \boldsymbol{C}_r + L\hat{\lambda}^{-1} \boldsymbol{I}_r \right)^{-1} \boldsymbol{C}_r^T \boldsymbol{y} \tag{98}$$

$$\boldsymbol{\Delta}_r = \hat{\tau}^{-1} \left( L\hat{\lambda}^{-1} \boldsymbol{I}_r + \boldsymbol{C}_r^T \boldsymbol{C}_r \right)^{-1} \tag{99}$$

Moreover, since Equation 13 shows $\boldsymbol{Q} = \begin{bmatrix} \boldsymbol{V}_{L\times r} & \boldsymbol{W}_{L\times(L-r)} \end{bmatrix}$, we rewrite Equation 89 as

$$\begin{aligned} \boldsymbol{\mu} = \boldsymbol{Q}^T \left( L\hat{\lambda}^{-1} \boldsymbol{I}_L + \boldsymbol{S}^T \boldsymbol{S} \right)^{-1} \boldsymbol{S}^T \boldsymbol{y} &= \begin{bmatrix} \boldsymbol{V}^T \\ \boldsymbol{W}^T \end{bmatrix} \left( L\hat{\lambda}^{-1} \boldsymbol{I}_L + \boldsymbol{S}^T \boldsymbol{S} \right)^{-1} \boldsymbol{S}^T \boldsymbol{y} \\ &= \begin{bmatrix} \boldsymbol{V}^T \left( L\hat{\lambda}^{-1} \boldsymbol{I}_L + \boldsymbol{S}^T \boldsymbol{S} \right)^{-1} \boldsymbol{S}^T \boldsymbol{y} \\ \boldsymbol{W}^T \left( L\hat{\lambda}^{-1} \boldsymbol{I}_L + \boldsymbol{S}^T \boldsymbol{S} \right)^{-1} \boldsymbol{S}^T \boldsymbol{y} \end{bmatrix} \end{aligned} \tag{100}$$

The top partition, $\boldsymbol{V}^T \left( L\hat{\lambda}^{-1} \boldsymbol{I}_L + \boldsymbol{S}^T \boldsymbol{S} \right)^{-1} \boldsymbol{S}^T \boldsymbol{y}$, equals the posterior mean of $\boldsymbol{\gamma}_r$ given $\boldsymbol{y}$; the second partition, $\boldsymbol{W}^T \left( L\hat{\lambda}^{-1} \boldsymbol{I}_L + \boldsymbol{S}^T \boldsymbol{S} \right)^{-1} \boldsymbol{S}^T \boldsymbol{y}$, equals the posterior mean of $\boldsymbol{\gamma}_{L-r}$ that always get cancelled in our LMM (Equation 25) because $\boldsymbol{c}_j = 0$ when $j = r+1, r+2, \cdots, L$. Moreover, this partition does not involve in the LMM (Equation 31) and we are not interested in $\boldsymbol{\gamma}_{L-r}$ as the accompanying projections do not reveal any variance in $\boldsymbol{S}$ (namely, no genetic variation is captured by these effects).

Similarly, we can convert the posterior variance-covariance matrix (Equation 58) of $\boldsymbol{\gamma}$ into the following form:

$$\begin{aligned}\boldsymbol{\Sigma} &= \hat{\tau}^{-1}\boldsymbol{Q}^T\left(\boldsymbol{S}^T\boldsymbol{S}+L\hat{\lambda}^{-1}\boldsymbol{I}_L\right)^{-1}\boldsymbol{Q} = \hat{\tau}^{-1}\begin{bmatrix}\boldsymbol{V}^T\\ \boldsymbol{W}^T\end{bmatrix}\left(L\hat{\lambda}^{-1}\boldsymbol{I}_L+\boldsymbol{S}^T\boldsymbol{S}\right)^{-1}\begin{bmatrix}\boldsymbol{V}_{L\times r} & \boldsymbol{W}_{L\times(L-r)}\end{bmatrix}\\ &= \hat{\tau}^{-1}\begin{bmatrix}\boldsymbol{V}^T\left(\boldsymbol{S}^T\boldsymbol{S}+L\hat{\lambda}^{-1}\boldsymbol{I}_L\right)^{-1}\boldsymbol{V} & \boldsymbol{V}^T\left(\boldsymbol{S}^T\boldsymbol{S}+L\hat{\lambda}^{-1}\boldsymbol{I}_L\right)^{-1}\boldsymbol{W}\\ \boldsymbol{W}^T\left(\boldsymbol{S}^T\boldsymbol{S}+L\hat{\lambda}^{-1}\boldsymbol{I}_L\right)^{-1}\boldsymbol{V} & \boldsymbol{W}^T\left(\boldsymbol{S}^T\boldsymbol{S}+L\hat{\lambda}^{-1}\boldsymbol{I}_L\right)^{-1}\boldsymbol{W}\end{bmatrix}\end{aligned} \tag{101}$$

The top-left $r\times r$ partition, $\hat{\tau}^{-1}\boldsymbol{V}^T\left(\boldsymbol{S}^T\boldsymbol{S}+L\hat{\lambda}^{-1}\boldsymbol{I}_L\right)^{-1}\boldsymbol{V}$, reveals the posterior variance-covariance between elements of $\boldsymbol{\gamma}_r$, in which we are interested for Model 31. Therefore, we have the posterior distribution of $\boldsymbol{\gamma}_r$, that is, Equation 97, in the null model of Equation 31 given $\boldsymbol{y}$ and $\boldsymbol{K}$.

$$\boldsymbol{\mu}_r = \boldsymbol{V}^T\left(L\hat{\lambda}^{-1}\boldsymbol{I}_L+\boldsymbol{S}^T\boldsymbol{S}\right)^{-1}\boldsymbol{S}^T\boldsymbol{y} \tag{102}$$

$$\boldsymbol{\Delta}_r = \hat{\tau}^{-1}\boldsymbol{V}^T\left(\boldsymbol{S}^T\boldsymbol{S}+L\hat{\lambda}^{-1}\boldsymbol{I}_L\right)^{-1}\boldsymbol{V} \tag{103}$$

which is exactly the posterior distribution of structural random effects calculated by the BugWAS package[1]. We can also deduce this distribution directly from the BugWAS model with the reduced form of SVD. More specifically, since $\boldsymbol{S}=\boldsymbol{U}\boldsymbol{D}\boldsymbol{V}^T=\boldsymbol{C}_r\boldsymbol{V}^T$, we can deduce that $\boldsymbol{S}\boldsymbol{\delta}=\boldsymbol{C}_r\boldsymbol{V}^T\boldsymbol{\delta}$. Let $\boldsymbol{\gamma}'_\delta=\boldsymbol{V}^T\boldsymbol{\delta}$, knowing the posterior mean (Equation 92) and variance-covariance matrix (Equation 93) of $\boldsymbol{\delta}$, we can derive the posterior distribution of $\boldsymbol{\gamma}'_\delta$ given $\boldsymbol{y}$ through an affine transformation of the posterior multivariate normal distribution of $\boldsymbol{\delta}$.

$$\boldsymbol{\gamma}'_\delta|\boldsymbol{y} \sim MVN_L\left(\boldsymbol{\mu}'_\delta,\boldsymbol{\Delta}'_\delta\right) \tag{104}$$

$$\boldsymbol{\mu}'_\delta = \boldsymbol{V}^T\boldsymbol{\mu}_\delta = \boldsymbol{V}^T\left(\boldsymbol{S}^T\boldsymbol{S}+\frac{1}{\hat{\lambda}'}\boldsymbol{I}_L\right)^{-1}\boldsymbol{S}^T\boldsymbol{y} \tag{105}$$

$$\boldsymbol{\Delta}'_\delta = \boldsymbol{V}^T\boldsymbol{\Delta}_\delta\boldsymbol{V} = \hat{\tau}^{-1}\boldsymbol{V}^T\left(\boldsymbol{S}^T\boldsymbol{S}+\frac{1}{\hat{\lambda}'}\boldsymbol{I}_L\right)^{-1}\boldsymbol{V} \tag{106}$$

[1]Note that the Equation 103 in the BugWAS paper is different to the one implemented in the codes of BugWAS. Based on algebra shown in this section, we inclined to treat the equation in the paper as incorrect due to a typo.

Evidently, $\boldsymbol{\mu}'_\delta$ and $\boldsymbol{\Delta}'_\delta$ are identical to $\boldsymbol{\mu}_r$ and $\boldsymbol{\Delta}_r$ in Equation 97, proving that the equivalence between posterior distributions in our LMM (Equation 31) and the BugWAS model still holds under the reduced form of SVD. This is an ideal conclusion. Nevertheless, we note that authors of the BugWAS paper used the following inference to the calculate the posterior distribution of structural random effects:

$$\boldsymbol{C} = \boldsymbol{S}\boldsymbol{V}_n \Longrightarrow \boldsymbol{S} = \boldsymbol{C}\boldsymbol{V}_n^{-1} \Longrightarrow \boldsymbol{S}\boldsymbol{\delta} = \boldsymbol{C}\boldsymbol{V}_n^{-1}\boldsymbol{\delta} \Longrightarrow \boldsymbol{\gamma}_\delta = \boldsymbol{V}_n^{-1}\boldsymbol{\delta} \tag{107}$$

where $\boldsymbol{V}_n$ denotes the first $n$ columns of $\boldsymbol{Q}$ and it is computed using the R function *prcomp*. We consider this inference invalid because $\boldsymbol{V}$ is a rectangular matrix, to which an ordinary matrix inverse does not apply. Furthermore, we argue that only the first $r$ columns of $\boldsymbol{Q}$ (that is, the $L \times r$ matrix $\boldsymbol{V}$) should be used in this reduced form of SVD for efficiency because $r$ is always less than $n$ (as BugWAS also uses the zero-centring process) and the $n - r$ excessive eigenvectors in $\boldsymbol{Q}$ do not capture population structure and hence are not informative.

##### 3.4.3 Interpretations of the fixed effect size

Herein, we are concerning the meaning of the fixed effect parameter $\beta$ in our LMMs (Equations 25 and 31). Let dichotomous random variables $X$ and $Y$ denote the presence-absence of alleles $\mathrm{a_x}$ and $\mathrm{a_y}$, respectively, in a sample. In particular, we do not perform zero-centring on observations of $X$ or $Y$ in this section to make our description clearer while the centring does not affect our conclusions.

The first interpretation of the $\beta$ is the change in the conditional mean of $Y$, namely, $E(Y|X)$, given a unit change in $X$ under the constraints that $\Delta X = -1$ when $X = 1$ and $\Delta X = 1$ when $X = 0$ at the beginning. To be more specific, let $E(Y|X) = E(Y|X + \Delta X) - E(Y|X)$. Since for a genome, $E(Y|X) = E(\alpha + X\beta + u + \varepsilon)$, and random variables ($u$ and $\varepsilon$) are independent of X following the model setting, we have $\Delta E(Y|X) = \beta \Delta X$, which can be denoted by $\Delta E(Y|\Delta X)$. This is a general interpretation and it is the same as the meaning of regression coefficients in a simple linear model.

Secondly, when $\beta > 0$ in particular, this fixed effect becomes the probability of observing an allele $\mathrm{a_y}$ in a genome given an acquisition event of the other allele $\mathrm{a_x}$. Assuming there are $n$ genomes (where $n$ is sufficiently large) in total, which were void of alleles $\mathrm{a_x}$ and $\mathrm{a_y}$. Then $n_x$ and $n_y$ of these genomes acquired $\mathrm{a_x}$ and $\mathrm{a_y}$, respectively, via HGT. As such, the observation of $\Delta X$, denoted as $\Delta x$, increases to one in these $n_x$ genomes. Let $n_{xy}$ denote the number of genomes harbouring both alleles. Noticing $\Delta x = 0$ if a strain does not have the $\mathrm{a_x}$ allele, we derived that

$$\Delta E(Y|\Delta X = 1) \approx \bar{\boldsymbol{y}}|_{x=1} = \frac{n_{xy}}{n_x} = \beta \tag{108}$$

for the $n_x$ observations where $\Delta x = 1$ and $\boldsymbol{y}$ is a vector for observations of $Y$ in the $n$ genomes. Therefore, a positive $\beta$ can be understood as the probability of observing $\mathrm{a_y}$ when the allele $\mathrm{a_x}$ is successfully transferred into a bacterium.

More particularly, when X and Y do not show perfect separation as required by ordinary logistic regression [15], in other words, neither $P(Y = 1|X = 0)$ nor $P(Y = 1|X = 1)$ equals one or zero and hence $0 < E(Y|X = x) < 1$, a positive $\beta$ can also be interpreted as an approximation of an odds ratio through expanding a logistic model into a Maclaurin series. Specifically, let $f(x) = P(Y = 1|X = x)$, and let scalars $m$ and $b$ be the intercept and slope of the logistic model, respectively, then $0 < f(x) < 1$ and the Maclaurin expansion

$$f(x) = \frac{e^{m+xb}}{e^{m+xb}+1} = \frac{e^m}{e^m+1} + \frac{be^m}{(e^m+1)^2} + \mathrm{o}\left(x^2\right), x \to 0 \tag{109}$$

holds for $x$ close to zero when treating $x$ as a continuous variable. Furthermore, we can approximate $f(1) = P(Y = 1|X = 1)$ using this expansion. Since $Y$ follows a Bernoulli distribution conditioning on $X$, we have $E(Y|X = x) = P(Y = 1|X = x)$. Comparing to the conditional mean of $Y$ in our LMM for any genome, which is $0 < E(Y|X = x) = \alpha + x\beta < 1$ given $x \in \{0, 1\}$ and $\beta > 0$, we can deduce that $0 < \alpha < 1$ and

$$\alpha + x\beta \approx \frac{e^m}{e^m+1} + \frac{be^m}{(e^m+1)^2}x \tag{110}$$

Let $\alpha = \frac{e^m}{e^m+1}$ and $\beta = \frac{be^m}{(e^m+1)^2}$, we have $m = \ln\frac{\alpha}{1-\alpha}$ and $b = \frac{\beta(e^m+1)^2}{e^m} = \frac{\beta}{\alpha(1-\alpha)}$. As such, the odds ratio of $Y$ is approximated using the fixed effect sizes $\alpha$ and $\beta$ in the LMM via the equation

$$OR(Y|X = 1 \text{ vs. } X = 0) = \frac{odds(Y = 1 \text{ vs. } Y = 0|X = 1)}{odds(Y = 1 \text{ vs. } Y = 0|X = 0)} = e^b = \exp\frac{\beta}{\alpha(1-\alpha)} \tag{111}$$

We use GEMMA to perform a Wald test to determine if $\beta = 0$ under a certain confidence level (Section 3.1.10).

##### 3.4.4 Incapability of parameter estimation between identical variables

A crucial constraint for GEMMA in estimating parameters of LMMs is that parameter estimation does not work between identical variables. Let $\boldsymbol{y} = \boldsymbol{x}$, then Model 25 becomes $\boldsymbol{0} = \boldsymbol{1}\alpha + \boldsymbol{x}\beta' + \boldsymbol{C}_L\boldsymbol{\gamma} + \boldsymbol{\varepsilon}$, where $\beta' = \beta - 1$. As such, the target function $l_{r1}(\lambda; \boldsymbol{0}, \boldsymbol{x}, \boldsymbol{K})$ (cf. Formula 38) does not exist because one of its items ($\boldsymbol{y}^T\boldsymbol{W}_x\boldsymbol{y}$) for logarithm becomes zero. Therefore, GEMMA does not return a valid log-likelihood or REML estimates for parameters of variance components under this condition. We confirmed that this behaviour is the same for ordinary maximum-likelihood (ML) estimates for a similar reason (the denominator in the ML estimate of $\tau$ reaches zero – cf. Section 3.1.1 in the supplementary document of the GEMMA paper [4]). As a result, our approach is not applicable to identically distributed alleles or genes.

### 4 Supplementary methods of the validation study

This section demonstrates steps of applying GeneMates to the *E. coli* and *Salmonella* data sets for validation [16, 17].

#### 4.1 Collection of whole-genome sequencing data

Supplementary file 2 tabulates details of bacterial genomes whose WGS data were recruited for our validation study. For *E. coli* data, we used paired-end 100 bp reads (Illumina HiSeq2000) from 185 atypical enteropathogenic *E. coli* genomes from Africa and South Asia between 2008 and 2010 during the Global Enteric Multicentre Study (GEMS) [18, 19]. Most genomes were obtained from faecal samples. To ensure high read quality and purity, we inspected base and read quality using MultiQC [20], which compiled reports of FastQC v0.11.5 [21]. All read sets were accepted at this stage. We merged overlapping reads with FLASH [22] because a high level of overlapping reads was detected.

As for *Salmonella* data, we took paired-end Illumina reads (length: 76–150 bp) from 373 genomes of *S. enterica* serovar Typhimurium definitive type 104 (DT104), which were sampled between 1990 and 2011 [17]. The reads were generated using Illumina Genome Analyzer II or IIx, HiSeq2000 and MiSeq platforms. According to the original publication [17], we removed 14 known atypical DT104 genomes (all of them were isolated in Scotland) as they are genetically more similar to non-DT104. The remaining genomes included 275 Scottish genomes, 23 English or Welsh genomes, 51 Canadian genomes and 10 Japanese genomes. Finally we carried out the same steps for quality control as those for the *E. coli* data except the use of FLASH, because the *Salmonella* reads showed a low level of overlapping.

#### 4.2 Extraction of core-genome SNPs

We mapped de-overlapped *E. coli* reads to the chromosome sequence of *E. coli* serovar 0103:H2 strain 12009 (GenBank association: AP010958) with RedDog version V1beta.10.3 under its default arguments. This reference genome had been previously shown to yield the highest read coverage for this data set [19]. To obtain reliable SNP calls, we identified repetitive and prophage regions in the reference genome using MUMmer v3.23 ($\geq 90\%$ nucleotide identity to call repetitive) and PHASTER (accessed on 26 November 2017), respectively [23, 24], and filtered out SNPs within those regions. We applied the same mapping process to *Salmonella* reads, for which the chromosome sequence of DT104 (GenBank association: HF937208) was used as the reference. Instead of using PHASTER, which was under maintenance at the time (22 March 2018), PHAST [25] was used to identify prophages.

A challenge in the analysis of HGcoT is the requirement for homogenous read sets. A DNA library for sequencing may be contaminated due to mixed bacterial culture or artefacts in the library preparation. To evaluate heterogeneity in a read set, we compared coordinates and minor allele frequencies (MAFs) of heterozygous SNP calls to those of homozygous SNPs, with the expectation that bacterial cells are haploid. Any read set showing congruent MAFs of the heterozygous SNPs across the whole reference genome was considered as potentially contaminated. Altogether, 16 *E. coli* genomes and one *Salmonella* strain were excluded due to contamination, leaving 169 *E. coli* genomes and 359 *Salmonella* genomes for the following steps.

#### 4.3 Phylogenetic reconstruction

For each bacterial species, five independent RAxML (v8.2.9) runs were launched on homozygous SNPs in sites that were present in 99% of genomes to obtain an optimal unrooted maximum-likelihood phylogeny [26]. A unique combination of seeds was set for the random-number generator of RAxML in each run. To calculate bootstrap supports for branches on each phylogenetic tree, in each run we specified RAxML to repeat the tree inference for 125 iterations for *E. coli* genomes and 200 iterations for *Salmonella* genomes. The tree displaying the greatest log-likelihood was chosen as the best tree for further analyses.

#### 4.4 Detection of antimicrobial resistance genes

Based on a curated ARG-ANNOT database (ARGannot_r2.fasta, available in the GitHub repository of SRST2) [27], we screened AMR genes in all genomes using SRST2 v0.2.0 (arguments determining presence of an ARG: $> 90\%$ query coverage, $\geq 90\%$ nucleotide identity, $> 5$ fold average read depth and $> 2$ fold edge depths) [28]. We evaluated reliability of ARG calls based on per-base read depths calculated by SRST2 (Section 4.4.1). Next, for each bacterial species, we pooled consensus sequences of reliable gene calls in all genomes and clustered them under a nucleotide identity of 100% using CD-HIT-EST v4.6 [29]. An identifier was then assigned to each unique sequence to represents an allele of an AMR gene.

Since there was a complete chromosome sequence and plasmid sequences included for the *Salmonella* strain DT104, we could achieve high resolution and accuracy in gene screen via directly aligning reference allele sequences from ARGannot_r2.fasta against complete genomes using nucleotide BLAST [30]. Therefore, geneDetector (github.com/wanyuac/geneDetector) was developed to convert BLAST results into SRST2 compatible formats. Consensus allele sequences extracted with geneDetector were pooled with those from SRST2 for sequence clustering and subsequent allele identifier assignment, producing an allelic PAM from reliable allele calls.

##### 4.4.1 Evaluation of allele-call reliability

We used PAMmaker (github.com/wanyuac/PAMmaker), a helper tool of GeneMates, to evaluate reliability of allele calls. Specifically, PAMmaker read score files in SRST2 outputs and considered an allele call sufficiently reliable if it met the following criteria:

- When no truncation (at one or both ends of a reference) or deletion (flanked by remaining bases) of at least two bases was present in the alignment of reads against a reference allele sequence, the average read depth $> 5$ fold, edge read depths $\geq 2$ fold, and the nucleotide divergence $\leq 10\%$.
- When any truncations were present, nucleotide divergence $\leq 10\%$, the average read depth $> 5$ and read depths that neighbour truncations $> 2$.

#### 4.5 *De novo* genome assembly

*De novo* assembly of *E. coli* and *Salmonella* genomes was conducted using Unicycler v0.4.1 [31], which implements SPAdes v3.10.1 for short-read assembly [32] and leverages paired-end reads to optimise initial assembly graphs from SPAdes via bridging gaps, merging nodes and correcting assembly errors. In this study, we specified Unicycler to turn the SPAdes option of read correction off as this process turned out to be too conservative for some read sets to get the assembly pipeline successfully run through. Other options were kept default (see the manual of Unicycler for details). Finally, we gathered assembly graphs from Unicycler for measuring APDs and concatenated assembly statistics for further quality assessment of read sets.

#### 4.6 Measurement of allelic physical distances

Physical distances between alleles of AMR genes were measured in genome assemblies for each bacterial species. Since the distance measurement was complicated in unfinished short-read assemblies, we took an empirical simulation-and-validation strategy to determine criteria for removing unreliable measurements. The overall procedure was that, for each species, we simulated Illumina reads from complete genomes of 10 MDR strains that were drawn from non-sibling lineages in a published species dendrogram

(NCBI genome database, accessed in November 2017), reassembled the synthetic reads into assembly graphs using the same method as real reads, and for each pair of coding sequences (CDS), we compared APDs from assembly graphs to their real distances. See Tables s1 and s2 for details of selected *E. coli* and *S.* Typhimurium strains. We used FastANI v1.0 to calculate whole-genome ANIs between these strains.

For *E. coli* genomes, we used ART (version MountRainier) and its build-in Illumina HiSeq2000 error profile to simulate 100-bp pair-end reads at a read depth of 89 fold (the median depth covering the 169 *E. coli* genomes according to the RedDog outputs), a mean insert size of 220 bp with a standard deviation of 71 bp (both arguments are medians for this data set) [33]. For *Salmonella* genomes, we simulated short reads from the Illumina HiSeq2000 platform under a 76 bp read length, a 251 bp insert size with a 76 bp standard deviation and a 75 fold read depth in accordance with summary statistics of actual read sets of the shortest read length (76 bp). To ensure sufficient specificity in locating CDS in every assembly for the distance measurement, a hit of each CDS was reported by the nucleotide BLAST to Bandage only if it covered at least 95% of the CDS under a nucleotide identity of at least 95% and displayed an e-value of at most $1 \times 10^{-5}$. These parameters were kept the same for subsequent applications. Further, in order to compare the distance measurements and the reference distances in an efficient manner, we randomly selected 250k measurements per strain from the distances measured in paths of at most 10 nodes in the assembly graphs.

For the 169 *E. coli* genomes and 359 *Salmonella* genomes, we took consensus allele sequences of accessory AMR genes as queries, took assembly graphs and contigs as subjects, and used Bandage to measure the shortest-path distances (SPDs) in genome assemblies of each species. It is self-evident that, for the same pair of alleles, their SPD equals the APD in a complete genome and contig. Since in this empirical study, we found that SPDs in contigs were more accurate than those in assembly graphs, we prioritised SPDs according to their sources and kept the SPD from contigs when there were SPDs measurable in both a contig and a graph. Furthermore, when calling the function *findPhysLink*, we specified the function to filter out any measurements obtained from more than two nodes or stretched longer than 250 kbp to ensure that more than 90% of remaining distances were accurate when tolerating errors not exceeding $\pm 1$ kbp. Note that since Unicycler may create a contig from multiple nodes in an assembly graph, SPDs in a single node may differ from those in a single contig in accuracy.

#### 4.7 Network analysis

**Network construction** We used the function *findPhysLink* to construct a linkage network for alleles of AMR genes detected in each example data set. In particular, we excluded alleles of known intrinsic AMR genes (*ampH*, *ampC1*, *ampC2* and *mrdA* of

*E. coli*, and *aac6-Iaa* of *S. enterica*). Following the validation method for BugWAS [6], we did not filter alleles for a minimum frequency or co-occurrence count in order to include all possible allele pairs. In order to investigate effects of controlling for population structure on estimates of fixed effects, we fitted simple penalised logistic models (PLMs) (implemented in GeneMates function *plr*) for the same allele pairs. A significant fixed effect of an explanatory allele on a response allele was determined when the hypothesis test on the effect size $\beta$ in an LMM (Wald test) or a PLM (two-sided chi-squared test) returned a Bonferroni-corrected p-value $\leq 0.05$. We compared LMM-based p-values to those based on PLMs for the same explanatory and response alleles, and grouped the differences based on the estimate $\hat{\lambda}_0$ for the null LMM ($Y \sim 1$) of each response allele because $\lambda_0$ reveals the proportion of variation in $Y$ explained (PVE) by population structure in the absence of any explanatory allele [4].

**Visualisation** Result tables produced by *findPhysLink* for each species were exported to Cytoscape v3.6.1 for network visualisation. In every network, each node represented one or more alleles sharing the same distribution among genomes (We confirmed that the distance score $s_d = 1$ for edges between each of these identically distributed alleles to its neighbour nodes) and each edge represented a significant fixed effect ($\hat{\beta}$ in an LMM or PLM) of an explanatory allele X on a response allele Y in the linear model $Y \sim X$. Edge attributes included the fixed effect size $\hat{\beta}$ ($-1 \leq \hat{\beta} \leq 1$) and a distance score $s_d$ ($0 \leq s_d \leq 1$).

**Analyses** LMM-based linkage networks were analysed for three purposes. First, in order to validate the ability of LMMs in identifying co-mobilised alleles of AMR genes, we compared edges to known mobile ARG clusters – plasmid-borne AMR genes in *E. coli* [16]. Second, in order to show the effect of controlling for population structure in association tests, we merged LMM-based and PLM-based linkage networks for each species into a single network highlighting shared and unique edges in each kind of linkage networks. Finally, we identified maximal cliques in each linkage network using the package igraph, extracted corresponding nucleotide sequences from genome assemblies and searched them against GenBank (www.ncbi.nlm.nih.gov/genbank) to discover co-mobilised alleles of AMR genes.
